# Triangulation reduces the polygon of error for the history of Transeurasian

**DOI:** 10.1101/2022.10.05.510045

**Authors:** Martine Robbeets, Mark Hudson, Chao Ning, Remco Bouckaert, Alexander Savelyev, Geonyoung Kim, Tao Li, Sofia Oskolskaya, Ilya Gruntov, Olga Mazo, Seongha Rhee, Kyou-Dong Ahn, Ricardo Fernandes, Ken-ichi Shinoda, Hideaki Kanzawa-Kiriyama, Rasmus Bjørn, Bingcong Deng, Deog-im An, John Bentley, Takamune Kawashima, Joanna Dolińska

## Abstract

In a recent study we used an interdisciplinary approach combining linguistics, archaeology and genetics to analyse the Transeurasian languages^1^. Our analysis concluded that the early dispersals of these languages were driven by agriculture. A preprint published on this server presents objections to the Transeurasian hypothesis and its association with farming dispersals^2^. However, close inspection of that text reveals numerous misinterpretations and inconsistencies. In the interest of furthering scientific debate over Transeurasian language and population history, we address the critiques, revising datasets and fine-tuning approaches. The linguistic critique questions the quantity and quality of our datasets. Here we show that the number of surviving cognate sets for Transeurasian is in line with that for well-established language families. In addition, we find that Tian et al.’s failure to reject a core of regularly corresponding cognates in the basic vocabulary creates ground for a consensus about the genealogical relatability of the Transeurasian languages. The archaeological critique attempts a re-analysis of one Bayesian test using re-scored data only for northern China. Over half of the suggested re-scorings contain inconsistencies and it is not explained why the re-analysis retains the original data for sites outside northern China, comprising almost 60% of the total. More importantly, the sweeping claim that there is no evidence supporting the prehistoric migrations analysed in our study is not backed by any discussion of the archaeological record. With respect to genetics, the preprint claims a re-analysis showing that the data ‘do not conclusively support the farming-driven dispersal of Turkic, Mongolian, and Tungusic, nor the two-wave spread of farming to Korea.’ In fact, the only genetic re-analysis presented is limited to samples from Korea and Japan and does not contradict our original conservative modelling of Neolithic individuals with Hongshan and our Bronze Age ones with Upper Xiajiadian. In sum, in bringing multiple lines of evidence together through triangulation, we gained a more balanced and richer understanding of Transeurasian dispersals than each discipline could provide individually. Our research doubtless leaves room for improvement but we remain confident that triangulation did not ‘fail’, but rather brought us a step closer to understanding the history of the Transeurasian languages.

## Linguistics

Our Article used the classical historical comparative linguistic method to propose the genealogical relatedness of the Transeurasian languages. We then combined Bayesian phylogenetics with classical approaches to draw inferences about tree branching, dating and homelands. The critique that ‘the number of cognate sets proposed for Transeurasian is exceptionally low’ is based on comparison of our cognate set distributions with those of long-established language families, such as Sino-Tibetan and Indo-European. However, the authors ignored the full body of evidence in our traditional linguistic approach indicated in red in Table 1 and counted only the portion of semantically equivalent cognates used in our Bayesian analysis. In fact, the distribution of cognates is completely in line with expectations, considering that Transeurasian has an older age, fewer surviving daughter languages and early attestations of a relatively shallow time-depth, so that traces of cognancy are more blurred than in the other families.

**Table 1:**
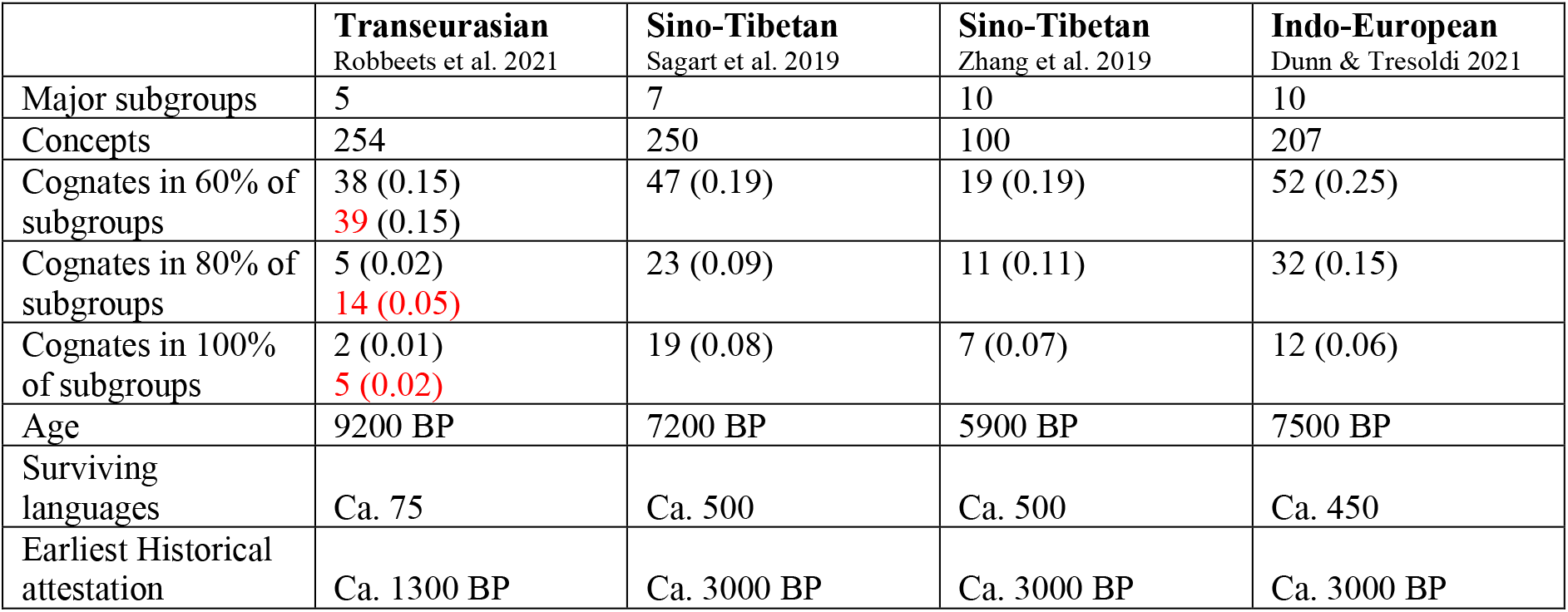
Distribution of cognates across a collection of datasets. Adapted from [^2^].

Furthermore, the criticism that ‘Out of the 3139 cognate sets listed, only 317 are found in more than one language family’ visualised in their Figure 1a is misleading because it is common in Bayesian cognate alignments for the bulk of cognates to be singletons; Sino-Tibetan and Indo-European would yield a similar graphic.

The authors claim we ‘ignore [our] own principles of historical language comparison and bend the evidence to [our] needs.’ However, in line with the requirements of the historical comparative method, all compared sounds correspond regularly, except for some root-final vowels (see SI for correspondences in each cognate set). Our item-by-item treatment of Tian et al.’s alleged ‘irregularities in the sound correspondences’ reveals that only 2 basic etymologies involve flaws, while other comments are open to interpretation.

Moreover, it is generally known^3^ that computer-assisted cognate detection methods, such as the one applied by the authors, still perform worse than expert analysis and that the algorithm fails when the datasets are too small. Therefore, it is misleading to apply these methods to only a small fraction of our evidence (46 out of 160 etymologies; see SI).

As with any work of massive scope, differences of interpretation are inevitable. Pokorny’s corpus of Indo-European data has been continuously reworked since the 1920s^4,5,6^. Nevertheless, the availability of a publicly available dataset that can be further improved aims to push the field forward.

## Archaeology

The archaeological critique focuses on one quantitative Bayesian test, ignoring our qualitative analyses which provide key support for the overall conclusions. Tian et al. attach a file which re-scores ‘missing data’ for selected categories. However, examination of their data shows inconsistencies for over half of the suggested re-scorings (SI). Moreover, data are re-scored only for northern China and not for all regions in our database. These issues lend little confidence to the new tree found in the Comment.

Although the authors justly point out that ‘different subsets of the data have different histories’, taken together, the entire body of cultural features can nevertheless contribute to an overall history.^7^ Their suggestion of inferring trees inside trees is appealing but computationally not yet feasible at this time.^8^

The second issue about the appropriateness of phylogenetic trees to represent archaeological data challenges research — including that by a member of Tian’s team — finding that certain levels of horizontal transmission do not invalidate phylogenetic approaches to cultural evolution.^7,9,10^ Recognizing that the phylogenetic analysis of our archaeological dataset does not only capture a signal of descent but also one of interaction, our Article used clustering in the Bayesian tree as an indication of cultural similarity between different archaeological assemblages. This result was broadly consistent with our qualitative analysis and with the large body of archaeological literature which models agricultural expansions from northeast China into Manchuria^11,12^, the Primorye^13,14,15,16,17^, Korea^18,19,20^, and Japan^21,22,23,24^. While debate has revolved around scale, timing and routes, these spreads themselves are widely accepted. Tian et al.’s sweeping claim that there is no archaeological evidence supporting these migrations goes completely against the existing literature, yet no archaeological evidence is provided in support.

## Genetics

Against our suggestion that the first farmers in the West Liao River area were of Amur-like genetic profile, Tian et al. object that there was long-term genetic continuity of Amur-related hunter-gatherer ancestry in Northeast Asia, which predated agriculture. However, our Article does not question this continuity but instead shows that while the genetic profile of Early Neolithic millet farmers in the West Liao Basin was originally Amur-like, it received gradual gene flow from the Yellow River Basin. The authors assert that ‘[t]his conjecture is contradicted by their claim that both the spread of Japonic and that of Koreanic were induced by West Liao River farmer ancestry with Yellow River farmer admixture.’ However, there is no contradiction because the originally Amur-like farmers in the West Liao Basin had become admixed with 60 % Yellow River-related ancestry by 5500 BP, the time associated with the dispersal of millet agriculture to Korea. That explains why Neolithic Koreans and Bronze Age Japanese can be modelled as an admixture of Amur, Yellow River and Jomon genomes, a finding confirmed in new research.^25^

Finally, the authors object to our distinction between Hongshan and Upper Xiajiadian as a genetic model for our ancient Korean and Japanese populations. However, both our Article (SI 13) and extended analyses (SI: Fig. 1-6) confirm that the precise mainland East Asian ancestry source for ancient Koreans and Japanese cannot be well distinguished by current genetic data. If models cannot distinguish which population provides the best ancestry source for the target populations, in population genetics the principle is to select a population which is close to the target population both in time and space. We therefore modelled our Neolithic individuals with Hongshan and our Bronze Age ones with Upper Xiajiadian. Even if our data lack resolution to identify the precise source, they still allow us to associate the spread of farming to Korea with different waves of eastward gene flow from mainland East Asia. The increasing Yellow river component in our Bronze Age samples can be associated with rice farming, an observation supported by archaeological and linguistic findings (see SI).

## Triangulation

Tian et al. insist that our methods ‘fail to replicate’ but concentrate on replicating a single Bayesian test, ignoring the multiple other approaches in our paper. Replication of a single test may lead to repeating the same methodological error and thus provide an unfounded sense of certainty.^26^ We believe that an essential protection against such flaws is triangulation. In contrast to the authors’ claim that we ‘neither define nor describe the method of “ triangulation” ‘, the concept is well described in the methodology section and illustrated in the supplementary information of our Article. Triangulation is the strategic use of multiple disciplines, methods and datasets to address one question in order to enhance the validity and credibility of research findings. Bringing the multiple lines of evidence together, we gained a more balanced and richer understanding of Transeurasian dispersals than each of them could provide individually. Our research doubtless leaves room for improvement but we remain confident that triangulation did not ‘fail’.

## Supporting information

Supplementary Info 1: Reanalyses

Supplementary Info 3: Item-by-item analysis

