## Supplementary Info 1: Reanalyses for "Triangulation reduces the polygon of error for the history of Transeurasian"

**Contents**

1 Linguistic reanalysis.

1.1. Basic vocabulary comparative sets

1.2. Comparative sets relating to subsistence

1.3. Regular sound correspondences

1.3.1. Expert analysis

1.3.2. Automatically computed judgments

1.3.3. Suprasegmental correspondences

1.4. Number of cognates

1.5. Sources

1.5.1. Japonic sources

1.5.2. Koreanic sources

1.5.3. Tungusic sources

1.5.4. Mongolic sources

1.5.5. Turkic sources

2 Archaeological reanalysis

2.1. Missing data

2.2. Archaeological phylogeny

2.3. Population movements

3 Genetic reanalysis

4 Methodological revisit

4.1. Overall conclusions

4.2. Triangulation

4.3. Recapitulation

**Abbreviations**

| | |
|---|---|
| J | Japanese |
| K | Korean |
| Ma. | Manchu |
| MK | Middle Korean |
| MMo. | Middle Mongolian |
| MT | Middle Turkic |
| Na. | Nanai |
| OJ | Old Japanese |
| OT | Old Turkic |
| pA | proto-Altaic |
| pJ | proto-Japonic |
| pK | proto-Koreanic |
| pMo | proto-Mongolic |
| pTEA | proto-Transeurasian |
| pTg | proto-Tungusic |
| pTk | proto-Turkic |
| WMo. | Written Mongolian |

### 1 Linguistic reanalysis.

Tian et al. launch their comment to our Article by describing the Transeurasian linguistic unity with tendentious language such as “highly disputed”, “always been controversial” and “still remains a minority view”. While these descriptions seem designed to encourage the reader to doubt the validity of the Article in question even before substantive critiques are presented, the existence of controversy surrounding a scientific hypothesis is not a reason for *a priori* dismissal. Below we examine their linguistic comments in more detail and show that their objections contain numerous misinterpretations.

#### 1.1. Basic vocabulary comparative sets

Tian et al. object that “Robbeets et al.’s analysis of the linguistic data does not conform to the minimal standards required by traditional scholarship in historical linguistics and contradicts their own stated sound correspondence principles”.

The traditional method applied for establishing a genealogical relationship between two or more languages is the historical-comparative method. It is a procedure for reconstructing an unattested ancestral state of a language based on the detection of regular sound correspondences among sets of cognates in related languages, available from a later period. The ancestral language reconstructed in this way is referred to as the "proto-language", i.e. the common source of all the languages in a given family. The systematicity by which the sounds of a language change over time allows linguists to reconstruct the sounds and words of the proto-language. In the comparison of languages, it is standard to give particular attention to ‘basic vocabulary’ items. These are concepts that are available across the languages of the world, relatively independent of cultural context and tend to resist borrowing more successfully than random lexical items.

In line with standard methodology, we applied the historical-comparative method to the basic vocabulary of the Transeurasian languages by identifying regular sound correspondences in each correspondence set. Table 1 summarizes the basic vocabulary comparative sets in support of Transeurasian affinity advanced in our Article (SI 2). All forms correspond regularly for —if applicable— a subsequent initial consonant, medial vowel and medial consonant, but in several cases the occasional root-final vowel corresponds regularly as well. In the penultimate column of Table 1, each sound correspondence is numbered in accordance with the list of consonant and vowel correspondences provided in our Article and summarized in Section 1.3.1 of this supplement.

The final column of Table 1 summarizes our item-by-item reply to the "detailed qualitative analysis" by Tian et al (SI: 5-33). For reasons of transparency and to avoid recapitulation of the same datasets, we inserted our full replies in the pdf comment function of the supplementary information by Tian et al. Grey shading marks members of etymologies that are commented on by Tian et al. but do not lead to the elimination of the proto-form in question. Red shading marks members of etymologies that in line with the authors' suggestion need to be removed from the body of evidence. White cells contain evidence that remains uncommented on by Tian et al.

In addition, there are several proposals by Tian et al. to add "missing cognates" (SI: 5-33). Since adding cognates strengthens a hypothesis of linguistic affiliation, rather than weakening it, we gratefully accepted these suggestions and added them to the growing body of evidence.

Our item-by-item treatment of Tian et al.'s alleged "irregularities in the sound correspondences" reveals that the majority of the members of the basic etymologies remain uncriticized. This silent recognition of a core of regularly corresponding cognates in the basic vocabulary creates ground for a consensus about the genealogical relatability of the languages concerned. Only 2 members of the basic etymologies are marked in red because they indeed represent irregularities necessitating their removal from the evidence. The remaining comments, marked in grey, are open to interpretation; some issues were explained in detail in our Article (SI 2) but were overlooked by the authors, others suggest an alternative explanation of a certain datapoint and yet others are non-crucial for the present purpose but offer incentives for future research.

With regard to criticism of consistency of data collection, we would like to point out that linguistic data sets are never as constant as we would like them to be: Pokorny's monumental Indo-European etymological dictionary has recently been intensively reworked[1,2,3], encouraging one member of Tian's team to clean up the IELex dataset, currently available for Bayesian analyses of Indo-European[4]. However, that does not seem to be a reason for Tian et al. to disregard all previous research based on this dataset or even for advancing it as the

[1] Walde, A. & Pokorny, J. *Vergleichendes Wörterbuch der indogermanischen Sprachen*, 3 vols. (De Gruyter, 1927–1932).
[2] Pokorny, J. *Indogermanisches etymologisches Wörterbuch,* 2 vols. (Francke, 1948–69)
[3] Lubotsky, A. (ed.) *Leiden Indo-European Etymological Dictionary Series*, 12 vols. (Brill, 1991–2021).
[4] Dunn, Michael & Tiago Tresoldi. 2021. evotext/ielex-data-and-tree: IELex data and tree (2021/11/08).Version r20211108. Zenodo. https//doi.org/10.521/Zenodo.555601.

standard for Indo-European (Tian et al. SI: 5). Like in any work of massive scope, differences of interpretation are inevitable for the Transeurasian corpus but having a publicly available dataset that can be further improved will certainly push the field forward. The detail of Tian et al.'s Supplementary Information is only possible because our Article made all data and code available, stated assumptions and ensured full transparency. Acknowledging this instead of claiming to uphold the standards would have been more constructive.

**Table 1** Summary of the basic vocabulary comparative sets in support of Transeurasian affinity with indication of criticized members

| No | LJ item | Proto-Japonic | Proto-Koreanic | Proto-Tungusic | Proto-Mongolic | Proto-Turkic | Corr. no | "Comment" and short reply |
|---|---|---|---|---|---|---|---|---|
| 01 | 1 fire | *pɨ(r)i 'fire' | *pɨl 'fire' | | | | 1, 37, 31 | |
| 02 | 3 to go | *na- 'go away' | *na- 'go out' | *-na:- 'go out to' | | | 27, 32 | |
| 03 | 4 water | | *mɨl 'water' | *mö: ~ mu: 'water' | *mören 'river' | | 25, 37, 29 | |
| 04 | 5 mout h | *kutɨ-i 'mouth, opening' | *kut 'hollow, pit, cave' | | | | 13, 38, 8 | |
| 05 | | | | *amga < *?ama-g 'mouth' | *ama-n 'mouth, opening' | | 41, 26, 41 | |
| 06 | | *ipa- 'tell' | *ip 'mouth' | | | | 46, 2 | |
| 07 | 7 blood | *ti 'blood' | | | *či < *ti 'blood' | *tï:n 'spirit, breath' | 7, 40 | |
| 08 | 8 bone | *pəni(C)a 'bone' | *peCi 'bone' | *peni- 'knee' | | | 1, 34, 28, 40 | |
| 09 | 9 2sg pronoun | *na '2 sg pronoun' | *ne '2 sg pronoun' | | | | 27, 33 | |
| 10 | | | | *si '2 sg pronoun' | | *si '2 sg pronoun' | 23, 40 | |
| 11 | 11 to come | *kə- 'come' | | | | *kel- 'come' | 13, 34 | |
| 12 | 12 brea st | *kɨkɨ-rə 'heart' | | *xökö-n 'breast' | *kökö-n 'breast' | *kökü-r$_2$ 'breast' | 21, 37, 14 | "two different words at the Tungusic level." If so, at least the one mentioned here is corresponding; see SI2: 8 for explanation |

| | | | | | | | | |
|---|---|---|---|---|---|---|---|---|
| 13 | 14 1sg pronoun | *wa-n- ‘1sg/pl pronoun' | | | *ba-n- ‘1pl excl. pronoun’ | | 3, 32 | |
| 14 | | | | *bi '1 sg pronoun' | *bi '1 sg pronoun' | *bi '1 sg pronoun' | 3, 40 | “Nugteren reconstructs pMo back vowel *bï?” Disagreement in literature about reconstruction of *ï in pMo. Arguments contra by Svantesson[5] (2005:117-118). Arguments pro by Nugteren[27] but he (2011: 281) nevertheless reconstructs pMo *bi < *pre-pMo *bï “Only -i- forms and Chu. go back to pTk *bi- and other forms go back to *be-“ This is a residue of vowel alternation for singulars and plurals in the Transeurasian pronominal system, which became blurred due to the augmentation with plural suffixes. |
| 15 | 16 louse | | | | *sir-ke 'louse' | *sir-ke 'nit' | 23,40, 18,33 | |
| 16 | 19 arm/ hand | *ta(r)i ‘upper limb, arm, hand’ | *tali ‘lower limb, leg’ | | | | 19, 32, 31, 40 | |
| 17 | | *sune ‘lower limb, leg’ | *son ‘upper limb, arm, hand’ | | | | 23, 36, 28 | |
| 18 | | | | | *gar 'hand, arm' | *karï 'arm' | 15,32, 29 | |
| 19 | 22 ear | | | | *kul ‘ear’ | *kul-kak ‘ear’ | 13, 39, 31 | |
| 20 | 24 far | *mara ‘rare, from afar’ | *melu- ‘be far’ | | | | 25, 33, 31 | |
| 21 | 25 to do/make | *-ka- iconic | *-ki- iconic | *-ki- iconic | *ki- ‘do, make’ | *kïl- ‘do, make’ | 13, 40 | “Yng. khírun does not belong here” OK, but no impact on pJ reconstruction. “No account is provided for the coda *-l in pTk “ SI 2:7 gives an account |
| 22 | 26 hous e | *(y)ipi ‘house, hut’ | *cip(i) ‘house' | *ji:b 'house' | | | 9, 47, 4, 47 | “*y- is not supported by internal evidence” pJ *yi- is supported by internal evidence, only the distinction between *yi- and *i- is not, see SI 2: 12 for explanation “pK *cipi ‘house’” OK, works as well. |

5 Svantesson, J.-O., Tsendina, A., Mukhanova Karlsson, A. & Franzén, F. 2005. *The Phonology of Mongolian*. (Oxford, 2005)

| | | | | | | | |
|---|---|---|---|---|---|---|---|
| 23 | 27 stone/ rock | | | *kada:-r 'rock, cliff' | *kada 'rock, cliff' | | 13, 32 10, 32 |
| 24 | 30 tooth | *pa ‘tooth’ | *pal ‘tooth’ | | | | 1, 32 |
| 25 | | | | | *ari-ga 'molar, fang' | *ar$_2$(-)ïg 'molar, fang' | 41,30 |
| 26 | | | | | *sil-sün ‘tooth' | *siš ‘tooth’ | 23, 40 20b |
| 27 | 31 hair | *kama ~ kami ‘hair of the head’ | *kama ‘whirl of hair on the head’ | | | | 13, 32, 26 |
| 28 | | *kara ~ ka(r)i 'hair' | *kal 'hair' | | | | 13, 32, 29 |
| 29 | | | | | *kilga 'coarse hair' | *kïl(k) 'hair' | 13, 40, 31 |
| 30 | 32 big | *kɨki- ‘much' | *kɨki- ‘big’ | | | *kök ‘big, thick, healthy’ | 13, 37 14, 37 |
| 31 | | *mana- 'be big, many' | | *mani 'crowd' | *man 'big, high' | *bạnɨ 'big' | 25, 32, 28 |
| 32 | | | | *amban 'big' | *amban 'big' | | 41, 6, 32, 28 |
| 33 | 33 one | | *pilɨ- ~ *pilʌ- ‘to begin’ | | | *bir ‘one’ | 3, 40, 31 |
| 34 | 36 to hit/beat | *tuk- ‘hit with force’ | *t(ʌ)ki- ‘hit, strike’ | *dug- ‘hit’ | | | 9, 39, 16 |
| 35 | 37 leg /foot | *panki ‘lower leg, foot’ | *pal ‘foot, leg’ pK *pʌlk ‘arm’ | *palgan ‘foot’ | | | 1, 32, 18 |
| 36 | 39 this | *i ‘you’ (derogatory 2sg) | *i ‘this’ (proximal demonstrative) | *i ‘he, she, it’ (3sg) | *i ‘he, she, it’ (3sg) | *ï(-)n- 'that' (distal demonstrative) | 40 |
| 37 | | *kɨ- ‘this’ (proximal demonstrative) | *kɨ- ‘that’ (medial demonstrative) | | | *kö 'this' (proximal demonstrative) | 13, 37 |
| 38 | | | | *e- 'this' | *e- 'this' | | 42 |
| 39 | 40 fish | *(y)iwə ‘fish’ | | | *diɣa- ‘fish’ | | 9, 40, 4, 34 |

| | | | | | | | | |
|---|---|---|---|---|---|---|---|---|
| 40 | 43 black | | | | *kara ‘black’ | *kara ‘black’ | 13, 32, 29, 32 | |
| 41 | 45 to stand | *tata- ‘stand, rise, run high’ | *tʌtʌ- ‘run’ | | | | 7, 32b, 8, 32b | |
| 42 | | | *ilʌ- / *ili- 'take shape, arise' | *ili- 'stand (up)' | | | 40, 18 | |
| 43 | 46 to bite | *kam- ‘bite’ | | | *keme- ‘bite’ | *kem- 'gnaw’ | 13, 33, 26 | “Martin 1987: pJ *kama-“. In line with Whitman (1999)[6] and Unger (2000)[7], we reconstruct both consonant and vowel-final verb roots. No internal evidence for final vowel suggests **kam-* here, although commentators’ suggestion would make a better external fit. “No MK kamuŋ” Indeed, not listed here. “pMo derived from *kemi ‘soft bone’”, see SI 2 for why denominal derivation is unlikely. |
| 44 | | | *mili- 'bite' | *mödö- 'gnaw, bite' | | | 25, 37 10, 37 | |
| 45 | 47 back | | | *ar-kan ‘back’ | *aru ‘back’ | *ar-ka ‘back’ | 41, 29 | “no sound correspondences involving pTg *-rk There is a morpheme boundary in pTg *ar-kan ‘back’ (< back-DIM); see SI 2: 21 |
| 46 | 50 what ? | *ka wh-interrogative particle | *ka interrogative particle | *xa- wh-interrogative pronoun | | *ka- wh-interrogative pronoun | 21, 32 | “Some Tungusic languages point rather to *xia” See SI 2: 22 for a straightforward reconstruction of pTg *xa- as a wh-interrogative. The *-i in in pTg **xa-i* ‘who, which one’ is a petrified suffix |
| 47 | 51 child | *wara-pa ‘child’ | | | | *ba:la ‘young animal, child’ | 3, 32, 31, 32 | |
| 48 | | | | *puril ‘child’ | *pure ‘child, seed’ | | 1, 39, 29 | |
| 49 | 53 to give | *tama- ‘give’ | | *tama- ‘pay’ | | | 7, 32, 26, 32 | |

[6] Whitman, J. B. *Primary root shape in Japanese and Korean*. Paper presented at the workshop on Japanese and Korean comparative linguistics at the international conference on historical linguistics, Vancouver, 9–13 August. (1999)

[7] Unger, J. M. Reconciling comparative and internal reconstruction: The case of Old Japanese /ti, ri, ni/. *Language* **76**, 655–681. (2000)

| | | | | | | | | |
|---|---|---|---|---|---|---|---|---|
| 50 | | *(w)ura- ‘sell’ | *pʌlʌ-kʌ- ‘sell’ | *bu:- ‘give’ | | | 3, 39, 30 | |
| 51 | 54 new | *ara- ‘new, pure' | | | *ari- ‘be pure’ | *arï- ‘be(come) pure’ | 41, 29 | |
| 52 | 55 to burn (intr.) | *tak- ‘burn (tr.)’ | *tʌkʌ- /*taki- ‘be on / set fire (intr.)’ | | | *yak- ‘burn (tr.)’ | 9, 32, 14 | “This cognate set mixes transitive and intransitive forms.” Transitivity is marked by way of suffixes added to the verb root, so the transitive and intransitive forms are cognate; see SI 2: 25-26. |
| 53 | 56 not | *ana- negation | *an- negation | *ana- negation | | *an- 'be unbecoming' | 41, 28, 32 | “Old Korean *an-ti* and *an-kʌy* could borrowed from Man. *akû*” See SI 2: 26 with reference to Robbeets[19] (2015: 174-205) for argumentation of why borrowing is unlikely. “pTg reconstruction cannot be derived from the Jur. and Xib. forms according to the correspondence tables” Both are derived from **an-* followed by different nominalizers, see SI 2: 26 and Robbeets[19]. |
| 54 | | | | *e- negation | *e-se negation | *e- negation | 42 | |
| 55 | 58 to know | | *alʌ- ‘know’ | *ala- ‘make known, know' | | | 41, 31 | |
| 56 | 62 to hear | *uka- ‘receive, perceive, hear’ | | | *uka- ‘understand’ | *uk- ‘understand, hear’ | 46, 14, 32 | |
| 57 | 63 soil | *tuti-i ‘soil, ground’ | *tutɨ ‘bank, ridge, ground’ | | | | 7, 38, 8 | |
| 58 | 65 red | | *pɨl(i)-ki- ‘be red’ | *pula- ‘be red’ | *pula-ɣan ‘red’ | | 1, 39b, 31 | “Corr. #39b does not apply since the second vowel is *a in pMo and pTg.” Indeed, we should reformulate 39b with a reflex *PuRV- in Mongolic and Tungusic. “Nugteren (2011: 363) *hulaan” Ultimately, this form can be derived with the deverbal noun suffix -Gan, see SI 2: 28 |
| 59 | 67 to | | *sum- ‘hide, lurk in’ | *sume- ‘hide, conceal’ | | | 23, 38, 26 | |

| | | | | | | | | |
|---|---|---|---|---|---|---|---|---|
| | hide | | | | | | | |
| 60 | 68 skin/hide | *kapa 'skin, bark, shell' | *kap(ʌ)-k 'skin, bark outer layer' | | | *ka:p-ïk 'bark, shell' | 13, 32, 2 | "*a and not *e is not supported by the Koreanic evidence" The Korean evidence supports *a ~ *e alternation for this etymon, see SI 2: 29 |
| 61 | 69 to suck | | | *xökö- 'to suck (breasts)' | *kökü- 'to suck (breasts)' | | 21 37, 14 | |
| 62 | 70 to carry | *əpə- 'carry on back' | *ep- 'carry on back' | *ebe- 'carry' | | | 34, 4, 34 | |
| 63 | 73 to take | *tɨra- 'take, hold' | *tɨlɨ- 'hold up, lift, raise' | | | | 7, 37, 29 | |
| 64 | | | | *al- 'take' | | *al- 'take' | 41, 31 | |
| 65 | 74 old | *muka-si 'be long ago, ancient' | *muk- 'be(come) old' | | | | 25, 38, 14 | |
| 66 | | | | | *kari- 'weaken' | *karï- 'be(come) old' | 13, 32, 29, 40 | |
| 67 | 77 thick | *puta- 'be thick' | *putʌ- 'increase (intr.)' | | *büdü- 'large' | | 3, 38, 10 | |
| 68 | 78 long | *nanka- 'be long' | *nʌlkʌ- 'be(come) old, long (in time) ' | | | | 27, 32b,18, 32b | |
| 69 | | | | | *uri 'before' | *ur$_2$a- 'be long (time/ space)' | 46, 29 | |
| 70 | | | *ola- 'last long' | | *ora- 'be late' | | 43, 29, 32 | |
| 71 | 79 to blow | | *puli- 'blow' | *pu:- 'blow' | | | 1, 38 | "No explanation is provided for the irregular absence of the expected final *r in pTg" The explanation is provided in SI 2: 3. Open monosyllabic with length in Tungusic corresponds to a disyllabic with a liquid onset in the second syllable in the other TEA languages. |
| 72 | 80 wood | *kɨ 'tree, wood' | *kɨl 'tree, wood' | | | | 13, 37 | |
| 73 | | *moro 'woods, wooded hill' | *molo 'hill, mountain' | *mo: 'wood, tree' | *mor wood, tree' | | 25, 35 | "vowel length in Tungusic is unaccounted for" It is |

| | | | | | | | | |
|---|---|---|---|---|---|---|---|---|
|  |  |  |  |  |  |  |  | accounted for in SI 2: 3-4<br>“pJ can only be*moro-, *mərə-, *mɨrɨ-“<br>pJ *moro is a perfect fit<br>“the pM root is *mo-”<br>No, it is pMo *mor-<br>Collective affix pMo *-dun eliminates last sonant of the root, e.g., ni-dün ‘eye’ vs nil-musun ‘tear’. Final -r is preserved in the Baonan form murtóŋ, murtuŋ Kangjia mʉrtʉn. |
| 74 | 81 to run | *pasa- ‘run’ | *pas- ‘hurry’ | | | | 1, 32, 24 | |
| 75 | 82 to fall | *tira- ‘fall, scatter’ | *ti- 'fall, scatter’ | | | | 7, 40 | |
| 76 | 84 ash | *papi ‘ash’ | *pap ‘waste' | | | | 3, 31, 2 | |
| 77 | 86 dog | *inu ‘dog’ | | *ina 'dog' | | | 47, 28 | |
| 78 | 87 to cry/weep | | *uli- 'cry, howl' | | *uli- 'howl’ | *u:lï- 'cry, howl’ | | |
| 79 | 88 to tie | *kuku- ‘tie, wrap' | | *xuku- ‘wrap’ | | | 21, 39, 14, 39 | |
| 80 | | | | | *boɣo- ‘tie, bind, bundle’ | *bog- ‘tie, strangle’ | 3, 36, 16 | |
| 81 | 90 sweet | *ma- ‘to be tasty, sweet’ | *ma- ‘to be tasty’ | | | | 25, 31 | |
| 82 | | | *tʌlʌ- ‘to be sweet’ | *da:l- ‘to be sweet' | | | 9, 32b, 31 | |
| 83 | 91 rope | *tura ‘rope, string, line' | *cul ‘rope, string, line’ | | | | 19, 38, 31 | |
| 84 | 92 shade/ shadow | *kanka ‘reflection, shade’ | *kʌnʌlh < ? *kʌnhʌl ‘shadow’ | | | | 13, 32b, 18, 32b | |
| 85 | 94 salt | *tura- 'bitter, unbearable' | | | | *tu:r$_2$ 'salt' | 7, 39, 30 | |
| 86 | 95 small | *tipi- ‘to be small' | | *čipi- ‘to be small, narrow’ | | | 19, 40, 2 | |
| 87 | 96 wide | *nənpa- ‘become | *nelpʌ- ‘be wide’ | *nepte- ‘become flat and wide' | *nebse- ‘be wide and long' | | 27, 34, 6, 33 | |

| | | | | | | | | |
|---|---|---|---|---|---|---|---|---|
| | | wide and long’ | | | | | | |
| 88 | | | | | *dalba-<br>‘to be wide and flat’ | *yalpa-<br>‘to be wide, flat’ | 9, 31, 6 | |
| 89 | 97 star | *pəsi<br>‘star’ | *peli<br>‘star’ | | | | 1, 34, 20b, 40 | |
| 90 | 98 in | *soko<br>‘depth’ | *soko<br>‘depth, deep inside’ | | | | 23, 35, 14 | |
| 91 | | | | | *örü<br>‘interior' | *ö:r$_2$<br>‘interior' | 44, 30 | |
| 92 | 99 hard | *kata-<br>‘be hard’ | *kata-<br>‘be hard’ | | *kata-<br>‘become hard’ | *kat-<br>‘be hard’ | 13, 32b, 8, 32b | “corr. #32b: expected pK reflex should be *kʌtʌ-“ The weakening process is blocked when a suffix is added to the root. The weakening process has also been blocked in 52 ‘burn’ where there is an alternation between suffix-less pK *tʌkʌ- ‘be on / set fire (intr.)’ and suffixed *taki- (*-i- intr.). Also in MK *talay*- ‘to wheedle, cajole’ with MJ taras- ‘to deceive’. |
| 93 | 100 to crush/ grind | *pinta-<br>‘crush’ | | *pinče-<br>‘crush’ | | | 1, 40, 12, 33 | |
| 94 | | *sura-<br>‘grind, rub, make smooth’ | | | | *sür-<br>‘rub, smear’ | 23, 38, 29 | |
| 95 | | | *niki-<br>'crush, knead' | | *niku-<br>'crush, knead' | yïk-<br>'crush' | 27, 40, 22 | |
| 96 | 101 mountain (hill) (n.) | *yama<br>'mountain' | *yem<br>'mountain' | | | | 46c, 26 | |
| 97 | 102 sit (v.) | *wo-<br>‘sit, be’ | *u-~ *o-<br>modulator | *o:-<br>‘become, make’ | *bol-<br>'become' | *bo:l-<br>‘be’<br>~ *ol-<br>‘sit down, be sitting’ | 3, 35, 31 | |
| 98 | 106 right | | *palʌ-<br>'be at the righthand side' | | *bara-<br>'be at the righthand side' | | 3,32,29 | |
| 99 | 108 grasp (v.) | *tuka -<br>‘grasp’ | *tʌkʌ- ‘get, obtain’ | *tuk-<br>‘take into one’s arms’ | | | 7,39,14 | |
| 100 | 111 raw | | | *niali<br>‘raw’ | | *yaš<br>‘raw’ | 27,40b, 20b | |

| | | | | | | | | |
|---|---|---|---|---|---|---|---|---|
| 101 | 113 two | *puta ‘two’ | *p(ʌ)cak ‘pair’ | | | | 1, 39, 20, 32 | |
| 102 | 114 bottom (n.) | *sita ‘bottom, below’ | *s(i)ta- ‘ground’ | | | | 23, 40, 8, 32 | |
| 103 | 115 lie (down) (v.) | *na- ‘lie down, sleep’ | | *ne:- ‘lay down’ | | | 27, 33 | |
| 104 | 118 year (n.) | | *selV 'year, age' | *se: 'year, age' | | | 23, 34 | |
| 105 | | | | | *jil 'year' | *yïl 'year' | 9, 40, 31 | |
| 106 | 121 weave (v.) | *orə- ‘weave’ | *olʌ ‘unit of woven fibers' | *poro- ‘to spin, rotate’ | *poro- 'tie around, entwine, rotate’ | *pö:r- ‘weave, plait’ | 1, 35, 29 | “pM *-o- and pTg *-o- do not correspond to pTk *-ö- (corr. #35)“ The expected vowel in pTk is *o indeed. |
| 107 | 122 at | *-tu denominal suffix expressing locative relationship | *tʌy locative postnoun 'place' | *-du dative-locative case suffix | *-du denominal suffix expressing locative relationship | | 9, 39 | |
| 108 | | | | *da: locative postnoun 'side' | *-da ~ *-de dative-locative | *-da ~ *-de locative | 10, 32 ~ 33 | |
| 109 | 123 edge (n.) | *pasi ‘edge' | *pas 'outside, beside' | | | | 1, 32, 24 | |
| 110 | | | | *kira ‘edge’ | *kira ‘edge, ridge’ | *kïr ‘edge, ridge’ | 13, 40, 29 | “Nugteren (2011: 415–416): pMo *kirbei ‘edge’” This may be a further nominalization from *kirbe- ‘to trim’. See SI 2: 48 for simplex cognates, not mentioned by Nugteren[27]. |
| 111 | 126 cheek (n.) | *po ‘cheek’ | *pol ‘cheek’ | | | | 1, 35 | “OJ popo is not attested (first in 934)” Attestation in EMJ does not change the reconstruction. |
| 112 | 129 hole (n.) | *ana ‘hole’ | *an ‘interior, inside’ | | | | 41, 28 | |
| 113 | 130 grow (v.) | | *yes- ‘grow grain’ | *üse- ‘grow, plant’ | *ös- ‘grow (animals/ plants)’ | *ös- ‘grow’ (animals/ plants)’ | 46d, 24 | “ pMo *ös- : Dag. reflects distinct etymon” Even if this is the case, the overall reconstruction and comparison remain valid. |
| 114 | | *ura- ‘mature, ripen’ | | *ure- ‘grow, ripen’ | *ur-ga- 'grow' | *ur ‘growth, excrescence’ | 45, 29 | “OJ ure- does not exist (first attestation 1603)” Later attestation does not invalidate the reconstruction. “Nugteren (2011: 533) *urgu ‘to come up’”, See SI 2: 50 for comment on Nugteren[27] |

| | | | | | | | | |
|---|---|---|---|---|---|---|---|---|
| | | | | | | | | and explanation of *ur-ga- |
| 115 | 132 belly (n.) | *para ‘stomach’ | *pʌ(l)i ‘belly’ | | | | | |
| 116 | 136 dig (v.) | *uka- ‘dig, excavate' | | | *uka- ‘dig, excavate' | | 45, 22, 32 | |
| 117 | 138 hot | *yu 'hot water' | *tʌ- ~ ti- ‘be hot, warm’ | *du:l- 'be warm' | *dula- 'be(come) warm' | *yïlï- 'be(come) warm' | 9, 39, 31 | “No explanation for the absence of the expected *l in pK” Given in SI2: 43, 52 “only pK *tʌs-“ SI2: 51 explains why -s- is considered a suffix. “pMo *dulaan ‘warm’” SI2: 51 explains -*GAn* as resultative deverbal noun suffix |
| 118 | 140 remain (v.) | *nəkə - ‘remain, be left over’ | *nek ‘remnant' | | | | 27, 34, 14 | |
| 119 | | *ama- ‘remain, be left over’ | | *eme:- 'remain, leave' | | | 41b, 26, 33 | |
| 120 | 141 cold | | | *bege- 'be cold, freeze' | *bege- 'be cold, freeze' | | 3, 33, 16, 33 | |
| 121 | | | | *gilči- 'be cold' | | *kïš 'winter' | 15, 40, 20b | |
| 122 | 144 thin | | | | *nari- 'be thin' | *yarï- 'be thin, poor' | 27, 32, 29, 40 | “ Nugteren (2011: 452) *narïn ‘thin, fine’.” Root *nari- supported by various Mongolic derivations with different suffixes; see SI2: 53-54. “Comments SI2: 52 gives meanings that differ from the concept ‘thin’ for the forms cited in this cognate set: Kaz. ‘to be poor’, Kir. ‘lean, skinny’, Tuv. ‘to spend, consume’. SI2 Table 3.9 gives no correspondence for the initial consonants involved here.” Leaving out the three participants does not invalidate the Turkic reconstruction. |
| 123 | 147 sour | *su- ‘be sour’ | *suy- ~ *sʌy- ‘be sour’ | | | | 23, 38 ~ 39 | |
| 124 | 149 day (n.) | | *nal 'day' | | *nara-n 'sun' | | 27, 32, 31 | |
| 125 | 152 white | *sera- ~ *sero 'be white' | *siala- 'be white' | *sia:ra- | *siara 'yellow' | *sia:rï- | 23, 40b, 29, 32 | “ this is the same root as ‘yellow’” The original meaning was |

| | | | | | | | | |
|---|---|---|---|---|---|---|---|---|
| | | | | 'be white, light’ | | 'be white, yellow’ | | ‘white’ and it developed into ‘yellow’; see SI 2: 54-56<br>“Long vowels are unaccounted for” Suprasegmental correspondences (between tone and length) are a promising research topic for the future, but not indispensable to establish cognates.<br>“pJ *siro-“ on double origin for OJ *i*, see SI 2: 54<br>“pMo *ia does not appear in SI2 Table 3.8.” It is indeed more likely that the Mongolic form is a borrowing from Turkic and that the reconstruction is pMo *šara, see SI 2: 56 |
| 126 | 153<br>sew (v.) | *nup-<br>‘sew' | *nupi-<br>‘sew' | *nup-<br>'prick, pierce’ | | | 27,<br>38, 2 | |
| 127 | | | *sili<br>'thread' | *sira-<br>'sew together' | *siri-<br>'sew, stitch, quilt' | *sïrï-<br>'sew, stitch, quilt' | 23,<br>40, 29 | |
| 128 | 161<br>many | *opə-<br>'be many' | *op-<br>‘be enough' | | | | 43, 2 | |
| 129 | 162<br>chew (v.) | | | | *kebi-<br>'chew, ruminate' | *ke:b-<br>'chew, ruminate' | 13,<br>34, 4 | |
| 130 | 164<br>wet | *uru-<br>‘be wet’ | *uli-<br>‘soak’ | *ula-<br>'wet, soak' | | | 45, 31 | |
| 131 | | *nura-<br>‘be(come) wet’ | | | *nor-<br>‘be(come) wet' | | 27,<br>36, 29 | |
| 132 | 165<br>four | *yɨ<br>‘four’ | | *dü-<br>‘four’ | *dö-<br>‘four’ | | 9, 37 | |
| 133 | 166<br>soft | *mɨrɨ-<br>‘be weak’ | *mɨlɨl-<br>‘be(come) soft' | | | | 25,<br>37,<br>29, 37 | |
| 134 | | *yapa-<br>'be soft, weak' | *yepu(-)y-<br>'be(come) weak' | | | *yaba-<br>'be soft, mild, quiet' | 46c,<br>32, 4,<br>32 | |
| 135 | 167<br>look (v.) | | | *kara-<br>'look, watch' | *kara-<br>'look, watch' | | 13,<br>32b,<br>29,<br>32b | |
| 137 | 169<br>that | *a-<br>'that' (distal demonstrative) | | | | *a-n-<br>'that' (distal demonstrative) | 41 | |
| 138 | | *ə-<br>'that' (distal demonstrative) | | *e-<br>'this' | *e-<br>'this' | | 42 | |
| 139 | 170<br>cut (v.) | *kira-<br>‘cut’ | | *giri-<br>‘cut out’ | | *kïr-<br>‘cut, break, scrape’ | 15,<br>40, 29 | |

| | | | | | | | | |
|---|---|---|---|---|---|---|---|---|
| 140 | 171 mother | *əmə 'mother' | *ema 'mother' | *eme 'mother (in law)' | *eme 'woman' | *eme 'mother, old woman' | 41b, 26, 33 | “nursery word is not really reliable” As the concept is part of the pre-set Leipzig Jakarta basic vocabulary list, we need to consider it. This was explained in SI 2: 63. |
| 141 | 176 warm | *nuku- ~ *noko- ‘to be warm’ | *nuk- ~ *nok- ‘to be(come) warm' | | | | 27, 35 ~ 38, 14 | |
| 142 | 177 cover (v.) | | | | *bürü- 'cover' | *bürü- 'cover up' | 3, 38, 29, 38 | |
| 143 | 178 woman (n.) | *me 'woman' | *mye 'woman' | | | | 25, 40b/c | |
| 144 | 179 deep | *kipa- ‘have a rapid decline' | *kip-kʌ- ‘be deep’ | | | | 13, 40, 2 | |
| 145 | | *puka- 'be deep' | *pokʌ- ~ *pukɨ- 'be deep' | | | | 1, 36 ~ 38, 14 | |
| 146 | 180 above | *u-pa-(C)i 'above, upside' | *u-kɨ 'above, upside' | | | | 45 | |
| 147 | 183 other | *pəka ‘other, besides’ | *pekɨ- ‘come next, be beside’ | | | | 1, 34, 14 | |
| 148 | 185 left | | | | *sola-gai ‘left’ | *so:l ‘left’ | | |
| 149 | 186 rise (v.) | *ɨkɨ- 'rise, raise’ | | *ög(ö)- 'rise, go up, mount’ | *ög- 'go up' | *ög- 'raise, heap up' | 44, 16 | |
| 150 | | *taka- ‘be high, elevated’ | *tekɨ- 'make go up, increase' | *dege- ‘rise, go up, fly' | *dege- ‘go up' | *yeg ‘upper part’ | 9, 33, 16 | |
| 151 | | | | | *kali- 'rise, fly' | *kalï- 'rise' | 13, 32, 31, 40 | |
| 152 | 189 break (v.) | *kaka- 'break’ | | *xaka- ‘cut off, tear off’ | *kaka- ~ *kaga- ‘break' | *kak- ‘beat, injure' | 21, 32b, 22, 32b | |
| 153 | | *yanpu- ‘break, tear, split' | | *delpe- ‘break, split' | *delbe- 'break, burst’ | | 9, 33, 6 | |
| 154 | | | | *butu- 'break' | *buta- 'break' | | 3,39, 8 | |
| 155 | 191 spin (v.) | *tumu 'spindle' | | *tomu- 'spin' | *tomu- 'spin' | | 7, 36, 26 | “Ryukyuan forms unambiguously point to pJ *u and not *o” Indeed, Ryukyuan evidence points to *u. Nevertheless correspondence 36; see Table 5 below. |
| 156 | 194 open (v.) | *aka- ‘open’ | *aki- ‘open’ | | *agu- 'open' | | 41, 16 | |
| 157 | | | *yele- 'open' | | *yara- 'open' | *ya:r- 'split' | 46c, 29 | |
| 158 | 204 father | | *apa ‘father’ | | *aba ‘father’ | *apa~ aba ‘father’ | 41, 4, 32 | *CaCa should |

| | | | | | | | | |
|---|---|---|---|---|---|---|---|---|
| | | | | | | | | correspond to pK *ʌpʌ (corr. #32b)” As there is no initial consonant here, corr 32b does not apply. |
| 159 | 207 five | | *ta- 'five' | | *ta- 'five' | | 7, 32 | |
| 160 | 212 green | | | *nog- | *nogo- | | 27, 35, 16 | “No sound Correspondence for pTg *ń-“ Palatalization in Tungusic is secondary: assimilation to colour suffix *-gian See SI 2: 74 |

#### 1.2. Comparative sets relating to subsistence

As for our etymologies relating to subsistence, Tian et al. use some questionable figures. They write “31 comparisons follow the proposed sound correspondences, only 29 could be reconstructed at the Proto-Transeurasian level, only 27 appear in more than three branches, only 22 have comparable semantics, and only 9 items belong to the realm of agropastoral vocabulary”, inferring from this that “[t]he identification of Proto-Transeurasian speakers with early millet farmers of the West Liao River area is thus not supported by empirical linguistic evidence.”

Table 2 summarizes the comparative sets relating to subsistence aimed at the reconstruction of the cultural environment of the speakers of Proto-Transeurasian advanced in our Article (SI 5). The final column of Table 2 summarizes our item-by-item reply to the “evaluation of the agropastoral vocabulary” by Tian et al (SI: 35-47). As in Table 1, we inserted our full replies in the pdf comment function of the supplementary information by Tian et al. and maintained the same principles for colour coding.

First, it should be noted that given the strong opposition of the authors to Transeurasian affinity and the high level of expertise of the linguistic team, it is reassuring that we can agree on a core of 31 agropastoral etymologies that correspond regularly. This recognition makes us hopeful that a consensus is within reach.

Second, the claim that “only 22 have comparable semantics” is highly subjective. A word consists of sound and meaning. Of course, not just sounds change over time, meanings are expected to shift as well. Such changes tend to be less regular and verifiable than sound change. For semantic comparison, linguists use some rules of the thumb but there are no regular and systemic correspondences as in the case of phonological comparison. One guideline in comparing the meaning of two words is that a certain degree of semantic latitude is acceptable to the extent that the assumed meaning change can be shown to have occurred elsewhere, in other languages of the world. A recurrent and stunning claim by Tian et al., for instance, is that a word designating a tree cannot be compared to the nut produced by that tree: see Table 2 (26), (27) and (29). However, across the languages of the world trees and their nuts are often co-lexicalized, e.g., in English chestnut, walnut, etc. This observation is sufficient to legitimize the comparison between the word for a tree and that for the nut it produces.

Another guideline is that semantic latitude is permissible to the extent that both words have a semantic denominator in common that can be explained as the source of both meanings. For instance, Tian et al. object that “forms ‘to milk’ have nothing to do with fermentation”, see Table 2 (14) and (16). However, as explained in detail in our Article (SI 5: 18), due to lactose intolerance of contemporary Mongolic and Turkic populations who subsist on dairying, milking must always be followed by fermentation to make the milk consumable. This explains the close semantic connection between ‘to ferment’ and ‘to milk’ in these languages and indicates that the semantic source of the compared words is ‘to ferment’.

Third, the criticism that “only 9 items belong to the realm of agropastoral vocabulary” disregards our broader interest in subsistence vocabulary, not exclusively restricted to agricultural terms *per se*. Besides, it implies an unrealistic expectation that all contemporary words derived from an agricultural source 9000 years ago would still preserve an agricultural meaning today. This is for instance reflected in the objection that “[t]he forms [for ‘pig’] are not semantically comparable since the Turkic forms do not denote an animal of the Suidae family but cattle, camels, or horses”; see Table 2 (31). However, semantic change is expected to take place over millennia of linguistic history and the direction of the reconstructed change from ‘pig’ to pastoral animals is consistent with our analysis that Proto-Turkic was the first daughter language to enter the Eastern steppe, where from the Bronze Age onwards pastoralism was added to its agricultural package. This may explain why several Turkic words that were agricultural in origin shifted to a more pastoral or generic meaning in the course of their history.

Finally, Tian et al. regard their partial refutation of our etymologies relating to subsistence as sufficient ground to reject our association of the Proto-Transeurasian language with early millet farming. However, in our Article, this inference is not based on a single strand of evidence, such as the reconstruction of subsistence vocabulary. By contrast, our approach triangulated multiple datasets, methods and disciplines to reach this conclusion.

**Table 2** Summary of the comparative sets relating to subsistence aimed at the reconstruction of the cultural environment of the speakers of Proto-Transeurasian, with indication of criticized members.

| No | MRCA | Proto-Japonic | Proto-Koreanic | Proto-Tungusic | Proto-Mongolic | Proto-Turkic | "Comment" and short reply |
|---|---|---|---|---|---|---|---|
| 01 | 1 pTEA *pata 'field for cultivation' | *pata '(dry) field' | *patʌ-k '(dry) field' | | | *(p)ata 'delimited field irrigated for cultivation' | "we expect pK *pʌtʌ- and not *patʌ-" Expected vowel lenition is blocked through petrification of place suffix<br><br>"Alternatively, OT atïz derived via the collective suffix -z from the verb at- 'to throw, to remove'" The collective suffix is a denominal suffix, it does not derive from verbs. |
| 02 | 2 pTEA *muda 'uncultivated field' | *muta 'marshland' | *mutʌ-k 'dry land' | *muda plain, open field, | | | " should correspond to pK *-l- (#10)" Block of intervocalic t > l - lenition in pK, maybe due to petrification of place suffix "second vowel should be pK *a" A correspondence is defined as CVC with exclusion of an eventual root-final vowel " None of the forms involved has the meaning 'field'" The forms mean marshland, dry land, meadow, etc. The fact that we can reconstruct various distinctions for 'field' suggests that agriculture played a central role in the subsistence of the ancestral speakers |
| 03 | 3 pTEA *iuse- 'to plant, grow (plants)' | | *yes- 'to grow grain' | *üse- ~ üsi- 'to plant' | *ös- 'to grow (of plants/animals)' | *ös- 'to grow' | "No Koreanic verb 'to grow grain' can be reconstructed." Meaning 'cooked cereal' and 'ear of grain' may indeed be semantically too distant. "not anyway specific to agriculture but generic verbs meaning 'to grow, to multiply, to |

| | | | | | | | |
|---|---|---|---|---|---|---|---|
| | | | | | | | increase'." Tungusic and Mongolic verbs are explicitly in an agricultural context. Turkic has broadened its meaning |
| 04 | 4 pTEA *urə- 'to grow, ripen (of plants)' | *ura- 'to mature, ripen (of plants)' | | *ure- 'to grow, ripen (of plants)' | *ur-ga- 'to grow (of plants)' | *ur 'growth, excrescence' | "the absence of cognates in Ryukyuan … forbids its comparison with other languages." The comparative method can be applied even if a cognate is missing in certain languages or dialects but present in others. |
| 05 | 5 pTEA *pisi- 'sprinkle with the hands, sow' | | *pis- 'to sprinkle, scatter, sow' | *pisi- 'to sprinkle with the hands' | *pisü- ~ *pesü- 'to sprinkle, scatter | | "the forms … just mean 'to sprinkle, to scatter'" The meaning 'to sow' can be reconstructed on the basis of the derived nouns for 'seed, seedling' and 'millet seed' in the etymology (6); see SI5: 9-10.<br><br>"MK spih- probably contains the intensive prefix s-, the noun pí 'rain' and the verbalizer ho- 'to do (Martin 1996: 47)." We accept Martin's explanation. |
| 06 | 6 pTEA *pisi-i (sow- INS.NMLZ) 'seed, seedling' | | *pisi 'seed; lineage' | *pisi-n 'seed' *pisi-ke 'broomcorn millet (Panicum miliaceum)' | *pisi ~ *pesi 'origin or base of a plant' | | "MK psí reconstructs as pK *pʌsi or *pisi, but not *pisi" In line with Ramsey[8],[9] (1993: 438-439, 1997), tonic monosyllabic words with complex initials and minimal vowels (MK o, u) or i are created through the loss of a first-syllable unaccented minimal vowel. Robbeets[10] (2005: 365) suggested that the lost vowel can also be *i*. pK **i* is supported by external |

[8] Ramsey, S. R.. Some remarks on reconstructing earlier Korean. *Language Research* 29, 433–441. (1993)
[9] Ramsey, S. R. in Kim-Renaud, Y. K. (ed.), *The Korean alphabet: Its history and structure*, 131–143. (University of Hawai'i Press, 1997)
[10] Robbeets, M. Is Japanese related to Korean, Tungusic, Mongolic and Turkic? (Harrassowitz, 2005)

| | | | | | | | |
|---|---|---|---|---|---|---|---|
| | | | | | | | comparison, whether one argues for borrowing (as Tian et al. SI do and Vovin[11] 2006: 260 does for Ma. fisike 'millet' from pre-MK psi 'seed') or inheritance.<br><br>"Mongolic forms mean 'basis, origin'" The Mongolic forms mean 'stem of a plant, stalk of grain, etc.' The stem or base of a plant is comparable to a seed or seedling, which is also at the origin of a plant. |
| 07 | 7 pTEA *kipi ~ kipe 'components that are removed from the grain harvest, barnyard grass (Echinochloa crusgali)' | *kinpi ~ *kimi ~ *kipi 'millets for human consumption such as barnyard or broomcorn millet' | *kipi<br><br>'barnyard millet (*Echinochloa esculenta*)' | *kipe<br><br>'components that need to be removed from the grain harvest, barnyard grass (Echinochloa crusgali)' | | *kepe-k<br><br>bran (from millet, barley), chaff | "p is expected in Japonic" Intervocalic /p/ was realized as [b] or [p] in OJ at a time when intervocalic /b/ was realized as [mb]. Confusion between intervocalic /p/ and /b/ may have arisen when [mb] became pronounced as [b].<br><br>"MK phí 'barnyard millet' can not go back to *kipi" Our evidence allows for the loss of high front vowels in tonic monosyllables with complex intials; see 06 above.<br><br>" pTk *e is irregular" Indeed |
| 08 | 8 pTEA *amʊ 'cooked cereal, (millet) gruel' | *amai 'cereal starch' | *amʌ 'cooked cereal' | | *amu-n 'cooked cereals; millet' | | "pJ is a derivation from ama- 'sweet'" Accepted. |
| 09 | 9 pTEA *sarpa 'spade' | | *salp(V) 'spade' | | | *sarpan 'wooden plough (breast), spade' | "*CaCa should yield pK *ʌ" The sequence is CaCCa, thus corr. 32 |
| 10 | 10 pA *tari- 'to cultivate' | | | *tari- 'cultivate; to sow, plant' | *tari- 'prepare the soil for cultivation; to sow, plant' | *tarï- 'to cultivate (land); to prepare the soil for cultivation | |

[11] Vovin, A. in Pozzi, A. et al. (eds.), Tumen jalafun jecen akū: Manchu studies in honour of Giovanni Stary, 255–266. (Harrassowitz, 2006)

| | | | | | | | |
|---|---|---|---|---|---|---|---|
| | | | | | | ; to sow, disperse’ | |
| 11 | 11 pMoTg *pure ‘seed, sprout, offspring’ | | | *puri 'sprout, offspring, child' | *püre ‘seed, fruit; offspring, child’ | | “do not belong to the agricultural lexicon” Allow for the reconstruction of 'seed' in pTEA. |
| 12 | 12 pJK *non ‘field for agriculture’ | *no ‘field’ | *non ‘irrigated field, rice paddy field’ | | | | “The Japonic form (‘uncultivated plain’) invalidates the comparison” The common semantic denominator is 'field' but as there is a probable external donorword relating to agriculture, it is more likely that the designation 'agricultural' got lost in Japanese than that it concerns a semantic specification in Korean. |
| 13 | 13 pJK *mati ‘delimited plot for cultivation’ | *mati ‘delimited plot for cultivation’ | *mat(i)-k ‘delimited plot for cultivation’ | | | | “the second vowel in pK can only be *ʌ” According to Robbeets (2005: 365) also i, but note that matches are defined as CVC correspondences, irrespective of root-final vowel<br><br>“A plausible etymology would be ma ‘space, interval’ + ti ‘path’, hence the general meaning ‘delimited area, section, plot’, in which case there is no relationship with agriculture.” OJ *mati* means '町 a measurement of land, a sector,' and '坊 divisions in a town.' in the *Wamyōshō*. *Nihon shoki* points to the measurement of land as the original meaning. In Shuri (Ryukyuan), the word is preserved as -*masi*, a counter for rice paddies. Hence an agricultural reconstruction is more sensible than the forced segmentation proposed by Tian et al. |

| | | | | | | | |
|---|---|---|---|---|---|---|---|
| 14 | 1 pTEA *saga- 'to ferment' | *saka- 'ferment, be in heat, bloom' | *sak- 'ferment, to rot' | | *saga- ‘to ferment, reduce (food); to milk’ | *sag- ‘to milk; to extract’ | “The meaning ‘to ferment’ is not attested in Mongolic.” The meaning 'to ferment' is actually attested, see Khalka *saga-* 'to overflow, rise up, be overfull of food, to trickle, to ferment, turn sour’[12]. In addition to ‘to milk, reduce (food)’, some languages also preserve ‘rise up (of food)’ This indicates a connection with fermentation. “Mongolic and Turkic forms ‘to milk’ have nothing to do not with fermentation” Especially for Turkic and Mongolic speakers, there is an undeniable link between ‘milking’ and ‘fermentation’, explained in SI 5: 18. Due to lactose intolerance, milking must always be followed by fermentation to make the milk consumable. |
| 15 | 2 pTEA *sugu- ‘to ferment’ | | *su(k)u- 'to make alcohol' | | *su(g)a- 'to milk' | *sug- ‘to procure cheese’ | “pMo *u and pTk *u only correspond to pK *ʌ and not *u (#39)” Accepted |
| 16 | 3 pTEA *su:- ‘to reduce/preserve food by fermentation’ | *su- 'to be sour, fermented in vinegar' | *swu-y- ‘to be sour’ | | *sü- ‘strain, filter, skim off, reduce (food)’ | *sü:- 'to filter, strain (milk or soup), reduce (food)' | “vowel correspondence?” Cor. 38<br><br>“only the Japonic and Koreanic forms are related to fermentation” The Turkic and Mongolic verbs cover all steps of the milk fermentation process: strain and filter milk and then reduce it to fermented dairy products |
| 17 | 4 *bilče- ‘to mix food with a liquid’ | *pisi ‘fermented liquid’ | *pici- ‘ferment (liquid), brew, | | *bilca- 'mix a liquid with food’ | *bilči- 'to ripen, to churn (milk, butter), to | “ OJ *pisipo* is obviously formed on the root sipo ‘salt’” This derivation is not without problems, |

[12] Pyurbeev. G. Ts. Bol’shoj akademicheskij mongol’sko-russkij slovar’ [Large Academic Mongolian-Russian Dictionary] (Academia Publ., 2001–2002)

| | | | | | | | |
|---|---|---|---|---|---|---|---|
| | | | make dough’ | | | stir up, beat a liquid in order to make butter’ | but we can leave the Japanese form out.<br><br>“No evidence for reconstructing a final *i in pK for this root” There is evidence because Cheju and Kangwondo preserve a final i in ppici-, see SI 5: 22<br><br>“pMo cannot be reconstructed as ‘mix a liquid with food’” Various meanings in Mongolic refer to food production, e.g. 'to be of thin consistency (of though, batter)’ and 'to flatten completely, squash into pulp, crush, pound (millet) (intr.)’ Mongolic was borrowed into Manchu bilca- ‘to mix (flour)' with exactly the reconstructed meaning; see SI 5: 22 |
| 18 | 5 pTEA *silɔ ‘broth, soup, juice; liquid food extracted from vegetables, fruit or meat’ | *siru ‘broth, soup, juice’ | | | *silö ‘meat broth, soup, juice’ | | “The second vowel is irregular” Matches are defined as CVC correspondences, irrespective of root-final vowel |
| 19 | 6 pTEA *niku- 'to crush to pulp’ | | *niki- 'to crush to a pulp, knead (flour)' | | *niku- 'to crush, knead (flour) ' | *yïk- 'to crush’ | “None of the forms compared are specific to agriculture” The Korean and Mongolic forms ‘crush, knead (flour)' relate to food production. We reconstruct cultural vocabulary beyond agriculture to determine the cultural environment and subsistence strategies of the Transeurasian speakers. |
| 20 | 7 pTEA *suru- 'to grind, rub' | *sura- 'to grind, rub' | | *suru- ‘to grind’ | | *sür(ü)- 'to rub, smear' | “None of the forms compared are specific to agriculture” Even if not directly associated with food production, the frequency of verbs for ‘grinding’ is observed and related to food |

| | | | | | | | |
|---|---|---|---|---|---|---|---|
| | | | | | | | production on SI5: 19. |
| 21 | 8 pTEA *simi- ‘to soak (food)’ | *sima- ‘to soak, permeate’ | | | *sime- ‘to soak (food), moisten, suck up’ | | “forms do not denote anything related to food production or preservation” Derived nouns in Mongolic are food products (SI 5: 24) Even if not directly associated with food production, the frequency of verbs for ‘soaking’ is observed and related to food production and/or textile technology on SI5: 19. |
| 22 | 9 pTEA *ulu- ‘to soak, wet’ | *uru- ‘to irrigate, make wet’ | *uli- ‘to soak’ | *ula- ‘to wet, soak’ | | | “the pJ form reconstructs as *oro” See SI 2: Table 3.2. Amami reflex lacks, so we cannot distinguish between pJ *o and *u, but external comparison suggests *u. |
| 23 | 10 pTEA *nɔr- ‘to soak’ | *nura- ‘to be(come) wet’ | | | *nor- 'to be(come) wet' | | “pJ *u does not regularly correspond to pM *o” Cor. 36; see Table 5. |
| 24 | 11 pTEA *deb- ‘to soak’ | | | *debe- 'to make wet, paste, paint' | *debe- ‘to soak (clothes), make wet' | *yebi- ‘to become wet, soak' | “None of the forms compared are specific to agriculture” Note meaning ‘paint’ in Tungusic and ‘soak (clothes)’ in Mongolic. The frequency of verbs for ‘soaking’ is observed and related to textile production on SI5: 19. |
| 25 | pJK *kama- ‘to soak, brew’ | *kamə- ‘to chew, brew, dye’ | *kʌmʌ- 'to put in a bath' | | | | “primary meaning of Japanese kam- is ‘to bite’” The common semantic denominator in Japanese is ‘to soak’ as ‘dying and brewing are both done by soaking and brewing is associated with mastication.<br><br>“The Japonic and Koreanic forms have incompatible semantics” The semantics ‘to soak’ and ‘to put in a bath’ are compatible. |
| 26 | 1 pTEA *kuru ‘edible nut used for starch production | *kuru ‘walnut, chestnut’ | *kul ‘oak < ? walnut’ | *kuri ‘pine | | | “Koreanic form denotes a tree and not a nut, so that it |

| | | | | | | | |
|---|---|---|---|---|---|---|---|
| | such as walnut, acorn, chestnut or pine nut' | | | cone, pine nut' | | | cannot be compared" Across the TEA languages (and beyond) trees and their products are often co-lexicalized, English chestnut, walnut, etc. are also used for both the tree and the nut. Note that the Korean tree name is often specified by namu 'tree'. |
| 27 | 2 pTEA *xʊsi 'edible nut used for starch production such as walnut, acorn, chestnut or pine nut' | *kusi 'chestnut' | | *xusi 'acorn' | *kusi 'walnut' | | "OJ kusi is …a dialectal word cognate with kuri" Even so, we cannot exclude that the kusi-kuri alternation was already present in pTEA. "Mongolic does not denote a nut but a tree" Trees and their products are often co-lexicalized; see above. WMo. *qusiga* 'walnut, nut; testicles' clearly denotes the tree as well as the fruit. "not directly related to food resource" In the West Liao River region in the Neolithic, farmers processed acorns for starch at least as frequently as millets. Precisely nuts such as walnut, acorn, chestnut or pine nut, which were targeted for their starch and consumed by Xinglongwa people, turn up in our etymologies. |
| 28 | 3 pTEA *abu 'plant of the Althaea genus with roots rich of starch' | *apu 'hollyhock (Althaea rosa)' | *apo-k 'marshmallow (Althaea officinalis)' | | *abu 'marshmallow (Althaea officinalis)' | | "Since the Japonic form does not denote a nut, it is not comparable with the Koreanic form." None of the languages denote a nut, all refer to the Althaea root native to Northern China, used medicinally. "pK *-k is left unexplained" *-(ʌ/i)k edible plant suffix, e.g., MK chulk 'kuzu (arrowroot, |

| | | | | | | | |
|---|---|---|---|---|---|---|---|
| | | | | | | | edible)', MK ·milh 'wheat', MK ·phoch ~ ·phosk 'red bean', etc.; see SI 5: 8. "no cognates in other Mongolic languages than Kh." WMo. abuɣa, 'marshmallow (Althaea officinalis) next to Khal. avga |
| 29 | 4 pJK *pami 'edible true nut, i.e. dry fruit with only one seed such as acorn, hazelnut and chestnut' | *pami 'edible true nut such as hazelnut or acorn' | *pami 'edible true nut such as chestnut or hazelnut' | | | | "Japonic form does not denote a nut" OJ turupami$_1$ is Quercus acutissima and by extension the name of its nut, the acorn; EMJ pasipami is Corylus heterophylla, the Asian hazel. "K *pamʌ and not *pami" The rising tone in MK suggests a disyllabic origin, while the palatal glide in KN paym and JB paym indicates diphtongization from an original stem-final *-*i*. The suggested reconstruction pK *pamʌ would be a regular match as well given that matches are defined as CVC correspondences, irrespective of root-final vowel |
| 30 | 1 pTEA *inu ~ *ina 'dog' | *inu ~ *ina 'dog' | | *inu:-ke 'dog, wolf' | | | "Dog precedes agriculture: not relevant" Relevant because only two potentially domesticated animals can be reconstructed: 'dog' and 'pig', indicating that pastoral vocabulary developed secondarily; SI 5: 28-29. "initial pTg *ŋ- is usually reconstructed, but there is no such consonant in the correspondence tables." According to Poppe[13] (1964: |

[13] Poppe, N. in Spuler, B. Et al. (eds.), 1–16 (Brill, 1964)

| | | | | | | | |
|---|---|---|---|---|---|---|---|
| | | | | | | | 4) pTg *ŋ- is the result of secondary assimilation to following nasal of PTg *g-, which explains why *ŋ- is absent from the correspondence table. “no basis for reconstructing the second vowel as pTg *u” Even *ŋõke* ‘male (of dog, wolf, fox)’, Sibe *juxə* ‘wolf’ and Ma. *nuxere* 'puppy' all reflect pTg *u |
| 31 | 2 pA *toru 'young male pig' | | | *toro 'male pig' | *toru 'young male pig' | *to:ru 'young cattle/camel/horse' | “the Turkic forms are not semantically comparable” The original meaning ‘pig’ was repurposed as animal names related to Bronze Age pastoralism, such as 'cattle,’ ‘camel’ or ‘horse'; see SI 5: 29. |
| 32 | 3 pMoTg *uli ‘pig’ | | | *uli-gan ‘pig’ | Khitan *uil(e) 'pig' | | “only appears in Khitan” Nevertheless thought-provoking and deserves mention |
| 33 | 1 pTEA *nap- ‘to make rope’ | *nap- ‘to make rope’ | *nap- ‘twist, spin’ | *nap- ‘to make rope’ | | | “PK *p is speculative and fails to explain the presence of a h” *p is actually reflected in dialectal forms and in derivations in Korean: K kkunapwul, KN napi-, JJ napi CN napi. The velar fricative *-G- may derive from *-p- (or -*k-) and yield -h-. “The Tungusic forms do not mean ‘rope’ or ‘to make rope’” The Tungusic words designate a rope or a thong of a flexible material used for securing the skis. They are probably derived from a verbal base through the resultative deverbal noun suffix *-ki. Plausible semantics for the underlying verb |

| | | | | | | | |
|---|---|---|---|---|---|---|---|
| | | | | | | | are thus ‘make rope, make thongs’ |
| 34 | 2 pTEA *nup- ‘to sew' | *nup- ‘to sew, stitch’ | *nupi- ‘to sew, quilt’ | *nup- ‘to prick, pierce’ | | | “The Tungusic form is not semantically comparable” 'pierce' represents a common colexification of 'sew'[14] |
| 35 | 3 pTEA *siri- 'to sew’ | | *sili 'thread' | *sira- 'to sew together, tie together' | *siri- 'to sew, stitch, quilt' | *sïrï- 'to sew, stitch, quilt' | “Tungusic reconstruction of a verb specifically meaning ‘to sew’ is not justified” Since Orok, Ma., Sibe, Olcha, Na., Oroch, Ud. all share meanings ‘to sew and/or tie together’ and as the deverbal nouns mean ‘thread’, we consider the semantic reconstruction legitimate. |
| 36 | 4 pTEA *pɔrɔ- ‘to weave (cloth)’ | *oro- ‘to weave’ | *olʌ ‘unit of woven fibers’ | *poro- ‘to spin (nettle and hemp threads); to rotate, turn’ | *poro- ‘to tie around, entwine; rotate, turn’ | *pö:r- ‘to plait, weave’ | “pJ should be *or(o)- or *ər(ə)-“ *or(o)- represents a match<br><br>“pJ and pK *p- expected” SI5: 33-34 explains conditioning factor: pJK *p drops before a (long) rounded pJK *o(:) Tungusic and Mongolic are generic verbs that do not mean ‘to weave’ but ‘to spin’. The deverbal nouns in Tungusic mean ‘spindle; spinin wheel; device for weaving nets’, which shows a semantic connection to complex textile technology, which justifies comparison. Even if the Mongolic meaning has broadened to ‘to tie around, entwine; rotate, turn’, the generic meaning is still semantically comparable. “pTk has *ö instead of the expected *o (#35)” Accepted |

[14] List, M. et al. The database of cross-linguistic colexifications, reproducible analysis of cross-linguistic polysemies. DOI: 10.1038/s41597-019-0341-x (2019)

| | | | | | | | |
|---|---|---|---|---|---|---|---|
| 37 | 5 pTEA *tәmʊ- 'to spin' | *tumu 'spindle' | | *tom(u)- 'to spin' | *tomu- ~ *tamu- 'to spin' | | "not attested in OJ" But well-attested in historical and contemporary varieties of Japanese in addition to Ryukyuan. "pM *-o- and pTg *-o- do not correspond to pJ *-u- (corr. #35–36)" Correspondence is no 36; see Table 5 below. |
| 38 | 6 pTEA *giri- 'to cut (cloth)' | *kira- 'to cut (e.g., cloth)' | | *giri- 'to cut out (e.g., cloth, paper, pelt)' | | *kïr- 'to cut, scrape' | "not related to textiles but just generic terms for cutting." Relation to textile is explained on SI5: 36-37 |
| 39 | 7 pJK *parʌ- 'to sew' | *par-i 'needle' | *palʌ-l 'needle' | | | | "better reconstructed as pJ *pari" OK " it is a case of corr. #39b." No, corr. 32 "LMK palol 'needle' "is .. hapax legomenon" Nevertheless, the form is attested and we cannot just ignore it. There is evidence for confusion of lVl and nVl sequences in Late Middle Korean and pre-MK *palól > panól as analogy to common nouns ending in -nol.[15,16,17] |
| 40 | 8 pJK *paca- 'to weave (cloth) with a loom' | *pata- 'to weave' | *pʌcʌ- 'to weave' | | | | "The MK form is pcó-, not specific to textile production" LMK ·pco- means 'to weave' with broadening of the meaning in later varieties (Park 2010[18] 10:1015: `vt. To weave. To interlace the woof and the warp to produce materials such as fabric.') |
| 41 | 9 pJK *pu- | *pu | *pu- | | | | "pJ reconstruction of this etymon as a verb meaning 'to |

[15] Martin, S. E. Lexical evidence relating Korean to Japanese. *Language* **42**, 185–251 (1966).
[16] Whitman, J. B. *The phonological basis for the comparison of Japanese and Korean*. (Harvard University Ph.D. dissertation, 1985).
[17] Francis-Ratte, A. *Proto-Korean-Japanese: A new Reconstruction of the common origin of the Korean and Japanese Languages*. (The Ohio State University PhD dissertation, 2016)
[18] Park, J. Y. *Koetaysacen* [A comprehensive dictionary of Old Korean], 8 volumes. (Hakkobang, 2010)

| | | | | | | | |
|---|---|---|---|---|---|---|---|
| | 'to spin, twist (thread)' | 'twisted thread, weaving' | 'to twist (thread)' | | | | spin, to twist' is problematic" We did not reconstruct such a verb. SI 5: 38. PJ **pu* 'twisted thread, weaving' "pK is at best speculative" MK *pwuy-* 'twisted (bound adverb), MK *pwuk* 'shuttle (loom instrument)' (< **pu-k* twist-NMLZ), JJ *pi* 'shuttle' (< **pu-i* twist-NMLZ) all share a common element *pu- that is derivable from a common denominator 'to twist thread' |
| 42 | 10 pJK *ʌsa 'hemp' | *asa 'hemp' | *ʌsa-ma 'hemp' | | | | "alternative explanation suggested in SI5 that it is a Wanderwort is the correct one." Yes, room for agreement. |
| 43 | 11 pJK *mosi 'ramie (cloth)' | *mosi 'ramie (cloth)' | *mosi 'ramie cloth' | | | | "pJ *musi is not attested in Ryukyuan, which makes it likely that it is a loanword. MK mwòsì is unusual. Since other such nouns are often loanwords this might be the case of this word too." Either Japanese is a loanword from Korean, or Korean is a loanword from Japanese, but it cannot be to be both at the same time |

### 1.3. Regular sound correspondences

#### 1.3.1. Expert analysis

Complementing previous research on morphology[19, 20], our Article demonstrated the genealogical relatedness of Transeurasian languages based on 160 etymologies for basic

[19] Robbeets, M. Diachrony of verb morphology. Japanese and the Transeurasian languages (*Trends in Linguistics* **291**) (Berlin: De Gruyter Mouton, 2015).
[20] Robbeets, M. Verbal morphology in Transeurasian. In: Robbeets, M. and Savelyev, A. (eds.) *The Oxford Guide to the Transeurasian Languages*. Oxford: Oxford University Press, 511-521 doi: 10.1093/oso/9780198804628.003.0031, (2020b).

vocabulary as summarised in Table 1 and obeying regular sound correspondences for consonants and vowels prescribed by Tables 3 and 4 below. In line with the requirements of the historical comparative method, all compared sounds correspond regularly for each subsequent (consonant)-vowel-consonant string and in several cases the occasional root-final vowel corresponds regularly as well. Considering the shape of the entire phonological system of the proto-language, emerging as multiple proto-sounds are aligned, the overall phonemic inventory displays symmetrical arrangements of phonemes such as voice-voiceless distinction for stops, three medial cluster series, the occurrence of two liquid phonemes and a harmonic vowel system. Therefore, we argued that our dataset supports the genealogical relatedness of Transeurasian languages within the limits of the historical comparative method. Nevertheless, various details of our correspondences, including conditioned correlations that are restricted to certain phonological contexts remain open for future research.

**Table 3** Consonant correspondences between the Transeurasian languages

| | pJ | pK | pTg | pMo | pTk | pTEA |
|---|---|---|---|---|---|---|
| 1. | *p- | *p- | *p- | *p- | *b-/ *p- | *p- |
| 2 | *-p- | *-p- | *-p- | *-ɣ- | *-p- | *-p- |
| 3. | *p- / *w- | *p- | *b- | *b- | *b- | *b- |
| 4. | *-p-/*-w- | *-p- | *-b- | *-b-/ -ɣ- | *-b- | *-b- |
| 5. | *-np- | *-pC- | *-PC- | *-PC- | *-P(C)- | *-m$^{(P)}$T- |
| 6. | *-np- | *-Rp- | -RP- | *-RP- | *-RP- | *-Rp- |
| 7. | *t- | *t- | *t- | *t- | *t- | *t- |
| 8. | *-t- | *-t- | *-t- | *-t- | *-t- | *-t- |
| 9. | *t- /*y- | *t- (ci-) | *d- (ji-) | *d- (ji-) | *y- | *d- |
| 10. | *-t-/ *-y- | *-t-/-l- | *-d- (-ji-) | *-d- (-ji-) | *-d- | *-d- |
| 11. | *-nt- | *-c- | *-TC- | *-TC- | *-TC- | *-n$^{(T)}$K- |
| 12. | *-nt- | *-Rc- | *-RT- | *-RT- | *-RT- | *-Rt- |
| 13. | *k- | *k- | *k- | *k- | *k- | *k- |
| 14. | *-k- | *-k- (-h-) | *-k- | *-k- | *-k- | *-k- |
| 15. | *k- | *k- | *g- | *g- | *k- | *g- |
| 16. | *-k- | *-k- (-h-) | *-g- | *-g- | *-g- | *-g- |
| 17. | *-nk- | *-kC- | *-KC- | *-KC- | *-KC- | *-ŋ$^{(K)}$T- |
| 18. | *-nk- | *-Rk- | *-RK- | *-RK- | *-RK- | *-Rk- |
| 19. | *t- | *c- | *č- | *č- | *č- | *č- |
| 20. | *-t- | *-c- | *-č- | *-č- | *-č- | *-č- |
| 20b. | *-si | *-l(i)/ -c | *-l(č) | *-l(č) | *-l(č)~ -š | *-lč |
| 21. | *k- | *k-, h- | *x- | *k- | *k- | *x- |
| 22. | *-k- | *-k- | *-x- | *-g-~-k- | *-g-~-k- | *-x- |
| 23. | *s- | *s- | *s- | *s- | *s- | *s- |
| 24. | *-s- | *-s- | *-s- | *-s- | *-s- | *-s- |
| 25. | *m- | *m- | *m- | *m- | *b- | *m- |
| 26. | *-m- | *-m- | *-m- | *-m- | *-m- | *-m- |
| 27. | *n- | *n- | *n- | *n- | *y- | *n- |
| 28. | *-n- | *-n- | *-n- | *-n- | *-n- | *-n- |
| 29. | *-r- | *-l- | *-r- | *-r- | *-r- | *-r- |
| 30. | *-r- | *-l- | *-r- | *-r- | *-$r_2$- | *-r- |

| | | | | | | |
|---|---|---|---|---|---|---|
| 31. | *-r- | *-l- | *-l- | *-l- | *-l- | *-l- |

**Table 4** Vowel correspondences between the Transeurasian languages

| | OJ < pJ | MK < pK | pTg | pMo | pTk | pTEA |
|---|---|---|---|---|---|---|
| 32. | -a- < *-a- | -a- < *-a- | *-a- | *-a- | *-a- | *-a- |
| 32b. | *CaCa | *CʌCʌ | *CaCa | *CaCa | *CaC | *CaCa |
| 33. | -a- < *-a- | -e- < *-e- | *-e- | *-e- | *-e- | *-ə- |
| 34. | -o- < *-ə- | -e- < *-e- | *-e- | *-e- | *-e- | *-ə- |
| 35. | -o- <? *-o- | -wo- < *-o- | *-o- | *-o- | *-o- | *-ɔ- |
| 36. | -u- < *-o- / *-u- | -wo- < *-o- | *-o- | *-o- | *-o- | *-ɔ- |
| 37. | -o- < *-ɨ- | -u- < *-ɨ- | *-ö- | *-ö- | *-ö- | *-o- |
| 38. | -u- < *-u- | -wu- < *-u- | *-u- (gü) | *-ü- | *-ü- | *-u- |
| 39. | -u- < *-u- | -o- < *-ʌ- | *-u- | *-u- | *-u- /-ï- | *-ʊ- |
| 39b. | PaRu- < *PauRu- | *PʌRʌ- ~ *PɨRɨ- | *PuRV- | *PuRV- | *PuR- | *PʊRʊ- |
| 40. | -i- < *-i- | -i- < *-i- | *-i- | *-i- | *-i-/-ï- | *-i- |
| 40b | -i- < *-e- | -ye- < *-ia- | *-ia- | *-ia- | *-ia- | *-ia- |
| 40c | -e(1) - < *-e- | -ye- < *-ia- | *-ia- | *-ia- | *-ia- | *-ia- |
| 40 d | -o- < *-ə- | -e- < *-ə- | *-ü- | *-ö- | *-ö- | *-iu- |
| 41. | a- < *-a- | a- < *a- | *a- | *a- | *a- | *a- |
| 41b. | a- < *-a- | e- < *e- | *e- | *e- | *e- | *ə- |
| 42. | o- < *ə- | e- < *e- | *e- | *e- | *e- | *ə- |
| 43. | o- <? *o- | wo- < *o- | *o- | *o- | *o- | *ɔ- |
| 44. | o- < *ɨ- | ø < ? *ɨ- | *ö- | *ö- | *ö- | *o- |
| 45. | u- < *u- /*o- | wu- < *u- | *u- | *u- | *u- | *u- |
| 46. | i- < *i- | i- < *i- | *i- | *i- | *i- | *i- |
| 46b | i- < *e- | (y)e- < *ia- | *ia- | *ya- | *ya- | *ia- |
| 46c | ya- < *ia- | ye- < *ia- | *ia- | *ya- | *ya- | *ia- |
| 46d | o- < *-ə- | ye- <*iə | *ü- | *ö- | *ö- | *iu- |

One such correlation, for instance, is correspondence 36 in Table 4 above, relevant for the etymologies (17), (131) and (155) in Table 1 and (23) and (37) in Table 2. For the correspondence, Tian et al. suggest a mismatch because “the Ryukyuan forms unambiguously point to pJ **u* and not **o*”. The general trend is that pJ **o* becomes *u* in the Ryukyuan languages, but pJ **u* devoices and syllabifies, or induces other sound changes in Ryukyuan. The etymologies in Table 5 suggest that sound correspondence 36 includes both cases of OJ -*u*- deriving from pJ *-*o*- and from pJ *-*u*-. In the large majority of cases, the conditioning factor is the presence of a stem-final *-*u* in Proto-Japonic, which triggers the assimilation of *o to *u (pJ *o > u / _Cu). In Table 5, etymologies (1) SWELLFISH (EMJ *fuku* ‘swellfish’ < pJ **puku* < **poku*), (3) STEM (OJ *kuki*$_2$ ‘stalk, stem’< pJ **kuku*- <  **koku*-), (13) CHOOSE (OJ *sugur*- ‘choose, select’ < pJ **sunku*- < **sonku*-), (16) SPIN (OJ *tumu* 'spindle' < pJ **tumu* < **tomu*), (17) FOLLOW (OJ *tutuk*- ‘follow, continue’ < pJ **tutu(-)ka*- < pJ **totu(-)ka*-) and (18) WEAK (OJ *yuru*- ‘loose’ < pJ **yuru*- < **yoru*-) fulfil this conditioning factor.  Tian et al. complain that our analysis “contradicts [our] own stated sound correspondence principles”, but in reality, the alleged phonological mismatch is nothing but a conditioning factor, that does more to strengthen the Transeurasian hypothesis than to weaken it.

**Table 5** Etymologies reflecting OJ -u- < pJ *-o- or *-u- < pTEA *-ɔ-

| | TEA | Japonic | Koreanic | Tungusic | Mongolic | Turkic |
|---|---|---|---|---|---|---|
| | *-ɔ- | -u-<br>< *-o- or *-u- | -wo- < *-o- | *-o- | *-o- | *-o- |
| (1) | SWELLFISH | EMJ fuku ‘swellfish’<br>< pJ *puku ~ *poku | K pok ‘swellfish<br>< pK *pok | | | |
| (2) | DEEP | OJ puka- ‘be deep’<br>< pJ *puka- | MK ·pwok ‘inside’, phwuk ‘deeply’<br>< pK *pok ~ puk | | | |
| (3) | STEM | OJ kuki$_2$ ‘stalk, stem’<<br>pJ *kuku ~ koku | MK kwokwoli ‘stem of fruit, handle’<br>< *koko | | | |
| (4) | PEAK | OJ kuki$_1$ ‘peak’<br>< pJ *kuki ~ koki | MK kwo·kay ‘peak’<br>< *kokai | | | |

| | | | | | | |
|---|---|---|---|---|---|---|
| (5) | JOIN | OJ kusar- 'join' < pJ *kusa-ra- ~ *kosa-ra- | | | WMo qolba- 'join, unite' pMo *kolba- | OT koš-, 'join, unite' < pTk *koš- < *kolC- |
| (6) | ADD | OJ kupape2- 'add' < pJ *kopa- | MK kwop- 'double' < pK *kop- | | | |
| (7) | WHALE | OJ kudira, kusi 'whale' < pJ *kuti | MK kwolay 'whale' < *kolai | | | |
| (8) | BEAUTIFUL | OJ kupasi- 'be beautiful' < pJ *kopa- | MK kwo·po- 'be beautiful' < pK *kopʌ- | | | |
| (9) | VALLEY | OJ kura 'valley' < pJ *kura ~ *kora | MK ¨kwol ~ ¨kwo·loy 'valley, deep hole' < pK *kolʌ | Ma. golo 'valley, watershed' < pTg *golo | MMo qol 'river, river valley, centre' < pMo *gol | OT kol 'valley' < pTk *ko:l |
| (10) | PAINFUL | OJ kurusi- 'be painful' pJ *koru- | MK kwolwuW- 'be painful' pK *kolu- | | | |
| (11) | GET TOGETHER | OJ no muta 'together' MJ mutum- ~ mutub- 'be intimate' < pJ *motu- | MK mwot- 'gather, get together, assemble', mwut- 'put together, < pK *mot-~ mut- | | | |
| (12) | WET | OJ nure- 'be(come) wet' < pJ *nora- ~ nura- | | | WMo. nor- 'to be wet, soak' < pMo *nor- | |
| (13) | CHOOSE | OJ sugur- 'choose, select' < pJ *sonku- ~ sunku- | | | WMo.soŋgu- 'choose, select' < pMo *songu-~ sungu- | |
| (14) | SCOOP | OJ suk- 'dig up earth, scoop up' < pJ *suka- | | Evk. soko- 'scoop, ladle' < pTg *soka- | | |
| (15) | HAND/ARM | OJ sune 'leg, shin, shank' < pJ *sune | MK ·swon 'hand' < pK *son | | | |
| (16) | SPIN | OJ tumu 'spindle' < pJ *tumu | | Even tum- 'spin, wind, spool' < pTg *tomu- | WMo. tomu- ' twist or spin thread or rope' < pMo *tomu- | |
| (17) | FOLLOW | OJ tutuk- 'follow, continue' <pJ *tutu(-)ka- | MK cwoch- 'follow' < pK *cocɨ(-)k- | | | |

| | | | | | | |
|---|---|---|---|---|---|---|
| (18) | WEAK | OJ yuru-<br>‘loose’<br>pJ *yuru- | | | WMo. doru(i)<br>‘weak’<br>pMo *doru- | Tk. yor-<br>‘exhaust’<br>pTk *yor- |

(1) SWELLFISH

pK *pok* ‘swellfish, Takifugu chinensis’: K *pok* ‘swellfish, Takifugu chinensis’

pJ **puku* ~ **poku* ‘swellfish, Takifugu chinensis’, J *hugu* (2.5), EMJ *fuku* ‘swellfish’, Takifugu chinensis’

There are no Ryukyuan cognates that allow us to distinguish between pJ **o* and **u* in this case. Assuming final vowel loss in Korean, the correspondences are regular.

(2) DEEP

pJ **puka*- ‘deep’: J *hukai* (B ), OJ *puka*- ‘deep, thick, dense’, Amami *hukasari*; Shuri *hukasaN*; Hirara *fuka:* ‘depth’ Ishigaki *fukasa:ŋ*; Yonaguni *k’a:N* (B)

pK **pok* ~ **puk* ‘deep inside, midst’: The bound noun MK *·pwok* ‘inside’ in MK *·poy s* ·p*wok*, K *pay-kkop* ‘navel’; K *pwok-phan* ‘ the very midst’, Kog. OK **pok(sa)* ‘deep’. MK *phwuk* ‘deeply’ < **puk-ɨk* (deep-ADV) suggests an alternation between **u* and **o* in Koreanic.

(3) STEM

pJ **kuku* ~ **koku* ‘stem, stalk’: J *kuki* (2.3 ), OJ $kuki_2$ ~ *kuku*- ‘stem, stalk’ in OJ *kukutati* < *kuku* + tati ‘stand’ and OJ *kukumira* < *kuku* + *mira* ‘leek’, Amami *hunï*; Shuri *pizi*, Hirara *fuku* or $fuk^s$*ï*, Ishigaki *fukï*; Kohama *fukï*, Hateruma *fuki* ‘stem’< PR **kuki*, Shuri *guci* B ‘stem’ ?< pR *[ ] *-n- koki*

pK *koko* ‘stem’: MK *kwokwo·li* ‘calyx, stem of fruit, handle’, Mod. K *kwokci* ‘stem, stalk’,

Vovin[93] claims the Ryukyuan forms are loans from the mainland, “since the word is attested only in Northern and Central Ryukyus.” This is incorrect as there are clearly attestations in Yaeyama. Therefore, the proto-Japonic vowel should most probably be **u*, even if Shuri *guci* B ‘stem’ leaves some room for **o*.

(4) PEAK

pJ **kuki* ~ *koki* 'mountainside cave, peak': J (2.1) *kuki*, OJ $kuki_1$ 'hole in mountain cliff; path between two cliffs', EMJ *kuki* 'peak'
pK **kokai* 'peak': K *kokay*, MK *kwokay* 'peak'. This etymon may be cognate with MK *·kwoh* 'nose'

Note that in EMJ, the gloss of 鼽 'bridge of the nose' is given as *fana-kuki* as well as *fana-mine* (*Shinsenjikyō*), which supports a potential semantic relationship of *kuki* to 'nose'.[17] There are no Ryukyuan cognates that allow us to distinguish between pJ **o* and **u* in this case.

(5) JOIN
pJ **kosa-ra-* ~ **kusa-ra-* 'join': J *kusar-* 'link', *kusari* 'chain', Shuri *kusai* A 'chain'
pMo **kolba-* 'to connect, join, unite': MMo (SH) *qolba'ara-*, *qolbara-* 'to join, unite (intr)' MMo (MA) *qolba-*, WMo. *qolbo-*, *qolba-* Khal *xolbo-*, Bur *xolbo-*, Kalm. *xolv-*, Dag *xɔlb-*, EYu *χɔlβɔ-*, *χolbo-* Mgr Huzu *xolo:-*, Mgr Minhe *xulo-*
pTk *koš-* < *kolC-* 'to join, unite': OT *qoš-*, Karakhanid *qoš-*, Turkish *koš-,* Tatar *quš-*, Middle Turkic *qoš-*, Uzbek *qọš-*, Uighur *qoš-*, Sary-Yughur *qos-*, Az. *Goš-*, Turkmen *Goš-* 'to add', Khakassian *xos-*, Oyrat *qoš-*, Chuvash *xoš-*, Yakut *xohuj-*, Tuva *qoš-*, Tofalar *qo'š-*, Kirghiz *qoš-*, Kazakh *qos-*, Noghai *qos-*, Bashkir *quš-*, Balkar *qoš-*, Gagauz *qoš-*, Karaim *qoš-*, Karakalpak *qos-*, Salar *qoš-*, Kumyk *qoš-*.

As for the Japanese verb, there are no Ryukyuan cognates that allow us to distinguish between pJ **o* and **u*. The derived noun *kusai* 'chain' is attested in Shuri but given the poor distribution of this word in the Ryukyuan languages, this may be a loan from Middle Japanese.

(6) ADD
pJ **kopa-* 'add': J *kuwaeru* (?B) 'add', OJ $kupape_2$- (< **kupa-pa-Ci*), *kuwawaru* 'be added, grow', J *sukuu* 'build a nest' (*su* 'nest'), J *kuu*, OJ *kup-* 'make, assemble (a nest)', Amami *ku'uri*; Shuri *kuwe:yuN*; Ishigaki *kubairuŋ*; Yonaguni *p'umuN* 'add' (< pR **kowa-*)
pK **kwop-* 'double': K *kop* 'double, twofold', MK *kwop-* 'double, increase twofold',

The Ishigaki form suggests the reconstruction of **o* to proto-Japonic.

(7) WHALE

pJ **kuti~ kunti-ra* ‘whale’ (pJ *-*ra* plural) : OJ *kudira* ‘whale’, Fudoki OJ 久慈 *kusi* ‘whale’, Amami *kuzira*, Shuri *guzira* Hirara *fudza*, Ishigaki *futtsa*, Hateruma *guzïra*, Yonaguni *kudira* < PR **guzira* ? < *[ ] -*n- kuzira* (or **m[i]-kuzira*).
pK **kolai* ‘whale’: MK *kwolay* ‘whale’

The Fudoki attestation with *si* reflecting the known shift of *ti* > *si* in certain dialects of Old Japanese[17] supports the proposed morpheme boundary in Proto-Japonic. The voiced *-*z*- (<*-*nt*-) in Ryukyuan supports the reconstruction of **u* in Proto-Japonic. The Yonaguni form is likely a loan from Shuri, its absence of initial voicing suggesting that the Shuri model was originally unvoiced and that the voicing in Ryukyuan developed through prefixation of a nasal morpheme such as the genitive *-*n*- or the honorific **mi*-.

(8) BEAUTIFUL
pJ **kopa*- ‘beautiful’: J *kuwasii* B*,* OJ *kupasi*- ‘to be beautiful, detailed, accurate’ (OJ -*si*- suffix). Ishigaki *kumasa:ŋ* or *kuwassaŋ* ‘beautiful’ (< pR **kopa*-)
pK **kopʌ*- ‘be beautiful: MK *kwo·po*- ‘to be beautiful, attractive’

Given its congruent register and its closely overlapping semantics, OJ *kupasi*- ‘beautiful, detailed, accurate’ may be related with OJ *koɪp*- ‘love’ and OJ *koɪposi*- ‘lovely’, suggesting original **o* in Proto-Japonic. This is supported by the Ishigaki form, which would suggest reconstruction of **o* in Proto-Japonic. However, as cognates for this word are rare in Ryūkyūan and the Ishigaki form is from Miyara, the word is possibly a loan from Japanese.

(9) VALLEY
pJ **kura* ~ *kora* ‘valley’: OJ *kura* ‘valley’, J (2.2b) *kura* 'seat, saddle', Ishigaki *fufa*, Taketomi *fufa*, Hatoma *fufa* ‘saddle’ (< pR **kura*), Yonaguni *hura* A ‘saddle’.
pK **kolʌ* ‘valley’: MK ¨*kwol* ~ ¨*kwo·loy* ‘valley, deep hole’
pTg **gola* ‘valley’: only based on Ma. *golo* ‘valley, watershed’.
pMo **gol* ‘river, river valley, centre’: MMo. *qol* (SH), *Ɣol*, WMo. *Ɣoul*, Khal. *gol*, Bur. *gol*, Kalm. *Ɣol*, Ordos *gol*, Mogh. *Ɣo:l*, Dag. *gol*, *gole*, Dong. *go*n, S.-Yugh. *gol*, Mgr. g*or*,
pTk **ko:l* ‘valley’: Karakh. *qol*, Tk. *kol* (dial), Tkm. *Go:l*, MTk. *qol*, Uig. *qol* (dial), Tat. *qul*, Bash. *qul* (dial), Kirgh. *qol*, KBalk. *qol*, Kum. *qol*, SUig. *qol* ‘gutter’, Khak. *xol*, Tuva *xol*

In Old Japanese the word *kura* means 'valley'. The contemporary meaning 'saddle' represents a metaphoric extension. There are no Ryukyuan cognates meaning 'valley' but the words for 'saddle' in Yaeyama point to an original *u. As the Yonaguni word for 'saddle' is a loan from Shuri, we cannot completely exclude the reconstruction of **u* for this word.

(10) PAINFUL

pJ **koru*- 'be hard, painful': J *kurusii* B, OJ *kurusi*- 'to be painful, difficult, hard' (OJ -*si*- suffix), OJ *kurup*- 'go crazy', Amami *kucinasari*; Shuri *kurisaN* (B), *kucisaN* (B); Hirara *kucci:*; Ishigaki *kutsïsaŋ*; Taketomi *kususaŋ*;Yonaguni *kucisaN* (A) 'to be painful, difficult, hard' (< pR **kot*- ~ *kor*-).

pK **kolu*- 'be hard, painful': MK *kwolwuW*- 'to be painful, troublesome, hard', (MK -*W*- < pK *-·*po*- adjective suffix 'be characterized by'; cf. Martin 1992: 759)[95]

If the Ryukyuan proto-form **kot*- can be considered in alternation with **kor*- reflected in Shuri *kurisaN,* the Ryukyuan forms may be cognate with OJ *kurusi*- 'to be painful, difficult, hard'. If this is indeed the case, the Proto-Japonic vowel can be reconstructed as **o*.

(11) GET TOGETHER

pJ **motu*- 'join, get together': OJ *(no) muta* 'in the way of, as, together with' deverbal noun from the root underlying in J *mutumazii* 'intimate, affectionate, harmonious', OJ *mutu-goto* 'friendly words, OJ *mutubi*- 'be close, friendly', MJ *mutum*- ~ *mutub*- 'be intimate', Ishigaki *mutsïmassa:ŋ* (< pR **motu*-)

pK **mot*- ~ **mut*- 'gather, get together': MK *mwot*- 'gather, get together, assemble (intr.)' from which MK *mwo ·ton* 'all, every, each' is the verbal modifier and MK *mwo ·two* 'all' is the derived adverb. (Martin 1992: 696)[95]; MK *mwut*- 'put together, assemble (tr.)

The Ishigaki word is the only token for this word in Ryukyuan and suggests that proto-Japonic had **o* for this word. MK *mwut*- 'put together, assemble (tr.) signals an alternation between pK **o* and **u*.

(12) WET

pJ **nura*- ~ *nora*- 'to be(come) wet': J *nure*-, OJ *nure*- (A) 'to be(come) wet, damp; to come undone, loose', J *nuru* (A), OJ *nur*- 'to paint, smear', J *nuri* 'paint' ; Amami *neeruri*, Yoron (Amami) *nidituɴ* 'to be wet', Shuri (Okinawa) *ndir*- (A) 'to get wet', Shuri *'NdiyuN, ɴditooɴ*

'to be wet', Hatoma (Yaeyama) *zoori bee* 'to be wet', Hateruma *duriŋ*; Yonaguni *NgaruN* ‘to be wet’ (< pR **nure-*); Irabu (Miyako) *nil*, Hirara *nuï*, Ishigaki (Yaeyama) *nu:rï* ‘paint’, *nuruŋ* ‘to paint, smear’, Hateruma (Yaeyama) *nuruŋ*, Kohama (Yaeyama) *no:ruŋ,*
pMo **nor-* 'to be(come) wet': WMo. *nor-* 'to become wet, soaked, drenched, damp, moist (intr.)', *norɣa-* 'to wet, moisten, soak (tr.)', MMo. *nur-* 'to be wet, soak', *norma* 'moistened', Khal. *nor-* 'to get wet', Bur. *noro-* '1 to become wet, soaked', Kalm. *nor-* 'to become wet', *norɣa:-* 'to make wet', *norɣu:(n), noru:* 'wet', Ordos *nor-* '1', Dgx. *noro-* ~ *nuru-* ~ *noru-* '1', Dag. *noir-* '1', *noirga*:- 'to make wet', Mgr. *no:ri-* ~ *nori-* '1'

The Japanese meanings ‘to paint, smear’ and ‘to be(come) wet’ go back to causative and anticausative formations of the same etymon. Ryukyuan cognates of ‘be(come) wet’ all indicate that **u* should be reconstructed to the Proto-Japonic. However, Shuri has *nur-* for ‘to paint, smear’, rather than **ndu-* that the reflex of *nureru* leads us to expect. Moreover, the Kohama *no:ruŋ* is odd. In many dialects in the Ryukyus **no* and **nu* have merged as *nu*. Therefore, **o* cannot be completely excluded as the Proto-Japonic vowel.

(13) CHOOSE
pJ **sunku-ra-* ~ **sonku-ra-* ‘choose, select’: J *suguru* (?A/B) ‘choose, select’, Shuri *su-yuN*; Ishigaki *suguruŋ* or *su:ruŋ*.
pMo **songu-* ~ **sungu-* ‘to choose, select, elect’: MMo (SH) s*o’ongqu-*, *songqu-*, MMo. (MA) *sonqu-*, WMo. s*ongɣo-*, Khal. s*ongo-*, Ordos *suŋġu-*, Bur h*unga-*, Kalm *suŋɣ-*, s*oŋgăχă*, EYu s*ɔŋġə-*, *soŋġo-*, Mgr. Huzu s*ɔŋġʊ-*, Kgj c*uŋgu-*, Dgx *suŋġu-*.

Since the merger is complete in Yaeyama for *so vs *su and we do not have a Yonaguni cognate, we are unable to distinguish between pJ **o* and **u* for this word. In Mongolic, Buriat and Kalmyck support the reconstruction of pMo **u* in alternation with **o*.

(14) SCOOP
pJ **soku-* ~ *suku-* ‘scoop’: J *suku* (?A ), OJ *suk-* ‘dig up earth, scoop, dip, ladle', J *sukuu* (A), OJ *sukup-* ‘rescue, scoop’, Shuri *sukur-* A ‘rescue, scoop’, Shuri *ficuN*, Ishigaki *sïkuŋ* ‘dig up earth, scoop, dip, ladle' (< pR **sik-*)
pTg **soka-* ‘to scoop, ladle’: Evenki *soko-*, Even: *hụqụ-*, Negidal *soxo-*, Ulcha *su:-su-* Orok *so:-,* Nanai *so:-lo-*, Oroch *soko-*, Ud. *so`-lo-*

The Ryukyuan vocalism points to an original *ɨ, while the Mainland Japanese forms leave room for the reconstruction of both *u and o.

(15) ARM/ HAND

pJ **sune* 'lower limb, leg': J *sune* (2.3), OJ *sune* ' leg, shin, shank'; Asama (Amami) *sɨnɨɨ*, Naze (Amami) *sïnï*, Shuri (Okinawa) *ʃini*, Hirara (Miyako) *karasïni*, Ishigaki (Yaeyama) *sïni* (B), Yonaguni *ccini* (B) (< pR **sune* 'leg, shin, shank')

pK **son* 'upper limb, arm, hand': K *son*, MK *·swon* 'hand'

Based on the Yonaguni data, the Proto-Japonic vowel should be *u. Since the quality of the mid vowel is not distinguished after *n* in Old Japanese, we could be dealing with either $e_1$ (<**i(C)a*, **i(C)ə*, **i(C)ɨ*, **e* ) or $e_2$ (< **a(C)i* /**ə(C)i*). However, as there are no apophonic alternations attested for this root and since nominal roots rarely consist of more than two syllables, preference is given to the reconstruction of pJ **sune*. The Ryukyuan forms support this reconstruction.

(16) SPIN

pJ **tumu* 'spindle': J *tumu* (2.4), OJ *tumu* 'spindle', J *tumug-* (B), OJ *tumug-* 'to spin, make into yarn', Shuri (Okinawa) *çing-* (B) 'to spin', Yamatohama (Amami) *cïɴg-* 'spin a thread with a spindle', Asama (Amami) *cïɴg-* 'spin', Yonamine (Amami) *ciɴg-* 'spin', Irabu (Miyako) *tsïmk-*, Ishigaki (Yaeyama) *tsïn-*

pTg **tom(u)- > tumu-* 'to spin' (pTg *-*ku* ~ *-*ko* deverbal instrumental noun suffix, e.g. Na. *xado-* 'to mow' → *xadoko* 'scythe'): Even *tum-* '1 to spin, wind, coil, spool, wrap', *tumenŋe* 'thread wind around a bobbin', *tomqo-* 'to spin strings for threads; to spin threads', *tomqon* 'spinning of strings; yarn'; Evk. *tum-* ' 1', *tomko* 'a thread'; *tomko-* 'to spin threads; to tie with a thread'; Neg. *tum-* '1', *tumu* 'string of thread, parcel, roll', *tomko* 'thread', *tomko-* 'to spin threads'; Solon *tum-* 'to bind, knit, string together, tie', *toŋxo-* 'to spin threads'; Oroch *tumu-* '1', *tompo* 'a sinew or nettle thread', *tompo-* 'to spin threads; to weave a net'; Olcha *tumu-* '1', *toŋpo* 'short threads of nettle or hemp', *toŋpo-* 'to spin threads'; Orok *tumu-* '1', *toqpo-* ~ *topqo-* 'to spin threads, rope, cord'; *toqpo* ~ *topqo* 'thread; rope'; Na. *tumu-* '1', *tompo-* 'to spin threads of fishskin', *tompo* 'threads made of fishskin'; Ud. *tompo-* 'to spin threads, ropes' pMo **tomu-* ~ **tamu-* 'to spin': WMo. *tomu-*, *tamu-* '1 to twist or spin thread or rope', MMo. *tamu-* ~ *toma-* ~ *tomu-* ~ *doma-* '1', Khal. *tam-* ~ *tom-* '1', Bur. *tomo-* '1', Ordos *tamu-* '1', Kalm. *tom-* ~ *töm-* 'to twist, twine; to string together (rope), make rope (by turning horse hair between the hands)', Eastern Yugur *tomu-* ~ *tɔmɔ-* ~ *tomə-* '1', Dgx. *tomu-* '1', Bao. *toməl-* '1', Mgr. *tomu-* ~ *tamu-* '1'

As Tian et al. correctly observe, the Ryukyuan forms unambiguously point to Proto-Japonic **u* and not **o* (e.g., Shu. *çing-* and not †*tung-*) but the stem-final *-*u* fulfils the conditioning factor pJ *o > u / _Cu. The Tungusic reconstruction displays a vowel alternation between **tom(u)-* and **tumu-* 'to spin'. Given the preservation of **tom(u)-* in derived nouns with the instrumental suffix and re-verbalizations thereof, I assume that **tom(u)-* assimilated to **tumu-*, independent of a similar development in Japonic. In Mongolic we find a vowel alternation between **tomu-* ~ **tamu-* 'to spin', reminiscent of the alternation between pMo **dalan* 'seventy' and **dolaan* 'seven'[27].

(17) FOLLOW
pJ **tutu(-)ka-* ~ **tuntu(-)ka-* ‘follow, continue’, OJ *tuduk-* ~ *tutuk-* ‘follow, continue’, J *tuzuku* ‘continue, be contiguous’, *tuzukeru* ‘continue’, Amami *cïziki*; Shrui *cizicuN*, *çizik-* B Ishigaki *tsïdzïkuŋ*; Yonaguni *c’idikiruN* (< pR **cuc-*)
pK**cocɨ(-)k-* ‘follow’: MK *cwoch-* ‘follow’

The Ryukyuan high, back vowel is hard to tease out but it is likely that Proto-Japonic had **u* here. The unusual trisyllabic shape in Japonic and Koreanic suggests that we are dealing with a case of the common inchoative suffix *-*kA*.[19]

(18) WEAK
pJ **yuru-* ‘to be loose’: J *yurui* B‘to be loose, lax, slow’, OJ *yuru* ‘loose, slack, lenient’, OJ *yura-* ‘to be loose, lenient, generous’, OJ *yurum-*, J *yurumu* ‘to loosen’, Amami ’*yorasari*; Shuri ’*yurusaN*; Hirara *yurumunu*, Irabu *yuru:*; Ishigaki *yuruho:nu*, *yurusa*, Yonaguni *dura:N* ‘loose’ (< pR **yuru-*).
Manchu *duru-* ‘to become worn out, old’ may be a borrowing from Mongolic
pMo **doru-* ‘to be weak, feeble’: WMo. *doru* ‘weak, incompetent’, *dorui* ‘weak, feeble, emaciated’ (cf. pMo *-*i* deverbal noun suffix; Section 8.3.4), Khal. *dor*, *doroy*, Bur. *doroy*, Kalm. *doru:*, Mgr. *duri*:

pTk *yor- 'to exhaust': MT *yorul-* '1 wear out, tire, exhaust', *yorγun* '2 tired, exhausted', Tk. *yor-* '1 wear out, tire, exhaust', *yorγun* '2 tired', Gag. *yorul-* '1', *yorγun* '2', Az. *yor-* '1', *yorul-* '1 ', *joryun* '2', Tkm. *yor-* '1', *yorγun* '2', Krm. *yorul-* '1', *yorγun* '2', Kumyk *yorul-* '1'.

All Ryukyuan data, except Amami point to the reconstruction of **u* in Proto-Japonic. Amami indicates **o* but our suspicion is that the Amami form is a diphthong. Therefore, **u* should be reconstructed here. The Manchu adjective is likely to have been copied from Mongolian, because the distribution in Tungusic is restricted to Manchu and the vowel correspondence is irregular. If this analysis is correct, the Manchu form, being a verbal property word, should have been copied from a verbally encoded model. This observation, together with the attestation of forms of the shape *dorui* in Mongolic, which are probably deverbal derivations with the suffix *-*i*, indicates that the Mongolic proto-form is a verbal adjective. The corresponding Turkic form is an original transitive verb.

Tian et al. note that a few columns are missing in our correspondence tables: "Neither SI2 Table 3.5 nor SI2 Table 3.6 for Tungusic sound correspondences includes Hezhe" and "Orq. is not included at all." Below we added the missing columns in red.

**Table 6** Reconstruction of the basic consonant inventory of Proto-Tungusic

| pTg | Jur. | Ma. | Sibe | Oroch | Ud. | Na. | Na. Bikin | Hezhe | Orok | Olcha | Even | Sol. | Oroqen | Evk. | Neg. |
|---|---|---|---|---|---|---|---|---|---|---|---|---|---|---|---|
| *p- | f- | f- | f- | x- | x- | p- | f- x- | f- x- | p- | p- | h- | ø- | ø- | h- | x- |
| *-p- | f | f b ø | v ø | p w ø | p f w | p | f | f | p | p | b w ø | p w g ø | b w | p b w ø | p w |
| *b- | b- | b- | b- | b- | b- | b- | b- | b- | b- | b- | b- | b- | b- | b- | b- |
| *-b- | b w ø | b f w ø | v ø | b w ø | b w ø | b w ø | w ø | w ø | b w ø | b w ø | b w ø | b p w ø | w ø g | w ø | w ø |
| *t- | t- | t- | t- | t- | t- | t- | t- | t- | t- | t- | t- | t- | t- | t- | t- |
| *-t- | t | t | t s | t | t | t | t | t | t | t | t | t | t | t | t |
| *d-<br>*ji- | d-<br>ʤi- | d-<br>ʤi- | d-<br>ʤi- | d- | d- | d-<br>ʤi- | d-<br>ʤi- | d-<br>ʤi- | d-<br>ʤi- | d- | d- | d- | d- | d- | d- |
| *-d-<br>*-ji- | d<br>ʤi | d<br>ʤi | d<br>ʤi | d | d | d<br>ʤi | d<br>ʤi | d<br>ʤi | d<br>ʤi | d | d | d | d | d | d |
| *k- | x- | x- | x- | k- ø- | k- ø- | k- ø- | k- ø- | k- x- | k- ø- | k- ø- | k- ø- | x- ø- | k- | k- ø- | k- ø- |
| *-k- | k x ø | k x | k ɣ | k ø | k x ɣ ø | k ɣ ø | k ɣ ø | k x | k ø | k ɣ ø | k | k x | k x | k x | k x |
| *g- | g- | g- | g- | g- ŋ- | g- ŋ- | g- ø- | g- ø- | g- ø- | g- ŋ- | g- ŋ- | g- ŋ- | g- n- | g- | g- ŋ- | g- ŋ |
| *-g- | ɣ w ø | ɣ w y ø | ø | ɣ w y ø | ɣ w y ø | ɣ w y ø | y ø | ø | ɣ w y ø | ɣ w y ø | ɣ y | ɣ ø | ɣ ø y | ɣ | ɣ y w |
| *č- | č- | č- | č- | č- | č- | č- | č- | č- | č- > t- | č- > t- | č- | s- | č- | č- | č- |
| *-č- | č | č | č | č | s | č | č s | č | č > t | č | č | š | č | č | č |

| | | | | | | | | | | | | | | | |
|---|---|---|---|---|---|---|---|---|---|---|---|---|---|---|---|
| *x- | w-<br>ø- | w-<br>ø- | v- ø- | x- ø- | w-<br>ø- | x- s- | x- s- | ø- | x- s- | x- s- | ø- | ø- | ø- | ø- | ø- |
| *-x- | x | x | x k<br>ɣ | k | ø | x ø | x k | x k | x ø | x ø | k | x | k x | k | k x |
| *s- | s- | s- | s- | s- | s- | s- | s- | s- | s- | s- | s- | s- | s- | s- | s- |
| *-s- | s | s | s | s | s h ø | s | s | s | s | s | s | s | s x | x | s |
| *m- | m- | m- | m- | m- | m- | m-<br>ŋ- | m- | m- | m- | m-<br>ŋ- | m- | m- | m- | m- | m- |
| *-m- | m | m | m | m | m | m | m | m | m | m | m | m | m | m | m |
| *n- | n- | n- | n- | n- | n- | n- l- | n- l- | n- l- | n- l- | n- l- | n- | n- | n- | n- l | n- |
| *-n- | n | n | n | n | n | n | n | n | n | n | n | n | n | n | n |
| *-r- | r | r | r | y ø | y ø | r | r | r ø | r | r | r | r | r | r | y ø |
| *-l- | l | l | l | l | l | l | l n | l | l | l | l | l | l | l | l |

**Table 7** Reconstruction of the basic vowel inventory of Proto-Tungusic

| pTg | Jur. | Ma. | Sibe | Oro<br>ch | Ud. | Na. | Na.<br>Bikin | Hez<br>he | Oro<br>k | Olch<br>a | Eve<br>n | Sol. | Oro<br>qen | Evk. | Neg. |
|---|---|---|---|---|---|---|---|---|---|---|---|---|---|---|---|
| *a | a | a | a | a | a | a | a | a | a | a | a | a | a | a | a |
| *e | e | e | e | e | e | e | e | e | e | e | e | e | e | e | e |
| *o | o | o | o | o | o | o | o | o | o | o | o | o | o | o | o |
| *ö | u | u | u | o u | o | u | u | u | o u | o u | o | u | u | u | u |
| *u | u | u | u | u | u | u | u | u | u | u | u ö | u ö | u ö | u ö | u ö |
| *ü | u ei | u ei | u i | i | i u | u o | i o | i u o | i u | i u o | i | i | i | i u | i o |
| *i | i | i | i | i | i | i i̥ | i i̥ | i | i | i i̥ | i | i | i i̥ | i i̥ | i i̥ |
| *ia | ie<br>$^{i}$a | iya<br>ai $^{i}$a | ia a | $^{i}$æ<br>ei | $^{i}$æ æ<br>a | $^{i}$a ea | $^{i}$æ | ia $^{i}$a | i̥: | i̥: | ia | i̥: | i̥: | i̥: | i̥: |

#### 1.3.2. Automatically computed judgments

Applying automatically computed judgments to a small fraction of our dataset, Tian et al. conclude that "[t]he low number of cognate sets that follow the criteria of sound regularity and distribution over three or more families seriously undermines the Transeurasian hypothesis."

First, this conclusion is misguided because the computation is restricted to only 46 cognate sets out of the 160 basic vocabulary sets proposed in our Article and summarized in Table 1 above. The motivation for reducing our body of evidence to such an extent results from misinterpreting a criterion we applied for distinguishing inherited from borrowed verbs in the subsistence vocabulary, notably "Phonologically regular correspondences of bare verb roots that are well distributed in 3 or more subfamilies are likely to be inherited" (Robbeets et al. 2021, SI 5: 2). As sound correspondences are established within the basic vocabulary, which is relatively resistant to borrowing by itself, it is completely arbitrary to introduce this requirement here. The historical-comparative method by no means prescribes that cognates should be attested in at least 60% of the daughter languages involved. As it is generally known that the algorithm fails when the datasets are too small[21], it is misleading to apply the automatic cognate detection methods to only a small fraction of our evidence.

Second, current research[7] has suggested several explanations why automated cognate detection methods still perform worse than expert-coded ones. One issue is that morphologically complex forms cannot be sufficiently detected by the algorithm. This can be illustrated by the item 55_7 **da-k-*,**da-n-* burn (v.) reconstructed as [*d a ?] in Tian et al.'s Table 1.3. (Tian et al. SI: 4). As suggested in our Article (SI 2: 25), the attestation of Old Turkic (OT) *yal-* 'to blaze, burn, shine (intr.)' and OT *yan-* 'to burn, blaze up (intr.)' in addition to OT (Karakh.) *yak-* '1 to ignite, burn (tr.)' indicates that these verbs are morphologically complex. The underlying verb being pTk **ya-* 'to burn (tr.)' and the Old Turkic verbs representing derivations with a passive suffix pTk *-*(X)l-* (Erdal 1991: 651-693), an anticausative pTk *-*(X)n-* (Erdal 1991: 584-638) and an inchoative pTk *-*(X)k-* (Erdal 1991: 645-650), respectively.[94] Interestingly, the anticausative and inchoative suffixes can be traced back to the proto-Transeurasian ancestor: *-*nA-* processive and *-*gA-* inchoative.[5] Therefore, our Article pointed to the possibility that the Japanese and Korean forms, OJ *tak-* 'to burn, boil, cook (tr.)' (< pJ **tak-* 'to burn, boil (tr.)') and MK *·tho-* 'to burn, be on fire (intr.)', MK *ta·hi-* 'to make (fire), heat (with fire) (tr.)' (< pK **takʌ-* 'to burn (intr.)'), may have inherited the Transeurasian complex inchoative form **da-ga-* 'burn-INCHOATIVE'. As such, we could reconstruct an alternation in Transeurasian between **da-ga-* 'burn-INCHOATIVE' and **da-na-* 'burn-PROCESSIVE'. However, such alternating morphologically complex forms cannot be detected by the algorithm.

Another issue is that automated cognate detection methods tend to overlook morphophonological alternation such as the *a* ~ *e* alternation in the demonstrative pronouns. This can be illustrated by the item 169_27 **ta-*, *te-* 'that' reconstructed as [t ? -] in Tian et al.'s Table 1.3. (Tian et al. SI: 4). The original Transeurasian distinction in demonstrative pronouns was probably a two-way proximal-distal distinction. As we find frequent vowel alternations between *a* and *e* in these pronouns, we cannot exclude that the alternation originally marked an opposition between proximal and distal forms. For instance, the demonstratives Proto-Transeurasian **e-* and **a-* in our Article (SI 5: 59-60) may suggest such a distinction. Similarly, the occurrence of demonstratives of the shape **te-* as well as **ta-* (e.g., MMo. *tere* 'that', Manchu *tere* 'that', Evk. *tar(i)* 'that', Even *tar* 'that', Orok *tari* 'that',

[21] Rama, T. et al. Are automatic methods for cognate detection good enough for phylogenetic reconstruction in historical linguistics? Proceedings of NAACL-HLT **2018**, 393–400 (2018).

Solon *tari* ‘that’, Negidal *taj* ‘that’, Oroqen *tare* ‘that’) may represent relics of this original vocalic distinction. Table 8 reconstructs this vocalic symmetry in demonstrative system. Unfortunately, such morphophonological complexities cannot be detected by automatically computed judgments.

**Table 8**. The comparison and reconstruction of demonstrative pronouns in Transeurasian

| Axis | MRCA | *Proto-Turkic* | *Proto-Mongolic* | *Proto-Tungusic* | *Proto-Koreanic* | *Proto-Japonic* |
|---|---|---|---|---|---|---|
| proximal | pTEA **e-* | ***e-**<br>Chuv. *agə̑*<br>PROX.DEM<br>Chuv. *avə̑*<br>DIST.DEM<br>(Savelyev 2021) | ***e-**<br>MMo. *ene*<br>PROX.DEM | ***e-**<br>Ma. *ere*<br>PROX.DEM | | ***ə-**<br>OJ *ore, ono₂*<br>‘self’<br>*Shuri ʔu-*<br>MES.DEM |
| distal | pTEA **a-* | ***a**<br>OT *an-ča*<br>DIST.DEM-way ‘in that (previous) way’ | ***a**<br>MMo. *a-nu*<br>3PL-POSS. | | | ***a**<br>OJ *a-*<br>DIST.DEM |
| proximal | pA **te-* | | ***te-**<br>MMo. *tere*<br>DIST.DEM | ***te-**<br>Ma. *tere*<br>DIST.DEM | | |
| distal | pA **ta-* | | | ***ta-**<br>Evk. *tari*<br>DIST.DEM | | |

Finally, assuming that it is not due to a human error made by Tian et al., the algorithm used by the authors appears to attribute reflexes of the same etymon with different meanings to different ancestral roots. This can be illustrated by the items 152_24 **siara*- white reconstructed as [*s ia: r] and 253_50 **siara*- yellow reconstructed as [? ? ?] in Tian et al.’s Table 1.3. (Tian et al. SI: 4). In reality, both reconstructions proposed by the algorithm go back to the same root, pTEA **siara*- ‘be white’, discussed in our Article (SI 2: 52-54) and listed in Table 1 (125) above. Surprisingly, especially given the fact that it concerns one and the same etymology, the algorithm suggested an impeccable reconstruction in the first case but completely failed in the latter. Whereas the Japonic, Koreanic, Common Turkic and most Tungusic reflexes preserved the meaning ‘be white’, the Western Old Turkic cognates developed the meaning ‘be yellow’, which is also found in Mongolic. The automated cognate detection did thus not take semantic shift into account, which can have disastrous consequences, especially when establishing cognacy at greater time depths.

Given these shortcomings, the search for genealogically related words in different languages is still regarded as only partially automatable today. It is thus hardly surprising that the method yields poor results in establishing sound correspondences between the

Transeurasian languages, not least because it was only applied to a small fraction of our evidence.

#### 1.3.3. Suprasegmental correspondences

A recurrent objection made by Tian et al. is that "long vowels are attested and reconstructed in Mongolic, Tungusic and Turkic, but they do not appear anywhere in the correspondence tables and remain unaccounted for" and that "[t]onal distinctions in Japonic are also completely ignored."

However, a basic requirement of the historical comparative method is that the segmentals — consonants and vowels— should correspond. Suprasegmental correspondences of tone, accent and vowel length are expected to strengthen a hypothesis of linguistic relationship but are not regarded as a *conditio sine qua non*. For instance, most scholars[22,23,24] accepted that the Baltic and Slavic languages descend from a common Balto-Slavic ancestral language, even before details of comparative Balto-Slavic accentology[25,26] were revealed.

As the reconstruction of vowel length in certain daughter branches such as Mongolic is still a highly controversial issue today[27], the comparison to manifestations in the other Transeurasian languages is very premature. Nevertheless, certain conditioning factors relating to vowel length were explained in our Article (SI 2) but they were overlooked by Tian et al., as well as by their automatic cognate detection. For instance, for 'blow (v.)' and 'woods (n.)', Tian et al. complain that "[n]o explanation is provided for the irregular absence of the expected final *r in pTg" or that "vowel length in Tungusic is unaccounted for". However, an explanation that accounts for the Tungusic vowel length and absence of the expected final *r is provided in our Article (SI 2: 3, 23, 41, 44-45). Notably, open monosyllabic words with length in Tungusic are found to correspond to disyllabic forms with a liquid onset in the second syllable in the other Transeurasian languages. In addition to 79 'to blow' and 80 'woods', the conditioning factor is also mentioned for the comparative sets under 4 'water', 53 'to give', 102 'to sit' and 118 'year'; see Table 1 (03), (50), (71), (73), (97), (118).

---

22 Beekes, R. Comparative Indo-European Linguistics.An Introduction. (John Benjamins, 1995)
23 Petit, D. Les langues baltiques et la question balto-slave. Histoire Épistémologie Langage. **26** (2): 7–41. (2004). doi:10.3406/hel.2004.2092.
24 Mallory, J. P. &Adams, D. Q. The Oxford Introduction to Proto-Indo-European and the Proto-Indo-European World. (Oxford University Press, 2006).
25 Olander, T. Balto-Slavic Accentual Mobility. (De Gruyter 2009)
26 Kortlandt, F. Balto-Slavic accentuation revisited. Wiener Slavistisches Jahrbuch 56: 61-81 (2010).
27 Nugteren, H. Qinghai-Gansu languages. (Leiden University dissertation, 2011): 134-135

Carrying out an in-depth investigation of the correlations of tonal distinctions in Japonic was beyond the interdisciplinary purpose of our Article. However, previous research has proposed a correlation between register in proto-Japonic on the one hand and voicing and vowel length in pre-proto-Japonic on the other [5,28,29,30,31,32,33]. With Middle Japanese prosodic patterns Type A corresponding to a high initial tone and type B to a low initial tone, the correlation is summarized in Table 9. In the basic vocabulary, for instance, we find that the Japanese verbs and verbal adjectives with low initial register under 36 'hit/beat (v.)', 77 'thick', 170 'cut (v.)', 186 'rise (v.)' and 189 'break (v.)' all derive from Proto-Transeurasian reconstructions with an initial voiced consonant, see Table 1 (34), (67), (139), (149) and (153).

**Table 9** Correlation between voicing and vowel length in Pre-Proto-Japonic and register in Proto-Japonic and Middle Japanese.

| pre-pJ | voiced initial | voiceless initial |
|---|---|---|
| short V (CVCV) | pJ low register (B) | pJ high register (A) |
| long V (CV:CV) | pJ low register (B) | pJ low register (B) |

Given these indications of suprasegmental correspondences between the Transeurasian languages, there is no doubt that this topic would provide a fruitful basis upon which to build future research.

#### 1.4. Number of cognates

Based on a questionable count of cognates, Tian et al. conclude that "The core evidence presented is thus inadequate, and the Transeurasian hypothesis remains unwarranted."
Table 1 summarizes the core evidence presented in support of the Transeurasian hypothesis advanced in our Article. The coloured cells mark cognates that are criticized by Tian et al., while uncoloured cells contain cognates that remain uncritized. Table 10 counts the number

[28] Murayama, S. in Robbeets, M. (ed.) Transeurasian linguistics, vol. 4 (Routledge, 2017)
[29] Kortlandt, F. The origin of the Japanese and Korean accent Systems. *Acta Linguistica Hafniensia* **26**, 57-65 (1993).
[30] Starostin, S. A. On vowel length and prosody in Altaic languages. *Moskovskij Lingvističeskij Žurnal* [*Moscow Linguistic Journal*] **1**, 191-235 (1993).
[31] Vovin, A. The origin of register in Japanese and the Altaic theory. *Japanese/Korean Linguistics* **6**, 113–133 (1995)
[32] Vovin, A. in Frellesvig, B. & Whitman, J. (eds.) *Proto-Japanese: Issues and prospects* (Benjamins, 2008).
[33] Robbeets, M. Review of John Whitman & Frellesvig, Bjarke (eds.) 2008, *Journal of language relationship* **2**, 144–150 (2009).

of cognates and calculates the proportion of criticized cognates. Interestingly, only 9% of our cognates are criticized in Tian et al.'s "detailed qualitative analysis". In terms of etymologies, i.e. cognate sets, we count that out of 160 etymologies, 26 etymologies contain criticized cognates while 134 remain uncriticized, so only 16% of our etymologies are criticized.

The logical fallacy lurking behind Tian et al.'s refutation of the Transeurasian hypothesis, on the basis of a small number of unfavorable etymologies, while ignoring favorable ones is known as "observational selection". Naturally, it would be naïve to assert that in this way the Transeurasian hypothesis might be refuted. That would be the same as if, for example, one would point to the numerous errors in the Indo-European dictionary of Walde-Pokorny[1] and believe that the Indo-European hypothesis would thereby be refuted.

A question that arises is whether all cognates that are not rejected by the authors are actually accepted. If this is indeed the case, we have probably reached common ground for the acceptance of the Transeurasian hypothesis.

**Table 10**. The number of cognates proposed in our Article along with the proportion of criticized cognates by Tian et al. Count based on colour markings in Table 1.

| | Number of cognates | Number of criticized cognates | Number of uncriticized cognates |
|---|---|---|---|
| Attested in 2 subgroups | 204 | 5 | 199 |
| Attested in 3 subgroups | 117 | 15 | 102 |
| Attested in 4 subgroups | 56 | 14 | 42 |
| Attested in 5 subgroups | 25 | 4 | 21 |
| TOTAL | 402 | 38 (0.09) | 364 (0.91) |

The numbers advanced by Tian et al. are highly misleading. They count "3166 cognate sets listed by Robbeets et al. in support of the Transeurasian hypothesis" while our article lists only 203 cognate sets in support of the Transeurasian hypothesis, i.e. 160 cognate sets in the basic vocabulary (Table 1) and 43 in the vocabulary relating to subsistence (Table 2). The 3166 rows in the dataset underlying our Bayesian approach are cognate alignments that do not serve to establish the Transeurasian hypothesis but rather to determine and quantify the likelihood of the branching of our tree.

Their surprise that "only 317 are found in more than one language family" is unfounded. It is in line with expectations because it is common in Bayesian cognate alignments for the bulk of the cognates to be singletons.

Their claim that "only 50 of these are shared by more than two families and thus could be taken as evidence for the Transeurasian hypothesis" is wrong. In reality, 58 cognate sets are shared by more than two families; see Table 1 and corrected numbers in Table 11 in red. But more misleading than the miscount is their requirement that cognates can only be accepted on condition that they are shared by more than two families. The historical-comparative method by no means prescribes that cognates should be attested in at least 60% of the daughter languages involved.

Finally, they eliminate the remaining evidence by asserting that "[a] systematic computer-assisted analysis of the remaining cognate sets reveals that only 17 etymologies follow the criteria for the identification of regular sound correspondences" As noted in Section 1.3.2, computer-assisted analyses of cognate sets are known to perform worse than expert-coded ones. In addition, the analyses are only applied to a small fraction of the evidence and reflect numerous inconsistencies, as illustrated above.

**Table 11** Distribution of cognates across a collection of datasets (adapted from Tian et al.)

| | **Transeurasian** Robbeets et al. 2021 | **Sino-Tibetan** Sagart et al. 2019 | **Sino-Tibetan** Zhang et al. 2019 | **Indo-European** Dunn & Tresoldi 2021 |
|---|---|---|---|---|
| Major subgroups | 5 | 7 | 10 | 10 |
| Concepts | 254 | 250 | 100 | 207 |
| Cognates in 60% of subgroups | 38 (0.15)<br>39 (0.15) | 47 (0.19) | 19 (0.19) | 52 (0.25) |
| Cognates in 80% of subgroups | 5 (0.02)<br>14 (0.05) | 23 (0.09) | 11 (0.11) | 32 (0.15) |
| Cognates in 100% of subgroups | 2 (0.01)<br>5 (0.02) | 19 (0.08) | 7 (0.07) | 12 (0.06) |
| Age | 9200 BP | 7200 BP | 5900 BP | 7500 BP |
| Surviving languages | Ca. 75 | Ca. 500 | Ca. 500 | Ca. 450 |
| Earliest Historical attestation | Ca. 1300 BP | Ca. 3000 BP | Ca. 3000 BP | Ca. 3000 BP |

Tian et al.'s critique that "the number of cognate sets proposed for Transeurasian is exceptionally low" is based on comparison of our cognate set distributions with those of long-established language families, such as Sino-Tibetan and Indo-European. However, the authors ignored the full body of evidence in our traditional linguistic approach indicated in red in Table 11 and counted only the portion of semantically equivalent cognates used in our Bayesian analysis. In fact, the distribution is completely in line with expectations, considering that Transeurasian has an older age, less surviving daughter languages and early attestations of a relatively young age, so that traces of cognancy are more blurred than in the other families.

The age of the Transeurasian family is roughly 2 millennia older than that estimated for Sino-Tibetan and Indo-European in the sources mentioned. In the course of time, languages branching off from a common ancestor tend to become more and more distant as they undergo changes independently of each other. Thus, linguistic evidence of relatedness gets lost due to a cumulative effect of increasing divergence. This process of gradually decreasing evidence is called cognate attrition. It explains why traces of cognancy are more blurred for language families of relatively old age like Transeurasian.

Moreover, the considerable time depth of the Transeurasian family is combined with a relatively high extinction rate of the daughter languages. By way of comparison, the Sino-Tibetan and Indo-European language families respectively count about 500 and 445 living languages, whereas hardly 75 Transeurasian languages have survived to the present. This may be due to the pressure of some dominant languages, such as Middle Mongolian under Jinghis Khan or Old Korean of the Silla Kingdom, erasing all pre-existing Mongolic or Koreanic languages. So, in the Transeurasian case, many languages that may have reflected cognates will have died out. Hence, the effect of cognate attrition for the Transeurasian languages is expected to be much stronger than for Sino-Tibetan and Indo-European, because fewer languages are left to testify to relatedness.

Whereas the estimated time depth is about two millennia earlier, the written sources for the Transeurasian languages are roughly two millennia younger than those for Sino-Tibetan and Indo-European. The written records for Indo-European are dated at ca. 1000 B.C. for Sanskrit, ca. 800 B.C. for Homeric Greek, ca. 500 B.C. for Latin, ca. 400 A.D. for Gothic and ca. 800 A.D. for Old Church Slavonic, while Old Chinese goes back to ca. 1000 B.C. However, the first unambiguous and fully accessible attestations for Japanese are restricted to ca. 700 A.D., for Korean to ca. 1400 A.D., for Tungusicto ca. 1600 A.D., for Mongolic to ca. 1200 A.D. and for Turkic to ca. 700 A.D. Given this time gap of almost four millennia all together, we would expect more cognate attrition in the lexicon, less rigorous sound correspondences and less basic vocabulary, which is indeed the case for Transeurasian reconstruction.

**1.5 Sources**

Tian et al. complain that "the data sources for the 98 languages involved are not indicated anywhere, which makes it difficult to check that the forms cited actually exist and have the meaning listed." However, our Article (SI 5: 79-83) lists an extensive inventory of sources arranged by daughter branch. For the reader's convenience, we repeat these sources here, complemented with additional Tungusic, Koreanic and Japonic sources. In addition, the Method's section of our Article explains that we combined dictionary search with fieldwork. Our Article acknowledges co-authors Alexander Savelyev (Turkic), Joanna Dolińska (Mongolic), Ilya Gruntov (Mongolic), Olga Mazo (Mongolic), Sofia Oskolskaya (Tungusic), Seongha Rhee (Koreanic), Kyou-Dong Ahn (Koreanic), John Bentley (Japonic) as well as external collaborators Thomas Pellard (Japonic), Alexander Francis-Ratte (Japonic), Wayne Lawrence (Japonic), Aleksandra Jarosz (Japonic), Matthew Miller (Tungusic), Karina Mishchenkova (Tungusic), Elena Perekhvalskaya (Tungusic), Irina Nikolaeva (Tungusic), Patryk Czerwinski (Tungusic), Natalia Aralova (Tungusic), Ian Joo (Koreanic), Jan-Olof Svantesson (Mongolic), Maria Levy (Mongolic), Julie Lefort (Mongolic), Rottar Máté (Mongolic) for their collection of linguistic datasets.

#### 1.5.1 Japonic sources

Chamberlain, B. H. *Essay in aid of a grammar and dictionary of the Luchuan language* [Transaction of the Asiatic Society of Japan 23, suppl.]. ( Z. P. Maruya & Co., 1895)

Chiyo, K. & Toshizō, T. *Yoron-hōgen Jiten* [Yoron dialect dictionary] (Musashino Shoin, 2005).

Frellesvig, B. *A history of the Japanese language*. (Cambridge University Press, 2010)

Hirayama, T.. *Ryūkyū Miyako shotō hōgen kiso goi no sōgōteki kenkyū* [A composite lexicon of basic vocabulary of the islands of Miyako in the Ryukyus]. (Ōfūsha, 1983)

Hirayama, T., et al. *Gendai Nihon-go hōgen dai-jiten* Vol. 1-6 [Dictionary of the modern Japanese dialects]. (Meiji shoin, 1992)

Hirayama, T., Ōshima, I. and Nakamoto, M. *Ryūkyū-hōgen no sōgō-teki kenkyū* [A comprehensive study of the Ryūkyūan dialects]. (Meiji Shoin, 1966).

Jarosz, A. Nikolay Nevskiy's Miyakoan Dictionary: Reconstruction from the Manuscript and its Ethnolinguistic Analysis. Studies on the manuscript. (Poznań University PhD. Thesis, 2015).

Kokuritsu Kokugo kenkyūjo [National Institute for Japanese Language and Linguistics] *Okinawago jiten* [A dictionary of Okinawan]. ( Ōkurashō Insatsukyoku/Zaimushoo Insatsukyoku, 1963/2001).

Lange, R. A.. *The phonology of eighth-century Japanese; a reconstruction based upon written records*. (Sophia University, 1973).

Lawrence, W. & Takahiro, O. Tokunoshima Asama hougen no rigen meishi akusento shiryou [Accent materials of Asama dialect (Tokunoshima) dialect nouns]. *Ryuukyuu-no Hougen* **33**, 173-180 (2009).

Martin, S. E. *The Japanese language through time* (Yale Univ. Press, 1987).

Martin, S. E. Shodon: A Dialect of the Northern Ryukyus, *Journal of the American Oriental Society* 90, 1: 97–139 (1970).

Miyagi, S. *Ishigaki hōgen jiten* [A dictionary of Ishigaki dialect]. (Okinawa Taimususha, 2003).

Miyara, T. *Yaeyama goi*. Volume 8 of *Miyara Tōsō zenshū*. (Dai'ichi Shobō, 1981).

OCOJ The Oxford Corpus of Old Japanese (http://vsarpj.orinst.ox.ac.uk/corpus/index.html)

Ogawa, M. *Tarama-shima chōsa hōkokusho*. (Okinawa Kokusai Daigaku Nantō Bunka Kenkyūjo, 1993)

Omodaka, H. et al., eds.. *Jidaibetsu kokugo daijiten: Jōdai hen* [Great Dictionary of Japanese by period: Ancient period]. (Sanseidō, 1967)

Osada, S. *et al. Amami hōgen bunrui jiten*. 2 Volumes. (Kasama Shoin, 1980).

Sadayoshi, T. *Miyako Irabu-hōgen Jiten* [Irabu dialect dictionary] (Okinawa Times, 2013).

Seizen, N. *Okinawa Nakijin-hougen jiten* [Nakijin dialect dictionary] (Kadokawa, 1983).

Shin'ichi, K. *Yaeyama hatoma-hougen* (1960).

Shin'ichi, K. Hatoma-hougen-no on'in-taikei-ni tsuite [On the phonological system of Hatoma dialect]. *Ryuukyuu-no Hougen* **3**, 3-55 (1961).

Shin'ichi, K. Yaeyama Hatoma-hougen-no joshi [Yaeyama Hatoma dialect particles]. *Ryuukyuu-no Hougen* **8**, 122-142 (1984).

Shin'ichi, K. *Yaeyama-hougen-no hikaku-on'in-josetsu* [Introduction to the comparative phonology of the Yaeyama dialects] (Ryuukyuu-no Hougen Ronsou, 1987).
Shin'ichi, K. Hatoma-hougen: Shoku-kankei-go'i [Hatoma dialect: food-related vocabulary]. *Ryuukyuu-no Hougen* **14**, 32-89 (1989).
Shin'ichi, K. Hatoma-hougen-no juu-kankei-go'i [Hatoma dialect housing-related vocabulary]. *Ryuukyuu-no Hougen* **15**, 51-106 (1991).
Shin'ichi, K. Hatoma-hougen-no saishi-kankei-go'i [Hatoma dialect festival-related vocabulary]. *Ryuukyuu-no Hougen* **16**, 56-104 (1992).
Shin'ichi, K. Hatoma-hougen-no jintai-kankei-go'i [Hatoma dialect human body-related vocabulary]. *Ryuukyuu-no Hougen* **18/19**, 215-229 (1995).
Shin'ichi, K. in *Nihongo-bunmatsushi-no rekishiteki kenkyuu* [Historical research into Japanese sentence-final particles], 237-253 (Miyai Shoten, 1998).
Shin'yū, M. *Ishigaki-hōgen Jiten* [Ishigaki dialect dictionary] (Okinawa Times, 2003).
Shimoji, K. *Miyako guntōgo jiten* [A dictionary of the language of the Miyako Islands]. (Shimoji Yoneko, 1979).
Shimoji, M. *A Grammar of Irabu, a Southern Ryukyuan Language*. (Ph.D. dissertation, The Australian National University, 2008).
Suma, O. & Nahoko, S. (eds) *Amami-hōgen Bunrui-jiten (Gekan)* [Amami dialect classified dictionary Vol. 2] (Kasama Shoin, 1980).
Takahashi, T. Yaeyama-Yonaguni. Volume 12 of *Ryūkyū no hōgen*. (1987).
Takahashi, T. *Tarama hōgen no goi -vol. 2*. (Okinawa Kokusai Daigaku Nantō Bunka Kenkyūjo, 1994)
Takahiro, O. Tokunoshima no minwa Kucu to Kwata [A folktale of Tokunoshima "Kutsu and K'wata (stupid brothers)"], *Amami-Okinawa Minkan Bungeigaku* [The science of Amami-Okinawa folk literature] **8**, 110-116 (2008).
Yonaha, Y. *Mya-ku sumafutu jiten* [A dictionary of Miyako dialect]. (Okinawa Koroni, 2003).
Zendō, U. in *Komatsushiro Yūichi kyōju Taishoku / Shima Minoru kyōju Taikan Kinen Kokugogaku Ronshū* [Papers in Japanese Language Studies in Honour of the Retirements of Professors Komatsushiro Yūichi and Shima Minoru] (ed Iwate Kokugo Gakkai Editorial Board), 188-220 (1977)
Zendō, U. Tokunoshima Asama-hougen-no Fukugou-doushi-no akusento [Accentuation of compound verbs in the asama dialect of Tokunoshima]. *Ryuukyuu-no Hougen* **28**, 1-42 (2004).

#### 1.5.2 Koreanic sources

Ahn, O.-K. *Ewen sacen* [A dictionary of etymology] (Hankuk Publisher 1996 [1989]).
Bae, D.-O. On the method of Old Korean study. *Paytalmal* **39**, 325-345 (2006).
Bae, D.-O. Itwuwa kwanlyentoyn mech kaci sayngkak [Some thoughts on Idu]. *Proceedings of the Conference of the Society of Kugyol,* 83-84 (2011).
Baek, M.-S. *Wulimal ewen sacen* [A dictionary of Korean etymology] (Pagijong Press, 2015 [2014]).
Chang, K.-J. On grammatical forms used in point-attached kugyol materials. *Journal of Korean Linguistics* **56**, 249-279 (English abstract, 317) (2009).
Chang, K.-J. An introduction to Seokdokkugyol materials made during the era of Korea dynasty and their utilization. *Korean Linguistics* **59**, 1-38 (2013).
Cho, H.-B. (ed) *Kwuke ewenyenkwu chongsel* (1) [A collection of Korean etymological studies] (Thayhaksa, 2014).
Cho, Y.-E. *Hankwuke ewensacen* [A dictionary of Korean etymology] (Dasom Publishing, 2004).
Choe, H. C. & Kim, M. R. *Wulimal ewensacen* [A dictionary of Korean etymology] (1999).
Choi, C.-R. *Wulimal ewenyenkwu* [A study of Korean etymology] (Iljisa, 1986).
Choi, C.-R. *Ewensanchayk* [A stroll into etymology] (Hanshin Publishing, 1992).
Choi, N.-H. Kotay kwukeuy coepep yenkwu [A study of Old Korean word formations]. *Han-Geul* **220**, 109-152 (1993).
Choi, N.-H. *Kotaykwuke hyengthaylon* [Old Korean morphology] (Pagijong, 1996).
Chon, S. Y. *Kotaykwukeuy ehwiyenkwu* [A study on Old Korean vocabulary] (Research Institute of Korean Studies, Korea University, 1990).
Gang, G. H. *A study of Idu on Simyangjanggye* MA thesis (Chonnam National University, 2001).

Ha, Y. W. et al. (eds) *Yeysphyenci nathmalsacen* [A lexicon of old letters] (Dolbegae, 2013).
Hong, S. M. *Kwuke ehwiuymiuy sacekpyenchen* [Historical change of Korean word meanings] (Hankuk Publisher, 2003).
Hong, Y. P. in *Kwukeuy sitaypyel pyenchen yenkwu 2: Kuntaykwuke* [Studies of historical change of Korean by period 2: Early Modern Korean] (ed National Institute of the Korean Language) 191-234 (NIKL, 1997).
Hong, Y. P. *Salaissnun wulimaluy yeksa* [A history of the living Korean] (Thayhaksa, 2009).
Hong, Y. P. Kwuke ehwisa yenkwu pangpep [Research methods on the history of Korean words]. *Proceedings of the Conference of the Society of Kugyol*, 9-36 (2014).
Hong, Y. P., Song, K.-J., Chung, K. & Song, C. E. (eds) *17-seyki kwukesacen* [A dictionary of the 17th century Korean] (The Academy of Korean Studies, 1995).
Hongo, T. The honorific forms of the Idu in the Nongpojip. *Journal of Kugyol Studies* **7**, 79-111 (2001).
Hongo, T. *The honorific forms in Idu texts.* Ph.D. dissertation (Korea University, 2002).
Hwang, K.-J. A study on the original phonetic value of coda 's' of the 15th century: Centering around the reading of *eumjilda. Korean Language and Literature in International Context* **32**, 25-62 (2004).
Hwang, M.-H. Cosen sitay enkankwa kwuke saynghwal [Enkan letters and the life of Korean in the Joseon Dynasty times]. *Saykwukesaynghwal* **12.2**, 133-145 (2002).
Hwang, S.-Y., Lee, J. & Ha, G. *Sektokkwukyelsacen* [A dictionary of Sektokkwukyel] (Bakmunsa, 2009).
Jeong, H. *Cwungseykwuke uyconmyengsa yenkwu* [A study of dependent nouns in Middle Korean] (Hyungseul Publishing, 1987).
Jeong, H. *Wulimaluy sangsanglyek* [Imaginative powers in Korean, revised edition) (Jimoondang, 2014).
Jun, J.-H. *Kwuke ehwisa yenkwu* [A study of the Korean word history] (revised edition) (Kyungpook National University Press, 1992).
Kang, H.-K. *Hankwuke ewenyenkwusa* [The history of the Korean etymological studies] (Jipmoondang, 1989 [1988]).
Kang, H.-K. Myech taneuy ewen [On etymology of some words]. *The Institute of Education of Korean Language and Literature* **5,** 177-189 (1997).
Kang, K.-W. *Hyangkasinhaytokyenkwu* [A new interpretation of Hyangka] (Hankuk Publisher, 2004).
Kim, D.-S. *A study on the change of Old Korean words.* Ph.D. dissertation (Sungkyunkwan University, 1987).
Kim, D. Kotay hankwukeuy conghapcek yenkwu (1) [A comprehensive study of Old Korean (1)]. *Han-Geul* **227**, 5-37 (1995a.)
Kim, D. Kotay hankwukeuy conghapcek yenkwu (2) [A comprehensive study of Old Korean (2)] *Han-Geul* **227**, 38-70 (1995b).
Kim, I.-H. *Cosene ewenphyenlam* [A handbook of the Joseon Dynasty language, 2 volumes] (Pagijong, 2001).
Kim, J. H. Investigation of linguistic origin in relation to the word 'fire'. *Korean Linguistics* **39**, 41-78 (2008).
Kim, J.-T. Ilponsekiey nathanan kotayhankwuke calyo yenkwu [A study of Old Korean material observed in Ilponseki (Nohon Shoki)]. *The Journal of Korean Language and Literature Education* **30**, 167-214 (1998).
Kim, J.-T. Kotaykwuke ehwihyengthayuy naycek caykwu [On internal reconstruction of Old Korean words]. *The Journal of Korean Language and Literature Education* **32**, 145-160 (2000).
Kim, K. J. The connection of a hyangga and that Chinese poetry. *Yelsangkocenyenkwu* **56**, 105-138 (2017).
Kim, M.-S. *Wulimal ewen sacen* [A dictionary of Korean etymology] (Thayhaksa, 1997).
Kim, M. R. *Hankwuke ewen sacen* [A dictionary of Korean etymology] (K&C Books, 2015).
Kim, T.-K. *Hampuk Pangen Sacen* [Dictionary of North Hamkyeng dialects]. (Kyonggi University Press, (1986)
Kim, W.-J. Hyangka haytokuy silcey [Practices in Hyangka interpretation]. *Journal of Korean Linguistics* **8**, 1-21 (1979).
Kim, Y. W. Wooden documents and words. *Journal of Kugyol Studies* **33**, 5-22 (2014).
Kim, Y.-B. Translation of Buddhist sutra in the Early Choseon Dynasty. *Maha Bodhi Thought* **5**, 9-40 (2002).
Kim, Y. The decipherment of Gyunyeo's hyangga Gwangsugongyangga. *Journal of Kugyol Studies* **25**, 47-81 (2010).
KLS [Korean Language Society] *Uli-mal khun sacen* [A great dictionary of the Korean language]. (Emunkak,

(1991).
Ko, J. E. (aka Go, J.-W.) Taymyenglyulcikhayuy itwuwa ku thukcing [The Idu texts in Taymyenglyulcikhay and their characteristic features]. *Journal of Kugyol Studies* **9**, 19-72 (2002).
Ko, J. E. (aka Go, J.-W.) A review of the study on Kugyol. Journal of Kugyol Studies **12**, 5-46 (2004).
Ko, Y.-K. Cheyongkauy han haytok [An interpretation of Cheyongka]. *Konkukemwunhak* **9/10**, 903-911 (1985).
Ko, Y.-K. Yangcwutonguy kwukehak yenkwu [A review of Yang Joo-Dong's studies on Korean]. *The Korean Language and Literature* **133**, 5-49 (2003).
Lee, B.-G. Kukyel calyouy ehwi [Vocabulary in Kugyol texts]. *Journal of Kugyol Studies* **33**, 23-61 (2014).
Lee, D. J. The meaning of inscription on ink-inscribed pottery excavated from Weolji Pond in the Silla Dynasty. *The Chin-Tan Hakpo* **131**, 1-24 (2018).
Lee, E.-K. 'Kwukyelcalyouy ehwi'-ey tayhan tholonmwun [A commentary on 'On the Kugyol vocabulary']. *Proceedings of the Conference of the Society of Kugyol,* 81-82 (2014).
Lee, I.-M. *Kocenkwukeuy yenkwu* (Sunmyung Publishing, 1971).
Lee, J.-H. Basic work for the word research of ancient Korean language: Understanding of texts and data. *Journal of Kugyol Studies* **33**, 63-96 (2014).
Lee, J. Mwunul nathanaynun kotaykwuke ehwiey tayhaye [On the words signifying 門 'door']. *Salimemwunyenkwu* **20**, 175-190 (2010).
Lee, K.-G., et al. *Cennam Pangen Sacen* [Dictionary of Southwestern dialects]. (Top Chwulphansa, (1998).
Lee, K.-M. *Kwuke umwunsa yenkwu* [A study of the phonological history of Korean] (Tower Press, 1972).
Lee, K.-M. Itwuey tayhaye [On Idu]. *The Korean Language and Literature* **86**, 286-287 (1981).
Lee, K.-M. & Ramsey, R. S. *A History of the Korean Language* (Cambridge University Press, 2011).
Lee, N.-D. *Hankwuke ewen yenkwu I: Wensihankwukeuy thamkwu* [Studies on Korean etymology I: An exploration into Proto-Korean] (Ewha Woman's University Press, 1993 [1985]).
Lee, N.-D. *Hankwuke ewen yenkwu II: Tongsa ehwiuy ewen* [Studies on Korean etymology II: Etymology of verbs] (Ewha Woman's University Press, 1995 [1985]).
Lee, N.-D. *Hankwuke ewen yenkwu III: Hyengyongsa ehwiuy ewen* [Studies on Korean etymology III: Etymology of Adjectives] (Ewha Woman's University Press 1998 [1985]).
Lee, N.-D. *Hankwuke ewen yenkwu IV: Eneyentayhakcek kochalkwa umwuntayungpepchikuy cenglip* [Studies on Korean etymology IV: A glottochronological exploration and establishment of the principles of phonemic contrast] (Ewha Woman's University Press, 1998 [1986]).
Lee, S. J. *A study on Yi-du of the Koryo period* (Thayhaksa, 1992).
Lee, S.-U. *Kwuke hyengthaysa yenkwu* [A study of historical morphology in Korean] (Thayhaksa, 1997).
Lee, Y. Cye-uy yeksacek kochal [A historical investigation on *-cye*]. *The Chin-Tan Hakpo* **105**, 187-206 (2008).
Lee, Y. Sinlaitwuey nathanan *nay*-ey tayhaye [On nay 內 in the Idu texts of the Silla Dynasty times]. *Proceedings of the Conference of the Society of Kugyol*, 53-71 (2011).
Martin, S. E. *A Reference Grammar of Korean* (Charles E. Tuttle, 1992).
Nam, K.-W. Kotaykwuke coepepuy han kochal: Swukay ehwiuy phasayngelul cwungsimulo [An investigation into word formation rules in Old Korean: Focusing on selected words]. *Han-Geul* **121**, 339-365 (1957).
Nam, P.-H. Wenwangsayngkauy saylowun haytok [A new interpretation of Wenwangsayingka]. *Journal of Kugyol Studies* **41**, 5-27 (2018).
Nam, P.-H. Kwuke sokuy chayonge: Kotaykwukeeyse kuntaykwukekkaci [Loan words in Korean: From Old Korean to Early Modern Korean]. *Kwukesaynghwal* **2**, 6-22 (1985).
Nam, P.-H. *Itwuyenkwu* [Idu Studies] (Thayhaksa, 2000).
Nam, P.-H. *Kwukesalul wihan kwukyelyenkwu* [Kugyol studies for Korean historical linguistics] (Thayhaksa, 2002 [1999]).
Nam, P.-H. On the sentence final partikle (sic) in Old Korean Yidu. *Journal of Kugyol Studies* **15**, 5-28 (2005).
Nam, P.-H. A new readings (sic) of Heonwhaga. Journal of Kugyol Studies **24**, 5-35 (2010).
Nam, P.-H. *Kotayhankwuke nonko* [Studies on Old Korean] (Thayhaksa, 2014).
National Institute of the Korean Language. *Kwukeuy sitaypyel pyenchen-silthayyenkwu 1: Cwungseykwuke* [Studies on the change and state of affairs of Korean by the period 1: Middle Korean] (NIKL, 1996).
National Institute of the Korean Language. *Kwukeuy sitaypyel pyenchen yenkwu 2: Kuntaykwuke* [Studies on the

change of Korean by the period 2: Early Modern Korean] (NIKL, 1997).
National Institute of the Korean Language. *Kwukeuy sitaypyel pyenchen yenkwu 3: Kotaykwuke* [Studies on the change of Korean by the period 3: Old Korean] (NIKL, 1998).
National Institute of the Korean Language. *Kwukeuy sitaypyel pyenchen yenkwu 4: Kayhwaki kwuke* [Studies on the change of Korean by the period 4: Korean in the Modern Reformation period] (NIKL, 1999).
National Institute of the Korean Language. *Tanepyel ewen cengpo* [Etymological information of words] (Human Culture Arirang, 2015).
Park, H. S. Taymyenglyulcikhay itwumwuney kwanhan yenkwu [A study of the Idu in Taymyenglyulcikhay]. *The Journal of Kwandong University* **10.1**, 1-20 (1982).
Park, H. S. *A study of 'I-doo' (Old Korean phonetic symbol) on Dai-myung-ryul-jik-hae*. Ph.D. dissertation (Myungji University, 1984).
Park, J.-Y. *Koetaysacen* [A comprehensive dictionary of Old Korean, 8 volumes) (Hakkobang, 2010).
Park, J. in *Kwukeuy sitaypyel pyenchen yenkwu 3: Kotaykwuke* [Studies on the change of Korean by the period 3: Old Korean] (ed NIKL), 121-205 (NIKL, 1998).
Park, S. *A linguistic study on Idu of the early choson period.* Ph.D. dissertation (Seoul National University, 1996).
Park, S. Chacaphyokiuy ehwilon [On borrowed character notation]. *Saykwukesaynghwal* **7.4**, 61-77 (1997).
Park, S. in *Kwukeuy sitaypyel pyenchen yenkwu 3: Kotaykwuke* [Studies on the change of Korean by the period 3: Old Korean] (ed NIKL), 77-120 (NIKL, 1998).
Park, S. A comprehensive study on Idu in the historical manuscripts of the 16th century. *Sengkoknonchong* **36.1**, 49-107 (2005).
Park, S. Retrospect and prospect of scholarly study on Idu in Korean old documents. *Journal of Kugyol Studies* **21**, 169-201 (2008).
Park, S. Cosen sitayuy itwuwa ku yenkwu pangpepuy phyenmo [On the Idu texts in the Joseon Dynasty times and some methodological aspects of their study]. *Proceedings of the Conference of the Society of Kugyol,* 33-53 (2011).
Park, S. Kotay hankwuke phyokieyseuy pyencheyhanmwunkwa itwu [On modified Chinese characters and Idu in the notation of Old Korean]. *Hankwukesa Yenkwu* **4**, 49-80 (2018).
Park, Y.-S. *Chokanpon twusienhay ehwicalyocip* [A lexicon of the first edition of Twusienhay] (Pagijong, 1998).
Ramstedt, G. J. *Studies in Korean etymology* [Suomalais-Ugrilaisen Seuran Toimituksia 95] (Suomalais-Ugrilainen Seura, 1949).
Ramstedt, G. J. *Additional Korean etymologies* [Suomalais-Ugrilainen Seuran Aikakauskirja] (Helsinki:Suomalais-Ugrilainen Seura, 1954).
Ramstedt, G. J. *A Korean grammar* [Suomalais-Ugrilaisen Seuran Toimituksia 82] (Anthropological Publications, 1968 [1939]).
Ramstedt, G. J. *Paralipomena of Korean etymologies* [Suomalais-Ugrilaisen Seuran Toimituksia 182]. Suomalais-Ugrilainen Seura, 1982).
Ramstedt, G. J. *A Korean grammar* (Suomalais-Ugrilainen Seura, 1997[1939]).
Rhee, S. *Semantics of verbs and grammaticalization: The development in Korean from a cross-linguistic perspective.* Ph.D. dissertation (Hankuk Publisher, 1996).
Ryu, C.-D. *Koesacen* [A dictionary of Old Korean] (Dongguk Publishing, 1955).
Ryu, C.-D. Hesahwa kokwu [From content word to function word]. *Inmwunkwahak* **7**, 1-23 (English abstract, 445) (1962).
Ryu, C.-D. *Ehwisayenkwu* [A study of word history] (1978).
Ryu, C.-D. *Icokwukesa yenkwu* [A historical study of Korean in the Joseon Dynasty times] (Iwoo Publishing (Revised ed. by Samwoo Publishing), 1979[1964]).
Ryu, C.-D. *Kwukepyenchensa* [A history of Korean language change] (Tongmoonkwan, 1980[1961]).
Ryu, C.-D. *Icoe sacen* [A dictionary of Korean in the Joseon Dynasty times] (Yonsei University Press, 2000[1964]).
Ryu, R. *Hyangkayenkwu* [A study of Hyangka] (Pagijong Press, 2003).
Shim, J.-K. (ed) *Kwuke ehwiuy kipankwa yeksa* [On the foundation and history of Korean vocabulary]

(Thayhaksa, 1998).
Shin, J. H. *Hyangkauy haysek* [An interpretation of Hyangka] (Jipmoondang, 2002).
Song, K.-J. Kotaykwuke yenkwuuy saylowun pangpep [A new method of Old Korean studies]. *Hanguk Munhwa: Korean Culture* **24**, 1-34 (1999).
Song, K.-J. Korean Transcriptions by the readings in meaning values of Chinese characters found in geographical names of the 19th century. *Journal of the Place Name Society of Korea* **6**, 177-216 (2001).
Song, K.-J. Hwunmincengum haylyeyuy umso-umsenghak [On phonemes and phonetics in Hunimjeongum Illustrations]. *Hankwukesa Yenkwu* **1**, 59-94 (2014).
Song, S.-J. *Ceycwumal Kun Sacen* [Great Cheju Dictionary]. (Hankuk Munhwasa, 2007)

Suh, J.-B. *Wulimaluy ppwuli* [Roots of Korean] (Korea One Books, 1996 [1989]).

Suh, J.-B. *Ewenuy seykyey* [The world of etymology] (Ubiquitous Culture Leader, 2005).
Whitman, J. et al. Toward an international vocabulary for research on vernacular readings of Chinese texts (Hanwen Xundu). *Scripta* **2**, 61-63 (2010).
Yang, H.-C. Kyunye [Wenwangka] yenkwuuy hyenwichi: Hyangchalcek sakowa sicek haytokul kyemhaye [On the status quo of Kyunye's Wenwangka studies: Focusing on Hyangchal-based reasoning and poetic interpretations]. *Mosanhakpo* **9**, 257-290 (1997).
Yi, D. L. Kotaykwuke ehwicalyo yenkwu (1). *Sungkyunemwunyenkwu* **32.1**, 5-18 (1997).
Yi, D. L. A study of lexical interpretation of Koryo Gayo. *Inmwunkwahak* **32**, 23-39 (2002).
Yoo, C.-K. Hyangkahaytokuy pansengcek kochal [Reflections on Hyangka interpretations]. *Mosanhakpo* **9**, 1-54 (1997).

### Tungusic sources

An, J. Hezheyu jianzhi [Introduction to the Hezhen Language]. (Minzu chubanshe, 1986)
Avrorin, V. A. Grammatika nanajskogo jazyka [The Grammar of Nanai]. Vol. 1. (Publishing house of Academy of Sciences of USSR, 1959)
Avrorin, V. A. Grammatika nanajskogo jazyka [The Grammar of Nanai]. Vol. 2. (Publishing house of Academy of Sciences of USSR, 1961)
Avrorin, V. A. Grammatika man’čžurskogo pis’mennogo jazyka [The grammar of the written Manchu language]. (Nauka, 2000)
Avrorin, V. A. and Lebedeva, E.P. 1978. Oročskie teksty i slovar’ [Oroch texts and dictionary]. (Nauka, 1978).
Benzing, J. Die tungusischen Sprachen: Versuch einer vergleichenden Grammatik. *Abhandlungen der geistes- und sozialwissenschaftlichen Klasse 11*. 949-1099. (Verlag der Akademie der Wissenschaften und der Literatur in Mainz, 1955)
Chaoke, D. O.. Xiandai Xiboyu Kouyu Yanjiu [A study of the modern spoken Sibe language]. (Minzu Chubanshe, 2006).
Cincius, V. I. Sravnitel’naja fonetika tunguso-man’čžurskich jazykov [Comparative phonetics of Manchu-Tungus languages]. (Učpedgiz, 1949)
Cincius, V. I. (ed.). Sravnitel’nyj slovar’ tunguso-man’čžurskix jazykov [Comparative Dictionary of the Tungus-Manchu Languages]. Vols. 1, 2. ( Nauka 1975, 1977).
Cincius, V. I. Negidal’skij jazyk [The Negidal language]. (Nauka 1982).
Dorji, D. Ehan cidian [An Evenki-Chinese dictionary]. (Neimenggu wenhua chubanshe, 1998)
Girfanova, A. Kh. Slovar’ udegejskogo jazyka [Udihe Dictionary]. (Nauka, 2001)
Gorelova, L. M. Manchu grammar. (Brill, 2002)
Hu, Z. Elunchun-yu jianzhi [Concise grammar of Oroqen]. (National Minorities Publ., 1986)
Hu, Z. and Chaoke. Ewenkeyu jianzhi (A sketch of the Ewenke language). (Minzu chubanshe, 1986).
Ikegami, J. Uirutago jiten [A Dictionary of the Uilta Language]. (Hokkaido University Press, 1997)
Janhunen, J. Material on Manchurian Khamnigan Evenki. (Castrenianum, 1991)
Kane, D. The Sino-Jurchen Vocabulary of the Bureau of Interpreters. (Indiana University, Research Institute for Inner Asian Studies, 1989)
Kazama, S. Basic Vocabulary (A) of Tungusic Languages. Endangered Languages of the Pacific Rim A2-037. (Kyoto, 2003)
Kormušin, I. V. Udyhejskij (udegejskij) jazyk [The Udihe language]. (Nauka, 1998)

Korovina, E. Leksika prirodnogo i kul'turnogo okruženiya v Tunguso-Man'čžurskich jazykach (v istoriko-tipologičeskom osveščenii) [Tungus-Manchu vocabulary related to natural environment and cultural activities (in historical and typological perspective)]. ( Russian State University of Humanities, BA dissertation, 2011)
Kubo, T., Kogura, N. & Sheng Zhuang. Shibe go goi shu [Sibe vocabulary]. ( Research Institute for Languages and Cultures of Asia and Africa Tokyo University of Foreign Studies, 2011)
Lebedeva, E. P. Oročskij jazyk [the Oroch language]. Jazyki mira: Mongolskie jazyki. Tunguso-manchzhurksie jazyki. Japonskij jazyk. Korejskij jazyk, ed. by Vladimir M. Alpatov, Igor V. Kormushin, Grigorij C. Pjurbeev and Olga I. Romanova, 215–226. (Indrik, 1997)
Li, F. & Whaley, L. J. Oroqen vocabulary. World loanword database, ed. by Martin Haspelmath, Uri Tadmor. (Max Planck Institute for Evolutionary Anthropology, 2009)
Li, L. Hojengo no dōshi kōzō [Verbal structure of Hezhe]. Chiba University Doctoral dissertation, 2006)
Li, S. Xibo-yu jianzhi [Concise grammar of Sibo]. (Zhongguo shaoshu minzu yuyan jianzhi congshu.) (National Minorities Publ., 1986)
Menges, K. H. Die tungusischen Sprachen. Tungusologie. (Brill, 1968)
Mishchenkova, K. O.. Refleksy praevenkijskogo *s v govorax evenkijskogo jazyka v kontse XVII v. i pervoj polovine XVIII v. [Reflections of the Proto-Evenki *s in the Evenki dialects in the late 17th century and the first half of the 18th century]. *Uralo-altajskie issledovanija* 3(34), 72-83. (2019)
Nikolaeva, I., Tolskaya, M..*A Grammar of Udihe*. (Mouton de Gruyter, 2001)
Nikolaeva, I. *A Historical Dictionary of Yukaghir*. (Mouton de Gruyter, 2006)
Norman, J.A Concise Manchu-English Lexicon. (University of Washington Press, 1978)
Novikova, K. A. Očerki Dialektov Evenskogo Jazyka: Ol'skij Govor 1 [Sketches of the Even dialects: Olsky dialect 1]. (Academy of Sciences of the USSR, 1960)
Novikova, K. A. Očerki Dialektov Evenskogo Jazyka: Ol'skij Govor [Sketches of the Even dialects: Olsky dialect]. (Academy of Sciences of the USSR, 1980)
Onenko, S. N. Nanajsko-russkij slpvar' [Nanai-Russian dictionary]. (Russkij jazyk, 1980)
Ozolinja, L. V. & Fedjaeva, I. J. *Oroksko-russkij i russko-orokskij slovar'* [Orok-Russian and Russian-Orok dictionary]. (Sakhalinskoe knizhnoe izdatel'stvo, 2003)
Petrova, T. I. Jazyk orokov (ulta) [Language of the Orok (Uilta)]. (Nauka, 1967)
Schmidt, Peter. The language of the Oroches. *Acta Universitatis Latviensis* 17, 2–62 (1928)
Sem, L. I. Ocherki dialektov nanajskogo jazyka. Bikinskij (ussurijskij) dialect [Outlines of the Nanai dialects. The Bikin (Ussuri) dialect]. (Nauka, 1976).
Sunik, O. P. Kur-urmijskij dialekt. Materialy i issledovanija po nanajskomu jazyku [Kur-Urmi dialect. Materials and studies on the Nanai language]. (Uchpedgiz, 1958)
Sunik, O. P. Tunguso-man'čžurskije jazyki [Manchu-Tungus languages]. (Nauka, 1959)
Sunik, O. P.. Ulčskij jazyk. Issledovanija i materialy [The Ulch language. Studies and materials]. (Nauka, 1985)
Tsumagari, T.. A basic vocabulary of Khamnigan and Oluguya Ewenki in Northern Inner Mongolia. *Bulletin of the Institute for the Study of North Eurasian Cultures* 21, 83-103. (1992)
Vasilevič, G. M.. *Ocherki dialektov evenkojskogo /tungusskogo/ jazyka* [Outlines of the Evenki /Tungus/ dialects]. (Uchpedgiz, 1948)
Vasilevič, G. M. Evenkijsko-Russkij Slovar' [Evenki-Russian dictionary]. (Gosudarstvennoe Izdatel'stvo Inostrannyx i Natsional'nyx Slovarej, 1958)
Vasilevič, G. M. K. Voprosu o klassifikacii tunguso-man'čžurskih jazykov [About the question of classification of Manchu-Tungus languages]. (Nauka, 1960)
Vasilevič, G. M. Značenie dnevnikov Messerschmidta dl'a tungusovedenija [Value of Messerschmidt's notes for Tungusic studies] in *Izvestija Sibirskogo otdelenija Akademii Nauk SSSR*, 6, 2,116-122 (1969).
Xuejuan, H. Binwei de hezhe yu [Endangered Hezhe]. (Heilongjiang Education Press, 2005)
Zhang, P. *The Kilen language of Manchuria*. (PhD Thesis. Univeristy of Hong Kong, 2013)
Zhang, Y., Zhang, Xi & Dai, S.. The Hezhen Language. (Jilin University Press, 1989)
Yang, Z. & Fuluntai, Y.. Sibe Nikan gisun kamcibuha tacibure buleku bithe [Xibe-Chinese teaching dictionary]. (Xinjiang renmin chubanshe, 1998)

### Mongolic sources

Bawden, Ch. *Mongolian-English dictionary* (Kegan Paul International, 1997).
Bolčuluu, et al. *Yuɣur kelen-ü üges / Dōngbù Yùgùyǔ cíhuì* [Vocabulary of Eastern Yugur] (1984, 1985).
BAMRS. Bolshoy akademicheskiy mongolsko-russkiy slovar [Big academic Mongolian-Russian dictionary] (2001)
Čenggeltei, et al. Mongɣor kele ba Mongɣol kele / Tǔzúyǔ hé Měnggǔyǔ [Monguor and Mongolian] (Nèi Měnggǔ rénmín chūbǎnshè, 1988 [1991]).

Chen, Y. *Dongxiang zu zizhixian Naleisi xiaoxue shuangyu jiaoxue shiyan ban shiyong jiaocai* [Textbook for bilingual teaching experimental class of Narisi Primary School in the Dongxiang Autonomous County] (Manuscript, 2002).

Chen Z. et al. *Folktales of China's Minhe Mangghuer* (LINCOM Publishers, 2005).

Cheremisov, K. M. *Buryat-mongolsko-russkiy slovar'* [Buryat-Mongol-Russian dictionary] (Gos. izd-vo-innostrannyx i nacionalnyx slovarej, 1951).

Damdinov D.G. & Sundueva E.V. Khamnigansko-Russkiy slovar [Khamnigan-Russian dictionary] (Irkutsk, 2015).

Dpal-ldan-bkra-shis, et al. Language Materials of China's Monguor Minority: Huzhu Mongghul and Minhe Mangghuer. *Sino-Platonic Papers* **69**, (University of Pennsylvania, 1996).

Enkhbat et al. *Daɣur kelen-ü üges / Dáwò'ěryǔ cíhuì* [Vocabulary of Dagur] (Neimenggu renmin chubanshe, 1984).

Godziński, S. *Język średniomongolski: słowotwórstwo, odmiana wyrazów, składnia* (Wydawnictwa Uniwersytetu Warszawskiego, 1985).

Haenisch, E. *Wörterbuch zu Manghol un niuca tobca'an (Yüan- chcao pi-shi), Geheime Geschichte der Mongolen* (Otto Harrassowitz, 1939).

Iwamura, S. *The Zirni Manuscript. A Persian-Mongolian Glossary and Grammar* (Kyoto University, 1961).

Junast [Zhaonasitu] *Dongbu Yuguyu Jianzhi* [Introduction to the Eastern Yugur language] (Minzu chubanshe, 1981).

Junast [Zhaonasitu] Tǔzúyǔ jiǎnzhì [Concise grammar of Monguor] (Minzu chubanshe, 1981).

Junast [Zhaonasitu], & Lǐ, K. *Tǔzúyǔ Mínhé fāngyán gàishù* [General overview of the Mínhé dialect of Monguor]. *Mínzú Yǔwén Yánjiū Wénjí*, 458-487 (1982).

Kałużyński, S. Dagurisches Wörterverzeichnis nach F. V. Muromskis handschriftlichen Sprachaufzeichnungen. *Rocznik Orientalistyczny* **33/1**, 103-44, **33/2,** 109-43 (1970).

Kane, D. *The Kitan Language and Script* (Brill, 2009).

Khasbaatar, et al. *Mongɣor kelen-ü üges / Tǔzúyǔ cíhuì* [Vocabulary of Monguor] (1985 [1986]).

*Langjun 郎君. Da Jin huang di dutong jinglüe langjun xingji.大金皇弟都統經略郎君行記.* [Record of the Journey of the Younger Brother of the Emperor of the Great Jin Dynasty] (Engraved on a stele, 1134).

Lefort, J. *Contacts de langues dans le Nord-Ouest de la Chine: Le cas du Dongxiang* [Contact language in North-West Gansu: The case of Dongxiang] (École des Hautes Études en Sciences Sociales, Doctoral dissertation, 2012).

Lessing, F. *Mongolian-English Dictionary* (University of California Press, 1960).

Lǐ, K. *Monghul Qidar Merlong / Tǔ Hàn Cídiǎn* [Monguor-Chinese dictionary] (Qinghai Renmin, 1988).

Tuotuo *Liaoshi* [Official history of the Liao dynasty] (Zhonghua Shuju, 1974).

Ligeti, L. Un vocabulaire d'Istanboul. AOH **XIV**, 3-99 (1962).

Ligeti, L. O mongol'skix i tyurkskix yazykax i dialektax Afganistana. Acta Orientalia Hungarica **4**, 93–117 (1955).

Ligeti, L. Le lexique moghol de R. Leech. Acta Orientalia Hungarica **4**, 119–58 (1955).

Ma Peiting, Z. *保安语汉语词典 bǎo'ān yǔ hànyǔ cídiǎn* [A Bonan-Chinese Diectionary] (民族出版社 Mínzú chūbǎn shè, 2016).

Ma, G. & Chen, Y. *Dōngxiāngyǔ hànyǔ cídiǎn* (**东乡语汉语词典**) [Dongxiang-Chinese language dictionary] (Gansu minzu chubanshe, 2001).

*Měnghàn cídiǎn 蒙汉词典*[Mongolian-Chinese dictionary] (Nèiměnggǔ dàxué chūbǎnshè, 1999).

Muniev, B. D. *Kalmytsko-russkij slovar'* [Kalmyk-Russian dictionary] (Russkij jazyk, 1977).

Naixiong, Ch. (Ed.). *保安语词汇 Bǎo'ān yǔ cíhuì [*Vocabulary of the Bonan Language]. *蒙古语族语言方言研究丛书 Ménggǔ yǔzú yǔyán fāngyán yánjiū cóngshū* [Mongolian Language Dialect Research Series] (Nèiménggǔ rénmín chūbǎn shè, 1986).

Nugteren, H. *Mongolic phonology and the Qinghai-Gansu languages* (LOT. Leiden University Ph.D. dissertation, 2011).

Potanin, G. N. *Tangutsko-tibetskaja okraina Kitaja i central'naja Mongolija* (Imperatorskoe Russkoje Geograficheskoe Obshchestvo, 1893).

Ramstedt, G. J. Mogholica. Beiträge zur Kenntnis der Moghol-Sprache in Afghanistan. *Journal de la Société finno-ougrienne* **23:4, I-IV**, 1-60 (1906).

Ramstedt, G. J. *Kalmückisches Wöerterbuch* (Suomalais-Ugrilainen Seura, 1935).

Róna-Tas, Á. Khitan studies I. The graphs of the Khitan Small Script 1. General Remarks, Dotted Graphs, Numerals. *Acta Orientalia Academiae Scientiarum Hungaricae* **69(2)**, 117-138 (2016).

Shagdarov, L. D. and Kazanceva, A. M. O yazyke mogolov Afganistana (po materialam Sh. Iwamura i X. F. Shurmana. *O zarubezhnyx mongolovednyx issledovaniyax po yazyku* (Buryatskoe knizhnoe izdatel’stvo, 1968).
Saitō, Y. *The Mongolian Words in the Muqaddimat al-Adab: Romanized Text and Word Index* (The Japan Society for the Promotion of Science, 2008).
Sečenčogt Kāngjiāyŭ yánjiū [Kangjia language research] (Yuandong chubanshe, 1999).
Sevortyan E.V. *Etimologicheskiy slovar tyurkskih yazykov* [Etymological Dictionary of Turkic languages] (Moscow, 1978).
Slater, K. W. *A Grammar of Mangghuer: A Mongolic Language of China’s Qinghai-Gansu Sprachbund* (Routledge, 2003).
Smedt, A. de, & Mostaert, A. Le dialecte Monguor parlé par les Mongols du Kansou occidental. IIIe partie: Dictionnaire monguor-français (Imprimerie de l'Université Catholique, 1933).
Starostin, G. *Tower of Babel: an etymological database project.* http://starling. rinet.ru (2008).
Starostin, S. A., Dybo, A. & Mudrak, O. *Etymological Dictionary of the Altaic Languages* (Brill, 2003).
Todaeva, B. Kh. *Mongorskij jazyk. Issledovanie, teksty, slovar’* (Nauka, 1973).
Todayeva, B. Kh. *Dagurskij jazyk* [Dagur] (Nauka, 1986).
Tumurdei, G. & Cybenov, B. D. *Kratkij dagursko-russkij slovar’* [Brief Dagur-Russian dictionary] (BNC SO RAN, 2014).
Weiers, M. Das Moghol-Vokabular von W. R. H. Merk (Otto Harrassowitz, 1971).
Weiers, M. Die Sprache der Moghol der Provinz Herat in Afghanistan. Sprachmaterial, Grammatik, Wortliste. *Abhandlungen der Rheinisch- Westfahlischen Akademie der Wissenschaften **49*** (1972).
Weiers, M. *Schriftliche Quellen in Mogoli* (Westdeutscher Verlag, 1975).
Yingzhe, W. & Janhunen, J. *New Materials on the Khitan Small script* (Global Oriental, 2010).
*Xuwang muzhi* [Epitath of Prince Xu] (Epitaph, 1109).
Zhòng, S. *Dáwò’ěryŭ jiǎnzhì* [Concise grammar of Dagur] (1982).

### Turkic sources

Ašmarin, N. *Thesaurus Linguae Tschuvaschorum. Vol. 1–17* (Krasnyj Pečatnik & Čuvašskaja Kniga & Čuvašpoligrafizdat, 1928-1950).
Bailey, H. W. *Dictionary of Khotan Saka* (Cambridge Univ. Press, 1979).
Bammatov, Z. Z. *Russko-kumykskij slovar'* [Russian-Kumyk dictionary] (Gosudarstvennoe izdatel'stvo inostrannyx i nacional'nyx slovarej, 1960).
Bammatov, Z. Z. *Kumyksko-russkij slovar'* [Kumyk-Russian dictionary] (Sovetskaja enciklopedija, 1969).
Baskakov, N. A. & Toščakova, T. M. *Ojrotsko-russkij slovar'* [Oyrot-Russian dictionary] (Gosudarstvennoe izdatel'stvo inostrannyx i nacional'nyx slovarej, 1947).
Baskakov, N. A. *Karakalpaksko-russkij slovar'* [Karakalpak-Russian dictionary] (Gosudarstvennoe izdatel'stvo inostrannyx i nacional'nyx slovarej, 1958).
Baskakov, N. A. *Nogajsko-russkij slovar'* [Noghai-Russian dictionary] (Gosudarstvennoe izdatel'stvo inostrannyx i nacional'nyx slovarej, 1963).
Baskakov, N. A. *Turkmensko-russkij slovar'* [Turkmen-Russian dictionary] (Sovetskaja enciklopedija, 1968).
Baskakov, N. A. *Gagauzsko-russko-moldavskij slovar'* [Gagauz-Russian-Moldovan dictionary] (Sovetskaja enciklopedija, 1973).
Baskakov, N. A., Zajączkowski, A. & Szapszał, S. *Karaimsko-russko-pol'skij slovar'* [Karaim-Russian-Polish dictionary] (Russkij jazyk, 1974).
Baskakov, N. A. *Turecko-russkij slovar'* [Turkish-Russian dictionary] (Russkij jazyk, 1977).
Bektaev, Q. *Bol'šoj kazaxsko-russkij i russko-kazaxskij slovar'* [Big Kazakh-Russian and Russian-Kazakh dictionary] (Kazaxstanskij projekt razvitija gosudarstvennogo jazyka, 1995).
Bekturov, Š. & Bekturova, A. *Kazaxsko-russkij slovar'* [Kazakh-Russian dictionary] (Foliant, 2001).
Čeremisov, K. M. & Tsydendambaev, Ts. *Burjat-mongol'sko-russkij slovar'* [Buryat Mongol-Russian dictionary] (Gosudarstvennoe izdatel'stvo inostrannyx i nacional'nyx slovarej, 1951).
Chen Z. et al. *Folktales of China’s Minhe Mangghuer* (2005).
Clauson, G. *An Etymological Dictionary of Pre-Thirteenth-Century Turkish* (At the Clarendon Press, 1972).
Doerfer, G. & Tezcan, S. *Wörterbuch des Chaladsch (dialekt von Charrab)* (Akadémiai Kiadó, 1980).
Dybo, A. V. *Lingvističeskije kontakty rannix tjurkov. Leksičeskij fond. Pratjurkskij period* [Linguistic contacts of the early Turks. Vocabulary. Proto-Turkic period] (Vostočnaja literature, 2007).
Dybo, A. V. Vokalizm rannetjurkskix zaimstvovanij v vengerskom [Vocalism of early Turkic borrowings in Hungarian]. *Gedenkschrift von E.A. Helimsky* [*Finnisch-Ugrische Mitteilungen* **32–33**, 83–132 (2010).
Judakhin, K. K. *Kirgizsko-russkij slovar'* [Kirghiz-Russian dictionary] (Sovetskaja enciklopedija, 1985).

Kary-Nijazov, T. N. & Borovkov, A. K. *Uzbeksko-russkij slovar'* [Uzbek-Russian dictionary] (Izdatel'stvo Uzbekistanskogo filiala akademii nauk SSSR, 1941).
Kurpeško-Tannagaševa, N. N. & Apon'kin, F. Ja. *Šorsko-russkij i russko-šorskij slovar'* [Shor-Russian and Russian-Shor dictionary] (Kemerovskoje knižnoje izdatel'stvo, 1933).
Lessing, F. *Mongolian-English dictionary* (University of California Press, 1960).
Luwsandendew, A & Tsedendamba, Ts. *Bol'šoj akademičeskij mongol'sko-russkij slovar'* [Big Russian-Mongol dictionary] *Vols. 1–4* (Academia, 2001-2002).
Malov, S. Je. *Jazyk želtyx ujgurov* [The language of the Yellow Uyghurs] (Izdatel'stvo Akademii nauk Kazaxskoj SSR, 1957).
Mostaert, A. *Dictionnaire Ordos* (Johnson Reprint Company, 1968).
Muniev, B. D. Kalmycko-*russkij slovar'* [Kalmyk-Russian dictionary] (Russkij jazyk, 1977).
Nadžip, E. N. *Ujgursko-russkij slovar'* [Uyghur-Russian dictionary] (Sovetskaja enciklopedija, 1968).
Nugteren, H. *Mongolic Phonology and the Qinghai-Gansu Languages* (Utrecht University, 2011).
Osmanov, M. M. et al. *Tatarsko-russkij slovar'* [Tatar-Russian dictionary] (Sovetskaja enciklopedija, 1966).
Rassadin, V. I. *Tofalarsko-russkij i russko-tofalarskij slovar'* [Tofa-Russian and Russian-Tofa dictionary] (Drofa, 2005).
Róna-Tas, A. & Berta, Á. West Old Turkic: Loanwords in Hungarian. *Turcologica* ***84***, Vol. 1–2 (2011).
Sauranbaev, N. T. *Russko-kazaxskij slovar'* [Russian-Kazakh dictionary] (Gosudarstvennoe izdatel'stvo inostrannyx i nacional'nyx slovarej, 1954).
Sevortjan, E.V. *Etimologičeskij slovar' tjurkskix jazykov. Obščetjurkskije i mežtjurkskije osnovy na glasnyje* [An Etymological dictionary of the Turkic languages. Common vowel-initial stems] (Nauka, 1974).
Sevortjan, E. V. Etimologičeskij slovar' tjurkskix jazykov. Obščetjurkskije i mežtjurkskije osnovy na bukvu B [An Etymological dictionary of the Turkic languages. Common b-initial stems] (Nauka, 1978).
Sevortjan, E. V. *Etimologičeskij slovar' tjurkskix jazykov. Obščetjurkskije i mežtjurkskije osnovy na bukvy V, G i D* [An Etymological dictionary of the Turkic languages. Common *v*-, *g*- and *d*-initial stems] (Nauka, 1980).
Sleptsov, P. A. *Jakutsko-russkij slovar'* [Yakut-Russian dictionary] (Sovetskaja enciklopedija, 1972).
de Smedt, A., & Mostaert, A. *Dictionnaire monguor-francais* (Imprimerie de l'Université Catholique, 1933).
Stachowski, M. *Dolganischer Wortschatz* (Uniwersytet Jagielloński, 1993).
Subrakova, O. V. *Xakassko-russkij slovar'* [Khakas-Russian dictionary] (Nauka, 2006).
Tağiyev, M. T. et al. *Azərbaycanca- Rusca lüğət* [Azerbaijani-Russian dictionary] (2006).
Tenišev, E. R. *Stroj salarskogo jazyka* [The system of Salar] (Nauka, 1976).
Tenishev, E. R. et al. *Sravnitel'no-istoričeskaja grammatika tjurkskix jazykov. Leksika* [A historical comparative grammar of the Turkic languages. Lexicon] *2nd edn.* (Nauka, 2001).
Tenišev, E. R. *Tuvinsko-russkij slovar'* [Tuvan-Russian dictionary] (Sovetskaja enciklopedija, 1968).
Tenišev, E. R. & Sujunčev, X. I. *Karačajevo-balkarsko-russkij slovar'* [Karachay-Balkar-Russian dictionary] (Russkij jazyk, 1989).
Tumurdej, G. & Tsybenov, B. D. *Dagursko-russkij slovar'* [Dagur-Russian dictionary] (Izdatel'stvo BNTs SO RAN, 2014).
Uraksin, Z. G. *Baškirsko-russkij slovar'* [Bashkir-Russian dictionary] (Digora, Russkij jazyk, 1996).
Useinov, S. M. *Russko-krymskotatarskij, krymskotatarsko-russkij slovar'* [Russian-Crimean Tatar, Crimean Tatar-Russian dictionary] (Tezis, 2007).
Verbitskij, V. I. *Slovar' altajskogo i aladagskogo narečij tjurkskogo jazyka* (Izdanie Pravoslavnogo Missionerskogo Obščestva, 1884).

### 2 Archaeological reanalysis.

#### 2.1. Missing data

Tian et al. raise two specific objections with respect to our archaeological database and to one of the analyses performed thereon. The first issue raised is of 'missing data'. While missing data can be a problem for testing certain specific hypotheses, our archaeological database was designed to collate a broad set of available information on Neolithic and Bronze Age assemblages from Northeast Asia. As explained in our Article (SI 7), this was regarded as the most practical way to reduce bias from differing standards of sampling, analysis and publication within the region. In most cases we scored large settlement sites in order to reduce variability from inter-site function, a method which also increases the utility of the data for phylogenetic analysis[34]. Tian et al. correctly note that we scored 171 archaeological features not 172.

Tian and colleagues attach a file which re-scores 'missing data' for selected categories from one area of our database (northern China). They write that they replaced 2184 'missing values' (we count 2192). It is not explained how this re-scoring was performed. We assume the procedure was to change 'absent' to 'missing' when the authors could find no published information on a certain *category* of artefacts or features. However, examination of their data shows numerous inconsistencies. Out of the 109 sites which were re-scored, at least 61 are problematic. Two main types of inconsistency present themselves in the re-scoring. In one type, one or members of a certain category are scored as 'present', but other members are re-scored as 'missing'. In the other type, while a certain category was originally scored as 'absent', this has been changed to 'missing' for some but not all sites concerned.

With respect to burials, while the authors state that 'For all sites where no burials were found, burial features were incorrectly coded as 0', at two sites (Nantaizi and Chengzishan Xiajiadian F2) one type of grave is reported present but others are re-classified as missing rather than absent. Furthermore, there are 25 sites where burials were originally reported as 'absent' but which have not been changed to 'missing'. It is unclear if this is an over-sight or if there is any archaeological basis for the differing evaluations.

Out of 40 sites re-scored with 'missing data' for flora and fauna, six have some reported plants or animals. Given that standard practice in both zooarchaeology and archaeobotany is only to report species that are identified from a site, we are unsure why unreported species are re-classified as missing rather than absent. In 36 cases, fish remains are accepted as absent or present but land animals are scored as 'missing'. In terms of faunal taphonomy, the preservation of fish bones suggests that animal remains would also be preserved if originally present at the site. For Shuangtuozi phase III, reported plant remains comprise rice and both foxtail and broomcorn millet; the basis for re-classifying other plants as missing rather than absent is therefore puzzling. At Xinglongwa, several plants (rice, wheat, barley, adzuki, *Perilla*, walnut and peach) are re-scored as missing, yet others (foxtail and broomcorn millet,

---

[34] Prentiss, A.M. et al. Cultural macroevolution in the middle to late Holocene Arctic of east Siberia and north America. *J. Anthropol. Archaeol.* **65**, 101388 (2022).

soybean, Cannabis and acorn) are present in our data. At Haminmangha, while pig, rabbit, horse, chicken, deer, foxtail millet, broomcorn millet and Cannabis are scored as present, all other plants and animals have been re-scored as 'missing' rather than absent. At Xiaozhushan Phase V, pig and dog are present but other animals re-scored as missing. At Zuojiashan Phase II, wild boar, sheep and deer were present but other animals re-scored as missing; at Phase III of the same site, deer was present but again other animals are re-scored as missing.

At Shuangtuozi phase II, the excavation of five storage pits suggests that other architectural features such as pit houses or ditches might also be identified if present. The basis for classifying other architectural features such as pit houses as 'missing' is therefore unclear but seven other assemblages are similarly re-classified as having missing data for some architectural features even though others are present: Dazuizi I and II (post holes), Houtaomuga I and II (ditches), Shangzhai (pit houses and hearths), and Shangdong Zhaogezhuang and Yinjiacun I (storage pits). At Gaolichengshan, surface buildings are re-scored as missing even though pit dwellings, hearths and door/entrance features are all present.

While most of the re-scoring is conducted on food, architectural and mortuary remains, an unexplained outlier is Dalian Guojiacun (Upper) where 'stone sword' is re-scored as missing even though there is a large corpus of other stone tools.

Not only does their re-scoring include numerous false or inconsistent data, Tian et al.'s re-analyses were conducted with missing data included for only one sub-set of the database, i.e., 109 sites from northern China. The 146 sites from other regions retain our original present/absent scoring. The authors provide no explanation as to why they decided to re-score only one part of the data or why such a skewed procedure might produce meaningful results.

#### 2.2. Archaeological phylogeny

With respect to the phylogenetic analysis, Tian et al. note a discrepancy in the number of taxa between the Article and the XML file. They correctly surmise that the 4 extra taxa in the latter were retained from an earlier version of the database. These are cases where scoring of the same site was initially duplicated and the results later combined. We have now provided the updated XML file under https://github.com/rbouckaert/Eurasia3angle/releases/download/v1.0/culturexml.zip

Nevertheless, the correct outcome of our cultural tree was provided in our Article in "Extended data Fig. 2. Bayesian phylogenetic analysis of the archaeological database" and in "SI 8 Bayesian cultural interpretation". Thus, although the results of our phylogenetic analysis were available to the reader, the encoding of the computational process was not. The inclusion of the prefinal XML file is unlikely to have made a major difference but we apologize that the authors have been hampered in the replication of our results.

Tian et al. base their critique of the phylogenetic analysis of our archaeological database upon methodological claims not supported in the literature, as well as confusing statements such as our 'analysis assumes that a phylogenetic tree is a good representation of the history of the archaeological sites.' The 'history of archaeological sites' would usually refer to taphonomic processes affecting a particular accumulation of cultural and natural remains over time. Our phylogenetic analysis was certainly not intended to explain site taphonomy, rather to model the history of human groups (proxied by archaeological assemblages and cultures). More broadly, Tian and colleagues insist that a single tree showing vertical inheritance is required to support our conclusions. This contradicts research, including that by one member of Tian's team, that real-world examples of historical evolution do not always fit the classic tree-like pattern; or, in other words, that certain levels of horizontal transmission do not invalidate the phylogenetic method[35,36]. One such study specifically argues that 'realistic levels of reticulation between cultures do not invalidate a phylogenetic approach to cultural and linguistic evolution.'[37]

In contrast to this previous research, Tian and colleagues regard the phylogenetic analysis of our archaeological database as a decisive test of the archaeological evidence for Transeurasian. This represents a narrow perspective which makes no attempt to engage with the broader archaeological literature. Our original standpoint was more circumspect, employing phylogenetics to compare cultural groupings and their estimated chronologies with classical archaeological methods. In our Article (SI 8), we noted that similarity in phylogenetic analyses can result from factors such as areal diffusion due to geographical proximity, chronological simultaneity and cultural continuity through inheritance from

---

[35] Gray, R.D. & Jordan, F.M. Language trees support the express-train sequence of Austronesian expansion. *Nature* **405**, 1052-1055 (2000).

[36] Currie, T.E., Greenhill, S.J., Mace, R. Is horizontal transmission really a problem for phylogenetic comparative methods? A simulation study using continuous cultural traits. *Phil. Trans. R. Soc. B* **365**, 3903-3912 (2010).

[37] Greenhill, S.J., Currie, T.E., Gray, R.D. Does horizontal transmission invalidate cultural phylogenies? *Proc. R. Soc. B* **276**, 2299-2306 (2009).

previous generations. We recognised that the features in our archaeological dataset were not necessarily always transmitted through vertical inheritance but could also result from social contact between one culture and its neighbour. Our phylogenetic tree thus captures not only a historical signal of descent of these cultures, but also a signal of interaction.

Tian et al. further object that “only a small number of clades have support above 50%. This aspect of the posterior was not evident in the original study, as Robbeets et al. only produced a Maximum Clade Credibility (MCC) tree.” While the MCC tree displayed in our Article does not have posterior support values shown, they are available in the MCC tree in the Supplementary Information (SI 25 cultural phylogeny.eps). We acknowledge that there is low clade support for many clades, but note that clade support is easily broken up by rogue taxa, and the overall structure still seems reasonable.

Tian et al. claim that phylogenetic analysis of archaeological data requires *all* artefacts and features to evolve in tandem. This claim is not supported by any discussion of how archaeological remains relate to cultural lineages and ignores the caveats presented in our Article (SI 7). It also ignores recent research stressing the need for careful consideration of the extent to which different cultural (including linguistic) traits might have different evolutionary histories[38,39]. Although Tian et al.’s suggestion of inferring trees inside trees is appealing, it is computationally not yet feasible because computational time to reach convergence will be too long. At the moment, StarBeast3 is the fastest implementation of the multi-species coalescent (i.e. a trees inside tree method)[40] but is not able to handle the amount of data if we assume every item has its own history.

By contrast, we considered some examples of different sub-sets of features in our Article (SI 7). Our database was scored from published assemblages associated in the literature with archaeological cultures. The resulting phylogenetic analysis grouped those cultures in a way largely consistent with previous research, but it cannot be assumed that separate features would all necessarily group in the same way. For example, building a tree based on only faunal and floral remains is unlikely to mirror cultural groupings since farmers continued to

[38] Greenhill, S.J., Atkinson, Q.D., Meade, A., Gray, R.D. The shape and tempo of language evolution. Proc. R. Soc. B. 277, 2443-2450 (2010).
[39] Evans, C.L., Greenhill, S.J., Watts, J., List, J-M., Botero, C.A., Gray, R.D., Kirby, K.R. The uses and abuses of tree thinking in cultural evolution. Phil. Trans. R. Soc. B 376, 20200056 (2021).
[40] Douglas J, Jiménez-Silva CL, Bouckaert R. StarBeast3. Adaptive parallelized Bayesian inference under the multispecies coalescent. Systematic Biology **71**, 901-916. (2022).

hunt and gather wild resources. Combining food remains with burials and shell and bone artefacts in a separate tree is likely to only increase uncertainty in the analysis.

In short, the results of our phylogenetic analysis of archaeological cultures were broadly consistent with previous archaeological analyses using traditional typological and other classical methods. Given our acknowledgement of the role of horizontal transmission, we do not assume that all archaeological cultures in the area of analysis derived from a single common ancestor in a strict sense. Nevertheless, we believe our phylogenetic analysis provides further support for a large body of previous archaeological literature which models agricultural expansions from northeast China into Manchuria, the Primorye, Korea, Japan and the Ryukyu Islands.

#### 2.3. Population movements

In concluding their section on archaeology, Tian and colleagues claim that there is no archaeological evidence supporting the population movements discussed in our Article. This sweeping claim goes completely against the existing literature yet no archaeological evidence is provided in support. In fact, the archaeological sections of Tian et al.'s Commentary and SI cite no archaeological references at all, nor do they engage with the archaeological record in any way. In contrast to what readers might assume from the Comment, there is considerable archaeological evidence for population movements associated with agriculture in Neolithic and Bronze Age Northeast Asia. While debate has revolved around the *scale*, *timing* and *routes* of these migrations, it is widely accepted that Neolithic farming populations moved from northeast China to the Primorye and the Korean peninsula and that Bronze Age groups then spread from Korea to Japan. With respect to the *scale* of these dispersals, our publication of the first series of human genomes from Yayoi Kyushu and from the early modern Ryukyus supports large-scale immigration in association with the spread of farming in the Japanese Islands. Our Article did not present direct evidence on the scale of the other migrations we discussed.

Table 12 shows four stages of Neolithic/Bronze Age farming dispersals in Northeast Asia based on archaeobotanical and zooarchaeological remains of domesticated plants and animals. Such remains provide the most direct evidence of the spread of agropastoral systems and are analysed in our Article using the largest database of radiocarbon dated cereals from

Northeast Asia so far published. Well-dated evidence from domesticated plants and animals is central to any discussion of farming dispersals (and possible linguistic correlations)[41] and underpins the framework tested in our Article. We find it curious that Tian et al. have nothing to say about this key aspect of Northeast Asian prehistory.

**Table 12.** Key farming dispersals in Neolithic/Bronze Age Northeast Asia discussed in our Article but rejected in Tian et al.'s Comment.

| **Farming dispersal** | **Chronology** | **Archaeological support** (in addition to archaeobotanical and zooarchaeological remains) |
|---|---|---|
| Millet farming to the Primorye and Korea | Middle Neolithic: fourth millennium BC | **(1) Primorye**. Stone tools: grinding slabs, pestles, constricted-waist hoes; cord-marked pottery; spindle whorls. (2) **Korea**. Comb-pattern pottery; stone tools (grinding stones, pestles, waisted hoes, polished arrowheads); spindle whorls. |
| Spread of rice cultivation into northern China (Shandong and Liaodong) | Middle-Late Neolithic: 4000 - 1500 BC | Polished stone reaping knives; wet rice technology |
| Dispersal of rice farming to the Korean peninsula; later (?) arrival of barley and wheat | Bronze Age (Mumun): ca. 1500 – 150 BC | Ceramic analyses suggest combination of influences from both Liaodong and the West Liao basin. Stone agricultural tools from the Bohai Sea tradition. Later (Early-Middle Bronze Age) bronzeware influences from Liaoning. |
| Spread of cereal farming (millets, rice, barley & wheat) from Korea to Japan | Bronze Age (Yayoi): 1000 BC – AD 250 | Extensive evidence from ceramics, lithics, burial customs, circular moated settlements, pit house architecture |

Our analysis of directly-dated cereal remains from Northeast Asia confirmed and extended previous research[41,42,43]. The discontinuous spread of cereals in the region shows that agriculture did not expand in a single, progressive wave of advance. Instead, there were numerous branch-like movements deriving from a complex range of factors including demography, climate and socioeconomic change associated with emerging Bronze Age political complexity.

---

[41] Stevens, C. & Fuller, D.Q. The spread of agriculture in eastern Asia: archaeological bases for hypothetical farmer/language dispersals. *Lang. Dyn. Chang*. **7**, 152-186 (2017).
[42] Leipe, C. et al. Discontinuous spread of millet agriculture in eastern Asia and prehistoric population dynamics. *Sci. Adv*. **5**, eaax6225 (2019).
[43] Leipe, C. et al. The spread of rice to Japan: insights from Bayesian analysis of direct radiocarbon dates and population dynamics in East Asia. *Quat. Sci. Rev*. **244**, 106507 (2020).

Earlier Neolithic millet dispersals in East Asia were likely conducted by low-density cultivators[41,44]. This scenario may be applicable to the Neolithic Primorye and Korea, although research in northeast China has found an exponential increase in population associated with millet farming[42]. Tian et al. argue that our phylogenetic analysis of archaeological assemblages does not support a Middle Neolithic expansion of millet farmers via two largely separate movements to the Primorye and Korea. It is these two movements which we propose can be triangulated with the initial branching of Proto-Tungusic and Proto-Koreanic, respectively. In the early 1990s, it was proposed that millet cultivation had first crossed Liaodong to northern Korea and only then moved north to the Primorye[45]. Over the past two decades, however, most specialists have supported separate movements to the Primorye and Korea[46], although debate continues over the exact routes involved[47]. Middle Neolithic coastal connections between Liaodong and southern Korea continue to be emphasised by specialists[48,49]. Links between Manchuria and the Primorye have been further confirmed as a result of excavations by a joint Russo-Japanese team in the early 2000s. New ceramic analyses, for example, have shown that the Khanxi 1 type cord-marked pottery, which can be dated to the earliest stage of the Zaisanovka culture, is similar to ceramics described from the Yabuli site located in the Mudanjiang valley in Heilongjiang province[50]. A recent detailed study of early millet dispersals in Northeast Asia concluded that the currently available archaeological evidence supports the bifurcated model of millet dispersals[51].

The scale of population movement associated with the spread of millet to Neolithic Korea has been controversial, in part because it has become linked with broader debates over

[44] Qin, L., Fuller, D.Q. Why rice farmers don't sail: coastal subsistence traditions and maritime trends in early China, in *Prehistoric Maritime Cultures and Seafaring in East Asia* (eds Wu, C. & Rolett, B.) 159-191 (Springer, 2019).
[45] Yan, W. Dongbeiya nongye de fasheng yu chuanbo [Beginning and dispersal of agriculture in Northeast Asia]. *Nongye Kaogu* **3**, 37-44 (1993).
[46] Kuzmin, Y.V. The beginnings of prehistoric agriculture in the Russian Far East: current evidence and concepts. Documenta Praehist. 40, 1-12 (2013).
[47] Vostretsov, Y.E. The interaction of marine and agricultural adaptations in the basin of the Sea of Japan, in *The Russian Far East in Prehistory and the Middle Ages* (ed Andreeva, Z.V.) 159-186 (Vladivostok: Dalnauka, 2005) (in Russian).
[48] Nelson, S.M. *The Archaeology of Korea* (Cambridge Univ. Press, 1993).
[49] Lee, G.A. The spread of domesticated plant resources in prehistoric northeast Asia, in *Routledge Handbook of Archaeology and Globalization* (ed Hodos, T.) 394-412 (Routledge, 2017).
[50] Miyamoto, K. The initial spread of early agriculture into Northeast Asia. *Asian Archaeol.* **3**, 1-12 (2014).
[51] Li, T. et al. Millet agriculture, dispersed from Northeast China to the Russian Far East: integrating archaeology, genetics and linguistics. *Archaeol. Res. Asia* **22**, 100177 (2020).

Korean national origins[52]. Korean archaeologists often assume that millet reached Korea through small-scale cultural diffusion from northeast China rather than by actual population movements[53]. This has been proposed on the basis that there were no major transformations in material culture or settlement patterns, that hunter-gathering remained the primary subsistence adaptation, and that there was little or no demographic growth as a result of the adoption of millet cultivation[54,55,56]. In a recent re-evaluation of these assumptions, we argued that the archaeological record is not inconsistent with population dispersals at this time[57]. While Miyamoto also critiques the Chulmun Proto-Koreanic dispersal theory on the grounds of ceramic typology[58], he notes the presence of northern Chinese style agricultural stone tools, such as mortars, pestles and hoes, in association with early millet sites in northwest Korea[50]. Similar use wear patterns on these tools to those found in Early and Middle Neolithic sites in northern China links these tools with millet processing[59,60]. If cultural diffusion alone was behind the spread of millet to Neolithic Korea, we might expect the same crops to have been taken up in Jōmon Japan, where the subsistence economy was otherwise similar to the Chulmun. There is, however, no evidence of millet in Japan until the first millennium BC, even though the two regions were in close contact[61,62]. This suggests that

52 Park, H.W. & Wee, K. The nationalistic trend in South Korean archaeology: documenting the development of a unilinear evolutionary trajectory of a homogenous Korean peoples. *Archaeologies* **12**, 304-339 (2016).

53 Kim, W.Y. *Art and Archaeology of Ancient Korea* (Taekwang, 1986).

54 Ahn, S.M. The emergence of rice agriculture in Korea: archaeobotanical perspectives. *Archaeol. Anthropol. Sci*. **2**, 89-98 (2010).

55 Kim, J.S. & Park, J. Millets vs. rice: an evaluation of the farming/language dispersal hypothesis in the Korean context. *Evol. Hum. Sci*. **2**, e12 (2020).

56 Kim, J.S. & Seog, C.T. Maritime prehistory of Korea: an archaeological review, in Maritime Prehistory of Northeast Asia (eds Cassidy, J., Ponkratova, I. & Fitzhugh, B.) 29-50 (Springer, 2022).

57 Hudson, M.J. & Robbeets, M. Archaeolinguistic evidence for the farming/language dispersal of Koreanic. *Evol. Hum. Sci*. **2**, e52 (2020).

58 Miyamoto, K. The emergence of 'Transeurasian' language families in Northeast Asia as viewed from archaeological evidence. *Evol. Hum. Sci*. **4**, e3 (2022).

59 Kamijō, N. Chōsen hantō senshi jidai no maban, mabō ni okeru shiyōkon bunseki [Use wear analysis of stone pestles and mortars from the prehistoric Korean peninsula], in Nihon suitō nōkō no kigenchi ni kansuru sōgōteki kenkyū [Interdisciplinary Research on the Origin of Rice Agriculture in Japan] (ed Miyamoto, K.) 87-104 (Fukuoka: Kyushu Univ., 2008).

60 Kamijō, N. Shandong ban dao mo pan yu mo bang de shi yong wei hen ji dian fen li fen xi [Use wear and starch analysis of stone pestles and mortars from the Shandong peninsula], in *Zao qi nong ye he ren lei xue yan jiu* [Anthropology and the Early Agriculture of the Shandong Peninsula] (eds Luan, F. & Miyamoto, K.) 149-186 (Peking: Ke xue chu ban she, 2008).

61 Bausch, I. Prehistoric networks across the Korea strait (5000-1000 BCE): 'Early globalization' during the Jomon period in northwest Kyushu?, in *Routledge Handbook of Archaeology and Globalization* (ed Hodos, T.) 413-437 (Routledge, 2017).

62 Hudson, M.J., Bausch, I.R., Robbeets, M., Li, T., White, J.A. & Gilaizeau, L. Bronze Age globalisation and later Jōmon social change. *J. World Prehist.* **34**, 121-158 (2021).

demic expansion was a significant factor behind the spread of millets into Korea but that the same populations did not spread to Japan.

Even if population movements into Neolithic Korea were on a relatively small-scale, language shift does not necessarily require a large influx of people. A new technological or social adaptation can provide sufficient benefits for the spread of a particular language[63]. The farming/language dispersal hypothesis argues that agriculture enables higher production and storage of food per area of land[64]. Even a relatively minor emphasis on millet cultivation in a foraging context would thus likely have generated social conditions conducive to the spread of the language concerned. Furthermore, alternative proposals for the arrival of Proto-Koreanic on the peninsula after the Neolithic come up against various linguistic problems. In terms of archaeology, linking Bronze Age farming with the arrival of *both* Macro-Koreanic and Japanic—as suggested by Kim and Park[55]—raises the problem of how two different ethnolinguistic groups might have originated from the materially homogenous Early Mumun horizon.

After around 4000 BC, both millet and rice agriculture began to spread into new ecological zones within China and these cereals often became components of the same farming economy[41,65]. While earlier rice has been reported from Shandong[66], its domesticated status is disputed and the expansion and integration of rice cultivation into the economies of the Lower Yellow River and Bohai Sea areas appears to have grown primarily within the context of the Late Neolithic and Bronze Age[41,67]. Wheat and barley were added to this agricultural economy after 2500 BC and it was this diverse or 'globalised' cereal farming that then spread to Korea and Japan between ca. 1500-1000 BC[68]. Archaeobotanical evidence of

---

early rice from Liaodong is not extensive but new discoveries are growing[69,70,71,72]. It is broadly accepted that rice spread to Korea from the Bohai Sea area through population movement[43,49,50,54].

The phylogenetic analysis of our archaeological database clustered Bronze Age assemblages from Korea and Japan with Xiajiadian rather than Liaodong-Shandong. By contrast, Tian et al.'s tree found 'no evidence in favour of a close relationship [between Bronze Age Korea and Japan] with Bronze age West Liao river sites.' This leads the authors to reject what they claim is our argument 'for a Bronze Age migration from the West Liao river and Shandong to Korea and Japan (Mumun and Yayoi cultures) by 3500 BP that would correspond to the spread of the Japonic family.' This statement misinterprets our results and obfuscates the issues on several levels. We did not propose a direct migration from the West Liao to Korea and Japan, instead noting that the clustering 'may be explained by cultural interaction with Xiajiadian groups in the Bronze Age, when the ancestors of the Mumun and Yayoi were living on the coasts between Liaodong and Shandong, neighbouring the Xiajiadian in the north' (SI 8). Archaeological evidence suggests that Bronze Age coastal farmers living around the Bohai Sea migrated to Korea and then Japan. However, the Liao river region was of key importance through its location at the edge of the eastern steppes from where numerous new cultural elements such as domesticated plants and animals, bronze working and weapons entered East Asia[68],[73,74,75]. The Bronze Age history of Northeast Asia cannot be discussed in isolation from the interplay between the interior Liao basin and coastal Liaodong regions.

Given a long history of research, the Bronze Age migration of farmers from the Korean peninsula to the Japanese archipelago is by far the best understood dispersal of those listed in Table 12. Although the new tree generated by Tian et al. clearly supports a Mumun/Yayoi clade, the authors still insist that the 'inferred histories does [sic] not correspond to the expected output, nor to the conclusions described by Robbeets et al.' The reluctance of Tian

---

[69] d'Alpoim Guedes, J., Jin, G. & Bocinsky, R.K. The impact of climate on the spread of rice to north-eastern China: a new look at the data from Shandong province. *PLoS ONE* **10**, e0130430 (2015).
[70] Ma, Y., Wu, W., Wang, Q., Zhang, Z. & Jin, G. Dalian Wangjiacun yizhi danhua yicun yanjiu. *Beifang Wenwu* **2**, 39-43 (2015).
[71] Obata, H., Saito, N. & Wang, Q. Wangjiacun yizhi chute taoqi yahen fenxi de chengguo he wenti. *Dongfang Kaogu* **27**, 1-20 (2018).
[72] Obata, H. in *Scientific research on the process of the spread of agriculture in Northeast Asia by archaeobotany* (Miyamoto, K. ed), 38-60 (Kyushu Univ., 2019)
[73] Rawson, J. China and the steppe: reception and resistance. *Antiquity* **91**, 375-388 (2017).
[74] Kobayashi, S. Bronze cultures around eastern Eurasia at 1st millennium BC and the origin of Yayoi bronzeware. *Bull. Nat. Mus. Jpn. Hist.* **185**, 213-238 (2014) (Japanese with English summary).
[75] Hudson, M. *Bronze Age Maritime and Warrior Dynamics in Island East Asia* (Cambridge Univ. Press, 2022).

and colleagues to accept an association between language and farming dispersals for the Japonic branch of Transeurasian even when supported by their own evidence suggests that their objections to our Article owe more to partisanship than to objective science.

In conclusion, as regards archaeology, Tian and colleagues display a remarkable lack of interest in the relevant archaeological sequences from Neolithic and Bronze Age Northeast Asia. Their Comment is limited to technical issues over the application of one analysis within our archaeological section and no specific critique is raised with our other archaeological conclusions, which provide extensive supporting evidence for the overall narrative. Nevertheless, our Article stressed the need for further analyses of our archaeological data set. We are pleased that Tian et al. have been able to use our publication to add further details and hope that other researchers will continue to test and refine the results.

**3 Genetic reanalysis**

Tian et al. object that we "associated the spread of farming to Korea with different waves of Amur and Yellow River gene flow, modelled by Hongshan for the Neolithic introduction of millet farming and Upper Xiajiadian for the Bronze Age addition of rice agriculture. This is, however, contradicted by the authors' own statement in the supplementary material that their data lack the resolution to distinguish between competing admixture models (SI 13 in Robbeets et al.)." More specifically, they question our modeling of Neolithic individuals with Hongshan and Bronze Age ones with Upper Xiajiadian, claiming that "[t]he populations they selectively assigned with Upper Xiajiadian ancestry in their Figure 3 could also be explained by using Hongshan instead of Upper Xiajiadian as a genetic source". To this end, they present a symmetry test in their Figure 2, which compares Hongshan and Upper Xiajiadian with a selection of our ancient Koreans and Japanese. This leads them to infer that Hongshan and Upper Xiajiadian are equally related to ancient Korean and Japanese ($|Z| < 3$). In other words, their genetic analysis indicates that Hongshan is an equally good proxy for the Bronze Age Korean and Japanese samples. If the Bronze Age Korean and Japanese samples can indeed be modeled with Hongshan, then — in Tian et al.'s reasoning— the association between the adoption of rice agriculture on the Korean Peninsula and the migration of rice farmers from the Shandong and Liaodong area suggested in our Article would be unjustified. Therefore, Tian et al. accuse us of "selective modelling of migration hypotheses", in other words, arguing that we cherry-picked Upper Xiajiadian as a model to intentionally alignthe eastward spread of rice agriculture with Bronze Age migrations to Korea and Japan.

First, our Article (SI 13) did not evade Tian et al.'s general observation that Upper Xiajiadian is not the only possible model for Bronze Age Korean and Japanese samples. In support of our previous findings about the lack of resolution to distinguish the precise source of East Asian ancestry in our data, we extended the symmetry tests performed by Tian et al. to a wider range of populations. In addition to Hongshan selected in Tian et al.'s analysis, we compared Upper Xiajiadian to a variety of other other ancient populations in the Amur River, West Liao River and Yellow River Basins. Our Figures (1-6) show that Bronze Age Upper Xiajiadian is not only indistinguishable from Middle Neolithic Hongshan (Fig. 2), but also from the Early Neolithic Jalainur in the Amur River Basin (Fig. 1), the Late Neolithic Lower Xiajiadian in the West Liao River Basin (Fig. 3), the Middle Neolithic Yangshao in the Yellow River Basin (Fig. 4), the Late Neolithic Longshan in the Yellow River Basin (Fig. 5) and Late Bronze/Iron Age individuals in the Yellow River Basin (Fig. 6). These results confirm the caveat made in our Article that the precise mainland East Asian ancestry source for ancient Korean and Japanese cannot be well distinguished by the current genetic data because of the low coverage of the ancient Korean and Japanese genome-wide data and the relative homogeneity of the ancient populations from the Amur, West Liao and Yellow River regions. If models cannot precisely distinguish which population is the best ancestry source for the target populations, the principle in population genetics is to select a population which is both in time and space close to the target population. We therefore modelled our Neolithic individuals with Hongshan and our Bronze Age ones with Upper Xiajiadian.

Even if our data lack the resolution to distinguish the precise source of mainland East Asian ancestry in the Bronze Age Korean data, they still allow us to associate the spread of farming to Korea with different waves of eastward gene flow from mainland East Asia in the Neolithic and the Bronze Age. In fact, our Article confirms previous research[76] that the difference between the Hongshan and Upper Xiajiadian genomes lies in the fact that Upper Xiajiadian had more Yellow River genetic contribution than Hongshan. The Middle Neolithic Hongshan (Banlashan 5400-5100 BP) had 60% Yellow River Ancestry, while it amounts up to 81% for the Upper Xiajiadian[76]. This gradually increasing Yellow river component, which is also observed in the Bronze Age Korean sample is associated with the adoption of rice farming on the Korean Peninsula. The ancient Korean individual from the Bronze Age (Taejungni) in our Article can be modeled entirely as mainland East Asian ancestry, i.e. an

[76] Ning, C. et al. Ancient genomes from northern China suggest links between subsistence changes and human migration. *Nat. Comm.* **11**, e2700.

admixture of higher proportions of Yellow River with lower proportions of Amur ancestry and lacking any Jomon component. As such, Upper_Xiajiadian serves as a better candidate than Hongshan (with larger p-value) even if they both provide an adequate model fit ($p > 0.05$). As for Bronze Age Japanese, it can be noted that new research confirms that they can be modelled as an admixture of Amur, Yellow River and Jomon genomes.[77]

Moreover, the association between farming and Bronze Age migration from northeast China to Korea is also compatible with extensive archaeobotanical research (see Section 2.3 above) about the spread of rice farming from the Liaodong-Shandong peninsula to Korea in the Bronze Age and linguistic research about the introduction of rice-vocabulary in Proto-Koreanic, presented in our Article (SI 5) as well as in previous research (Hudson & Robbeets)[57]. In sum, the areas of uncertainty noted by Tian et al. do not invalidate our proposed hypothesis for the formation of Bronze Age populations in Korea and Japan.

[77] Cooke, N. et al. Ancient genomics reveals tripartite origins of Japanese populations. *Science Advances* 7: eabh2419 (2021).

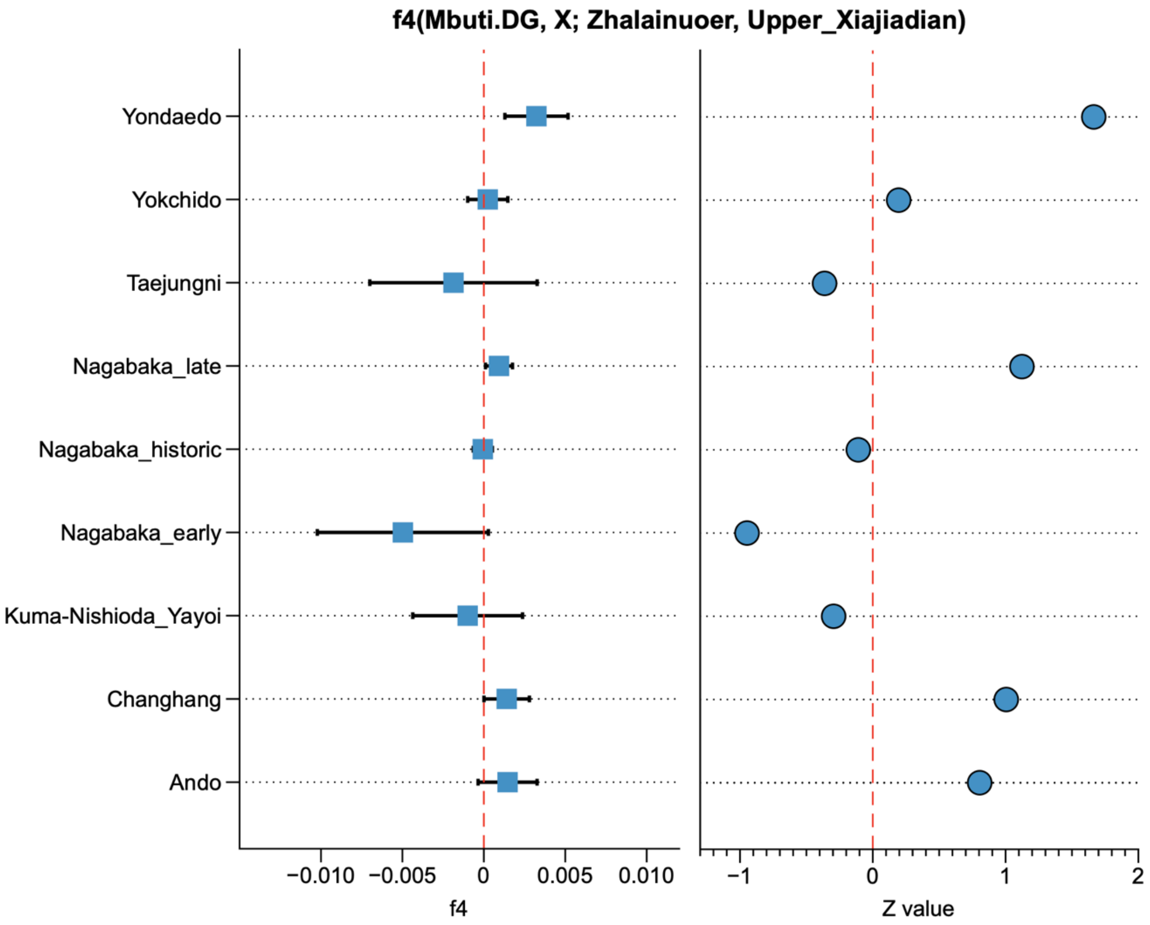

**Figure S1. The genetic difference between the Bronze Age Upper Xiajiadian in the West Liao River Basin and the Early Neolithic Jalainur in the Amur River Basin compared with the ancient Korean and Japanese individuals.** Horizontal bars represent the point estimate ± 1 s.e.m. s.e.m. are estimated using 5 cM block jackknifing. We find no significant difference ($|Z| < 3$) between the two populations with respect to the ancient Korean and Japanese individuals, suggesting that the low coverage of the data cannot provide enough resolution to distinguish who is a better proxy for the ancient Korean and Japanese populations.

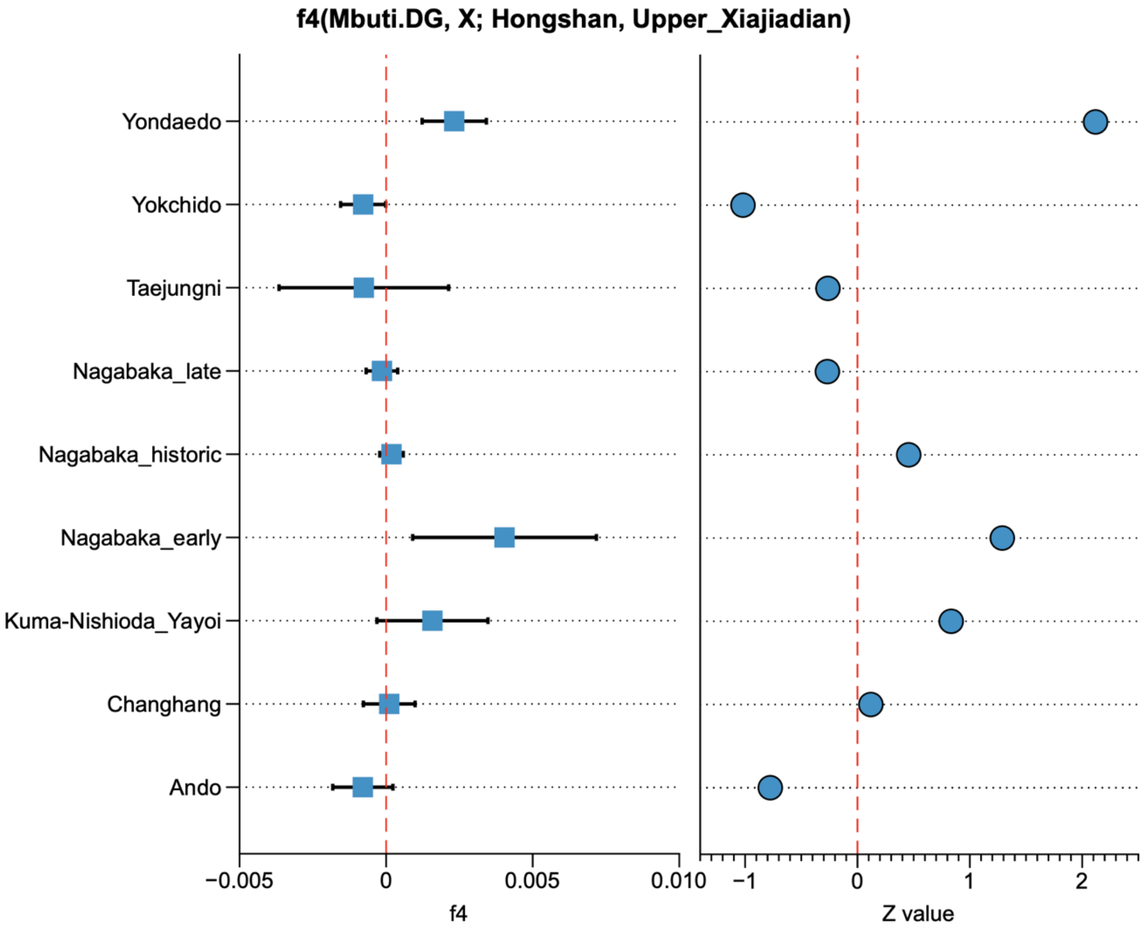

**Figure S2. The genetic difference between the Bronze Age Upper Xiajiadian and the Middle Neolithic Hongshan in the West Liao River Basin compared with the ancient Korean and Japanese individuals**. Horizontal bars represent the point estimate ± 1 s.e.m. s.e.m. are estimated using 5 cM block jackknifing. We find no significant difference ($|Z| < 3$) between the two populations with respect to the ancient Korean and Japanese individuals, suggesting that the low coverage of the data cannot provide enough resolution to distinguish who is a better proxy for the ancient Korean and Japanese populations.

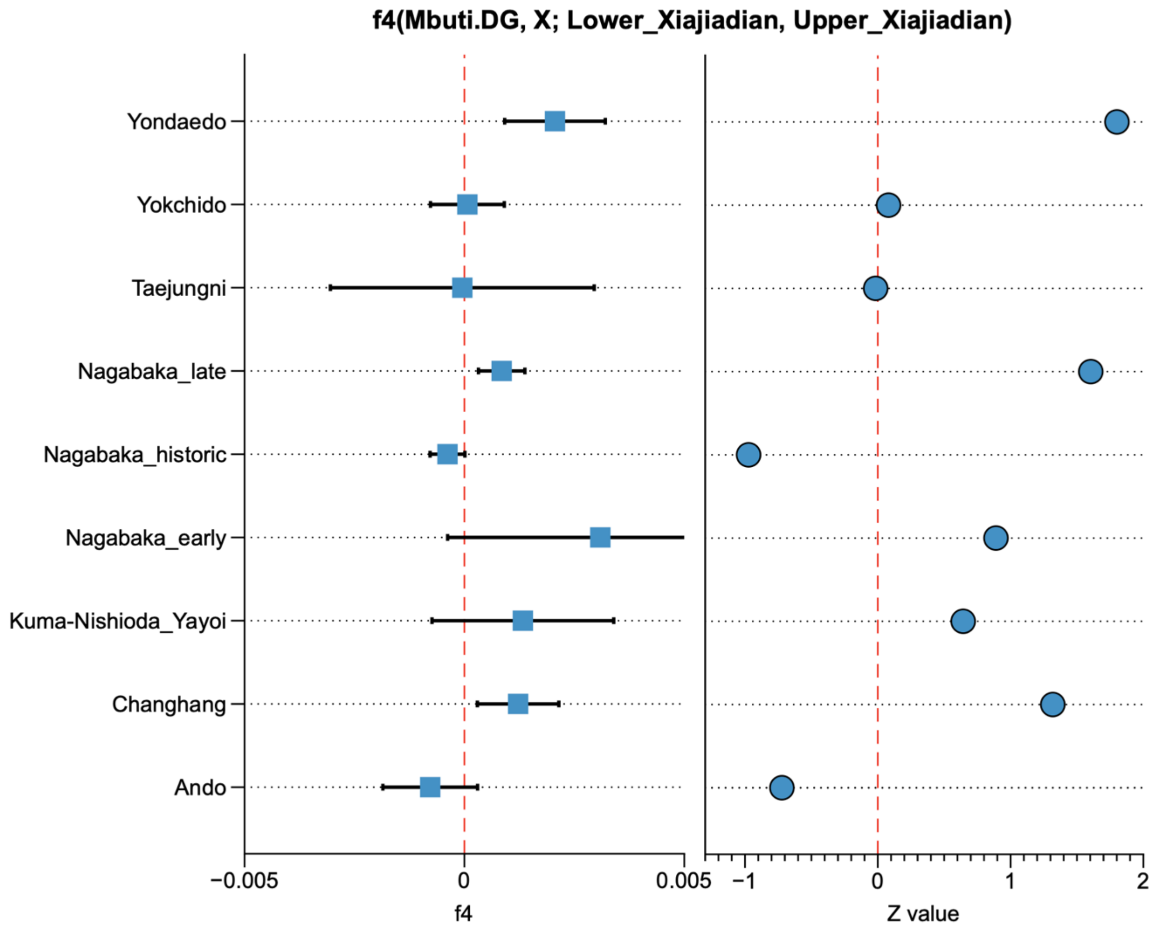

**Figure S3. The genetic difference between the Bronze Age Upper Xiajiadian and the Late Neolithic Lower Xiajiadian in the West Liao River Basin compared with the ancient Korean and Japanese individuals**. Horizontal bars represent the point estimate ± 1 s.e.m. s.e.m. are estimated using 5 cM block jackknifing. We find no significant difference ($|Z| < 3$) between the two populations with respect to the ancient Korean and Japanese individuals, suggesting that the low coverage of the data cannot provide enough resolution to distinguish who is a better proxy for the ancient Korean and Japanese populations.

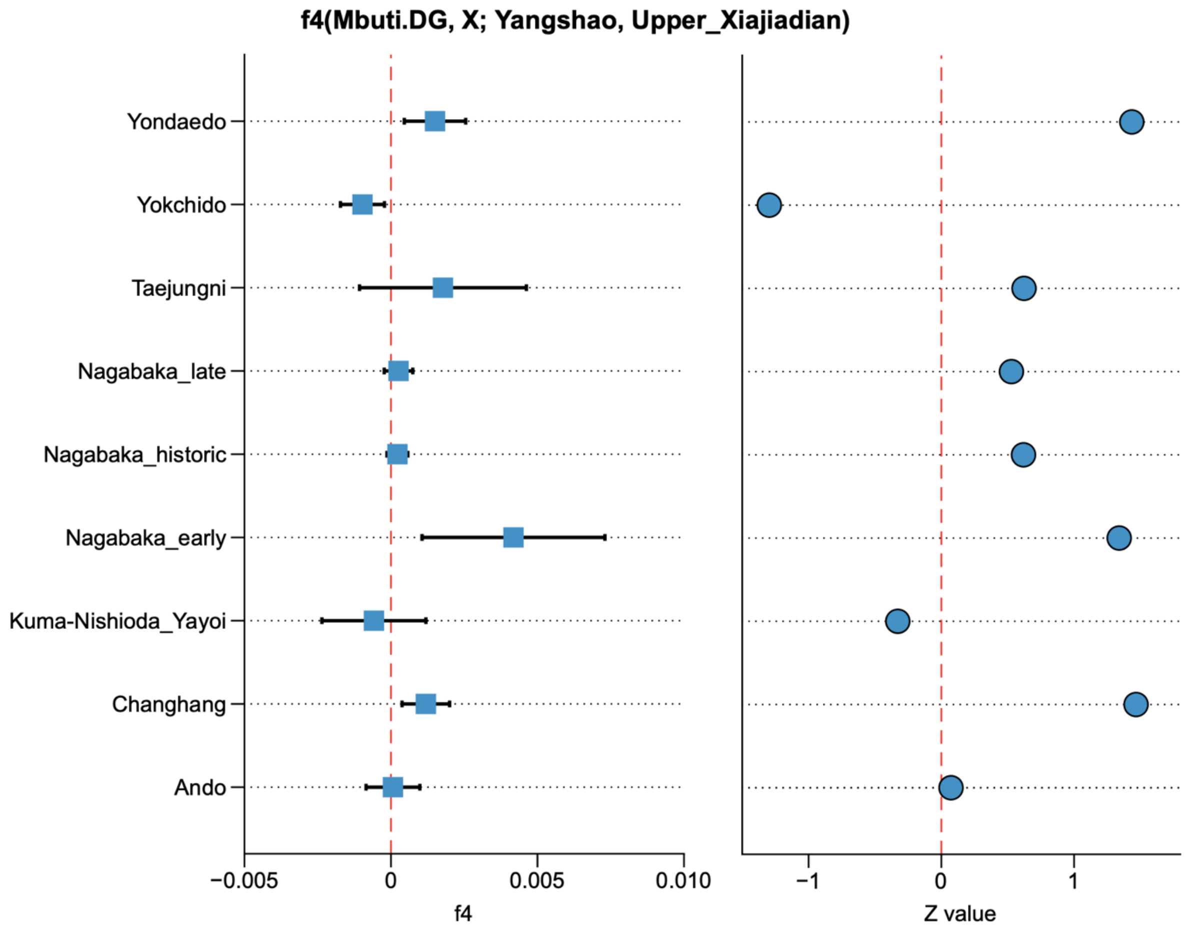

**Figure S4. The genetic difference between the Bronze Age Upper Xiajiadian in the West Liao River Basin and the Middle Neolithic Yangshao in the Yellow River Basin compared with the ancient Korean and Japanese individuals**. Horizontal bars represent the point estimate ± 1 s.e.m. s.e.m. are estimated using 5 cM block jackknifing. We find no significant difference ($|Z| < 3$) between the two populations with respect to the ancient Korean and Japanese individuals, suggesting that the low coverage of the data cannot provide enough resolution to distinguish who is a better proxy for the ancient Korean and Japanese populations.

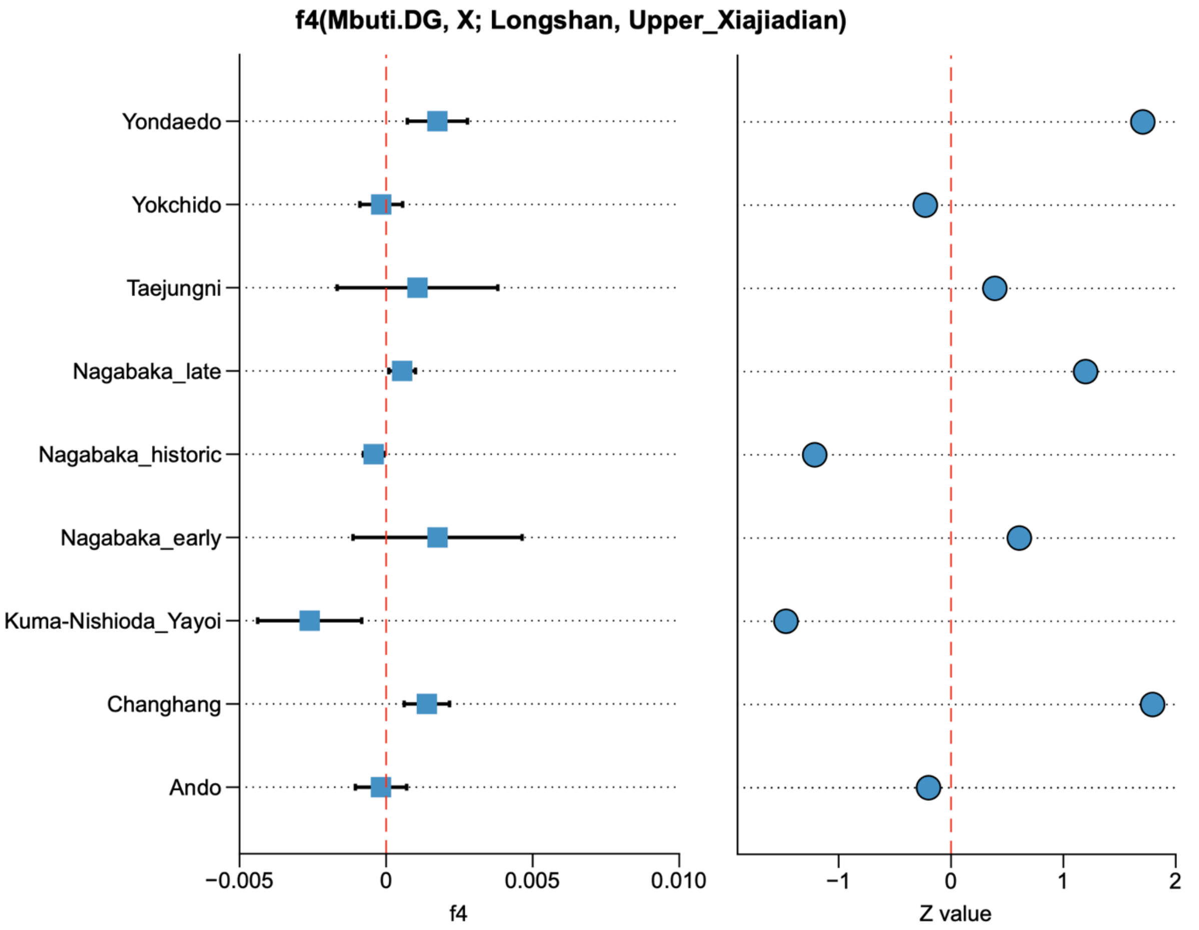

**Figure S5. The genetic difference between the Bronze Age Upper Xiajiadian in the West Liao River Basin and the Late Neolithic Longshan in the Yellow River Basin compared with the ancient Korean and Japanese individuals**. Horizontal bars represent the point estimate ± 1 s.e.m. s.e.m. are estimated using 5 cM block jackknifing. We find no significant difference ($|Z| < 3$) between the two populations with respect to the ancient Korean and Japanese individuals, suggesting that the low coverage of the data cannot provide enough resolution to distinguish who is a better proxy for the ancient Korean and Japanese populations.

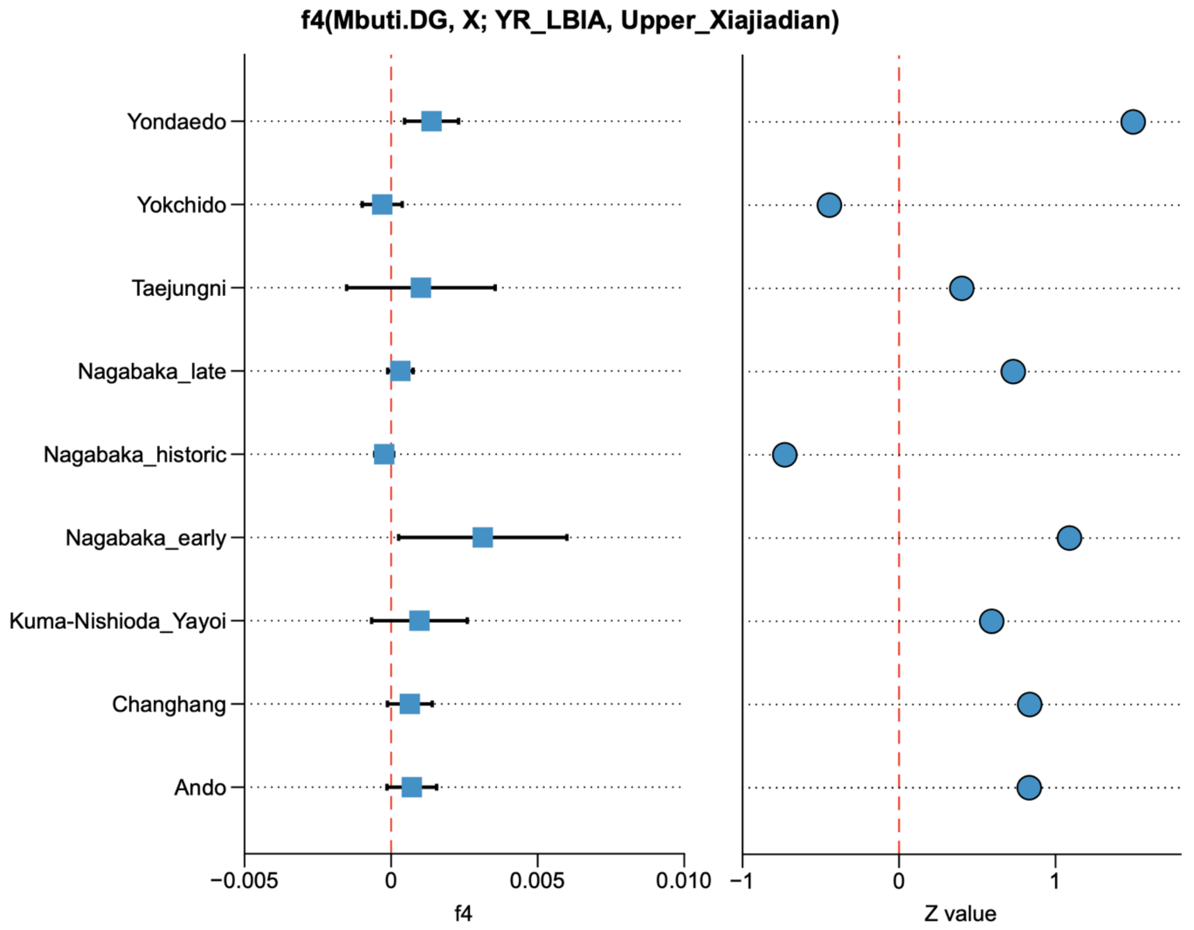

**Figure S6. The genetic difference between the Bronze Age Upper Xiajiadian in the West Liao River Basin and the Late Bronze/Iron Age individuals in the Yellow River Basin compared with the ancient Korean and Japanese individuals**. Horizontal bars represent the point estimate ± 1 s.e.m. s.e.m. are estimated using 5 cM block jackknifing. We find no significant difference (|Z| < 3) between the two populations with respect to the ancient Korean and Japanese individuals, suggesting that the low coverage of the data cannot provide enough resolution to distinguish who is a better proxy for the ancient Korean and Japanese populations.

### 4 Methodological revisit

#### 4.1 Overall conclusions

Above we have shown that Tian et al.'s re-evaluation of our linguistic, archaeological and genetic datasets contains multiple mis-scorings and misinterpretations that do more to keep the current state of the field down than to level it up. Their objection to our overall conclusions boils down to a criticism of consistency of data collection. As such, the fact that the authors overlooked substantial parts of the supplementary information in our Article might explain why we radically disagree on certain conclusions.

In addition to the tendentious language found in their Comment, Tian et al. also make several imprecise statements. It is claimed, for instance, that our Article argued Transeurasian dispersals were "driven by Neolithic farmers in the West Liao River region", whereas our conclusion was that Transeurasian farmers expanded *from* the West Liao basin. The Commentary suggests there is insufficient evidence that Transeurasian dispersals "can be correlated with the spread of Early Neolithic millet farmers across Northeast Asia" but our conclusion was that, while primary dispersals began in the Early Neolithic, further expansions across the region occurred over the Late Neolithic and Bronze Age. The dispersals were a historical *process* not an event.

#### 4.2 Triangulation

With regard to the methodology of "Triangulation" applied in our Article, the authors limit themselves to the statement that "Robbeet's et al. … neither define nor describe the method of "triangulation" and that "[i]t is thus impossible to follow their method and combine the three corrected analyses." However, the concept of "Triangulation" is well described in the methodology section and illustrated in the Supplementary Information (SI 26) of our Article. Triangulation is the strategic use of multiple disciplines, methods, datasets and researchers to address one question in order to enhance the validity and credibility of research findings. Over the last decades, triangulation has been applied in many research fields, including sociology[78], criminology[79], medicine[80,81], mathematics[82,83,84] and anthropology[85]. In combining biology, language and culture, our Article draws on the holistic tradition of anthropology and cites Kirch and Green's monograph[85] which provides extensive historiographic, theoretical and empirical support for the triangulation method. The importance of triangulating evidence from historical linguistics, archaeology and biological

---

78 Campbell, D. T. & Fiske, D. W. Convergent and discriminant validation by the multitrait-multimethod matrix. *Psychological Bulletin*, **56** (2), 81-105 (1959)

79 Saks, M. News and views. Criminology. Methodological Triangulation. Nature Human Behaviour **2**, 806-807 (2018).

80 Lawlor, D. et al.. Triangulation in aetiological epidemiology. International Journal of Epidemiology (2016), 1866–1886 .

81 Munafò, M. R. & Smith, G. D. Comment. Repeating experiments is not enough. Nature **553**, 399-401 (2018)

82 Mathison S. Why triangulate? Educ Res **17**, 13–17 (1988).

83 Flick U. Triangulation revisited: strategy of validation or alternative? J Theory Soc Behav **22**, 175–97 (1992).

84 Tegmark M. Our mathematical universe: My quest for the ultimate nature of Rreality. (Penguin Books, 2014).

85 Kirch, P.V. & Green, R.C. *Hawaiki, ancestral Polynesia: An essay in historical anthropology* (Cambridge Univ. Press, 2001).

anthropology has been stressed by numerous previous scholars, including several authors of our Article and one member of Tian et al.'s team[51,86,8788,89,90,91,92]. According to Kirch and Green, the triangulation method aims at a historical reconstruction located within a "triangle or polygon of error" (p. 42). In our Article, a range of both qualitative and quantitative methods were used to reduce the "polygon of error" for the history of Transeurasian. Triangulation is not a quantitative method that can be said to "fail"; rather, it employs parsimony in evaluating all the available scientific evidence in the most comprehensive way. Tian et al.'s apparent inability to understand the method generates numerous problems, especially in their genetics and archaeology sections.

#### 4.3. Replication

Tian et al. insist that "[their] attempts to replicate Robbeets et al.'s results show significant discrepancies on all three fronts". However, on a closer look, there is no real attempt at replication of our research but rather an application of self-designed approaches to self- (and even mis-) scored data. On the linguistic front, neither our historical-comparative approach nor our Bayesian analysis is replicated. The authors prefer to criticize a small fraction (ca. 10%) of our core evidence and to apply questionable automated cognate detection methods to a limited selection of our cognates. On the archaeological front, Tian et al. replicate a single Bayesian test on the basis of erroneous re-scorings, ignoring our qualitative analyses which provide key support for the overall conclusions. On the genetics front, there is no attempt at re-running our analysis, just an additional calculation of f-statistics, which we have performed more carefully above. As such, the authors' claim of having replicated our research is unfounded to begin with.

---

[86] Hudson, M. *Ruins of Identity: Ethnogenesis in the Japanese Islands* (Univ. Hawai'i Press, 1999).

In addition, it can be noted that the Commentary puts too much weight on the recapitulation of a single Bayesian test. Repeating one selected approach is not enough. Reliably repeatable results will not always yield robust conclusions. Repetition could actually make things worse. Consistent findings could take the status of confirmed truths when they reflect failings in the study design, the methods or analytical tools.[81] We believe that an essential protection against such flaws is triangulation. Each dataset, method and discipline has its own strengths and weaknesses but results that agree across different datasets, methodologies and disciplines are less likely to be artefacts. Verifying results requires disparate lines of evidence. That is why triangulation is much more telling than mere replication in this case.

The commentary concludes with a *contradictio in terminis*, notably that Tian et al. "*respectfully* suggest the paper be withdrawn". From the point that Tian et al. start to use language such as "the authors bend the evidence" and "we suggest withdrawal", our feelings about the discussion changed from colleagues having a respectful scientific disagreement to an all-or-nothing battle where any admission of weakness could provide ammunition for their retraction campaign. We believe that Tian et al.'s attitude lacks professionalism and is counterproductive to progress in the field.

[93] Vovin, A. *Koreo-Japonica: A re-evaluation of a common genetic origin* (University of Hawai'i Press, 2010).
[94] Erdal, M. *Old Turkic word formation. A functional approach to the lexicon* (Harrassowitz, 1991).
[95] Martin, S. E. *A reference grammar of Korean* (Tuttle, 1992).
