## Supplementary Info 2: Data and Methods for "Triangulation reduces the polygon of error for the history of Transeurasian"

**SI 2 Data and Methods**

**1. Data**

**1.1. Linguistics**

**1.1.1. Basic vocabulary**

We revised a comparative dataset representing 254 basic vocabulary concepts for 98 Transeurasian languages, including contemporary and historical varieties. These concepts are based on a merger of the Leipzig-Jakarta 200 list ^[1]^ and the Jena 200 list.^[2]^ The inventory of basic vocabulary datasets is given in SI 1: Table 1. The identification of cognate sets is supported sound correspondences across the Transeurasian languages presented in SI 1: Tables 3 and 4.

**1.1.2 Cultural vocabulary**

We revised a dataset of cultural vocabulary relating to subsistence including 43 cognate sets. The inventory of comparative sets relating to subsistence aimed at the reconstruction of the cultural environment of the speakers of Proto-Transeurasian is given in SI 1: Table 2. The identification of cognate sets is supported sound correspondences across the Transeurasian languages presented in SI 1: Tables 3 and 4.

**1.2. Archaeology**

We revised a rescoring of our dataset of 171 cultural traits for a total of 255 Neolithic-Bronze Age archaeological sites/phases from the West Liao River basin (33), the Amur (Jilin, Heilongjiang and inland Liaoning) (32), the Primorye (4), the Liaodong peninsula (37), the eastern steppes (1), the Shandong peninsula (4), the Yellow River basin (2), the Korean peninsula (57) and the Japanese Islands (85). The sites date from 8400-1700 BP and include the Early Neolithic to Bronze Age in northeast China, the Middle Neolithic Zaisanovka culture in the Primorye, the Middle-Late Neolithic Chulmun and Bronze Age Mumun cultures in Korea, and the Late Neolithic/Bronze Age Final Jomon and Yayoi cultures in western Japan. Categories of cultural traits scored comprised ceramics (69), stone tools (37), buildings and houses (9), plant and animal remains (26), shell and bone artefacts (17), and burials (13). We found numerous inconsistencies in the rescored dataset by Tian et al. (2022). Out of the 109 sites which were re-scored, at least 61 are problematic (see SI 1: Section 2.1).

**1.3. Genetics**

In order to quantify the lack of resolution to distinguish the precise source of East Asian ancestry in our data more precisely, we extended the symmetry test performed by Tian et al. to a wider range of populations. To this end we compared Upper Xiajiadian to a variety of other ancient populations in the Amur River, West Liao River and Yellow River Basins, including the Early Neolithic Jalainur in the Amur River Basin (SI 1: Fig. 1), the Middle Neolithic Hongshan (SI 1: Fig. 2), the Late Neolithic Lower Xiajiadian in the West Liao River Basin (SI 1: Fig. 3), the Middle Neolithic Yangshao in the Yellow River Basin (SI1: Fig. 4), the Late Neolithic Longshan in the Yellow River Basin (SI 1: Fig. 5) and Late Bronze/Iron Age individuals in the Yellow River Basin (SI 1: Fig. 6).

**2. Methods**

**2.1. Linguistics**

We applied the standard historical-comparative method, which is the traditional method ~~a~~ for establishing a genealogical relationship between two or more languages^[3]^,^[4]^,^[5]^,^[6]^,^[7]^. It is a procedure for inferring an unattested ancestral state of a language on the evidence of data that are available from a later period. The systematicity by which the sounds of a language change over time allows linguists to reconstruct the sounds and words of the proto-language. In line with the historical-comparative method, the identification of cognate sets in our linguistic dataset is supported sound correspondences across the Transeurasian languages presented in SI 1: Tables 3 and 4. We applied this method to the datasets within the sphere of the basic vocabulary (see Section 1.1.1) as well as the cultural vocabulary (See Section 1.1.2).

In addition, we analyzed the distribution of cognates across cross-linguistic datasets, based on comparison of the Transeurasian cognate set distributions with those of long-established language families, such as Sino-Tibetan and Indo-European. To this end, we considered variables such as number of surviving daughter languages, estimated time depth of the root of the family and the age of the earliest written records (see SI 1: Table 11).

**2.2. Archaeology**

**2.2.1 Qualitative analysis**

The database was used to analyse changes in the distribution of Neolithic and Bronze Age artefacts over time, especially in relation to the spread of agricultural systems in Northeast Asia. This led to the identification of four stages of Neolithic/Bronze Age farming dispersals in Northeast Asia based on archaeobotanical and zooarchaeological remains of domesticated plants and animals (see SI 1: Table 12).

**2.2.2 Archaeological phylogeny**

The cultural data in our archaeological database were analyzed using Bayesian phylogenetic methods.With respect to this analysis, we produced an updated XML file, correcting the number of taxa in our Article, under <https://github.com/rbouckaert/Eurasia3angle/releases/download/v1.0/culturexml.zip>

Even if the inclusion of the XML file does not reflect any major differences in the original results, it facilitates their replication.

**2.3. Genetics**

We compared Upper Xiajiadian to a variety of other other ancient populations in the Amur River, West Liao River and Yellow River Basins, as specified in Section 1.3. We calculated 𝑓4-statistics in the form of 𝑓4 (Mbuti, X; Upper_Xiajiadian) using a list of ancient and modern populations published in our Article as X by qpDstat v755 implemented in ADMIXTOOLS^[8],^^[9]^ with the parameters *f*4-mode set to YES and printsd set to YES.

1. Haspelmath, M. & Tadmor, U. *Loanwords in the World’s Languages: A Comparative Handbook*. (Mouton de Gruyter, 2009).
2. Savelyev, A. & Robbeets, M. Bayesian phylolinguistics infers the internal structure and the time-depth of the Turkic language family. *J. Lang. Evol.* 1-15 (2019) doi: 10.1093/jole/lzz010
3. Campbell, L. & Poser,W. J.. Language classification: History and method. (Cambridge Univ. Press, 2008)
4. Hock, H. H. & Joseph, B. D. Language history, language change, and language

   relationship: An introduction to historical and comparative linguistics. 2d ed. (Mouton

   de Gruyter, 2009)
5. Crowley, T. & Bowern, C.. An introduction to historical linguistics. 4th ed. (Oxford Univ. Press, 2010)
6. Bowern, C. & Evans, B. (eds.) The Routledge handbook of historical linguistics. (Routledge, 2015).
7. Robbeets, M. Comparative reconstruction in linguistics. *Oxford Bibliographies/ Linguistics* (Oxford Univ. Press, 2018). <http://www.oxfordbibliographies.com/abstract/document/obo-9780199772810/obo-9780199772810 0215.xml>
8. Peter, Benjamin M. Admixture, population structure, and F -statistics. Genetics 202(4). 1485–1501. https //doi.org/10.1534/genetics.115.1 3913. (2016)
9. Patterson, Nick, Priya Moorjani, Yontao Luo, Swapan Mallick, Nadin Rohland, Yiping Zhan, Teri

   Genschoreck, Teresa Webster & David Reich. Ancient admixture in human history. Genetics

   192(3). 1065–1093 (2012). https//doi.org/10.1534/genetics.112.145037.
